# Characterization of the secretome, transcriptome and proteome of human β cell line EndoC-βH1

**DOI:** 10.1101/2021.09.09.459582

**Authors:** Maria Ryaboshapkina, Kevin Saitoski, Ghaith M. Hamza, Andrew F. Jarnuczak, Claire Berthault, Kaushik Sengupta, Christina Rye Underwood, Shalini Andersson, Raphael Scharfmann

## Abstract

Early diabetes research is hampered by limited availability, variable quality and instability of human pancreatic islets in culture. Little is known about the human β cell secretome, and recent studies question translatability of rodent β cell secretory profiles. Here, we verify representativeness of EndoC-βH1, one of the most widely used human β cell lines, as a translational human β cell model based on omics and characterize the EndoC-βH1 secretome. We profiled EndoC-βH1 cells using RNA-seq, Data Independent Acquisition (DIA) and Tandem Mass Tag proteomics of cell lysate. Omics profiles of EndoC-βH1 cells were compared to human β cells and insulinomas. Secretome composition was assessed by DIA proteomics. Agreement between EndoC-βH1 cells and primary adult human β cells was ~90% for global omics profiles as well as for β cell markers, transcription factors and enzymes. Discrepancies in expression were due to elevated proliferation rate of EndoC-βH1 cells compared to adult β cells. Consistently, similarity was slightly higher with benign non-metastatic insulinomas. EndoC-βH1 secreted 671 proteins in untreated baseline state and 3,278 proteins when stressed with non-targeting control siRNA, including known β cell hormones INS, IAPP, and IGF2. Further, EndoC-βH1 secreted proteins known to generate bioactive peptides such as granins and enzymes required for production of bioactive peptides. Unexpectedly, exosomes appeared to be a major mode of secretion in EndoC-βH1 cells. We believe that secretion of exosomes and bioactive peptides warrant further investigation with specialized proteomics workflows in future studies.

**Graphical abstract:** 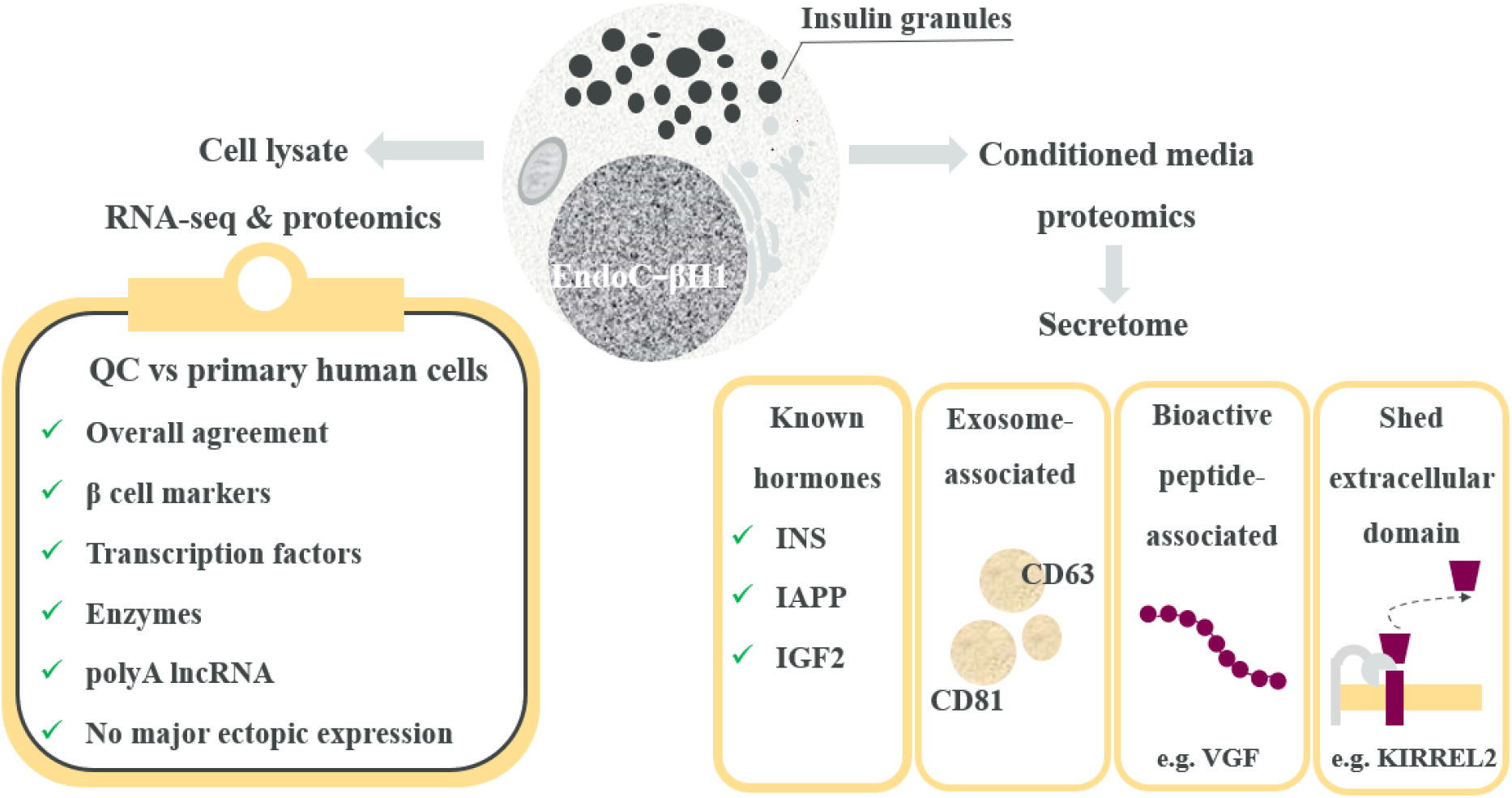

**Highlights:**

- We validate EndoC-βH1 as a translational human β cell model using omics.
- We present the first unbiased proteomics composition of human β cell line secretome.
- The secretome of human β cells is more extensive than previously thought.
- Untreated cells secreted 671 proteins and stressed cells secreted 3,278 proteins.
- Secretion of exosomes and bioactive peptides constitute directions of future research.

## INTRODUCTION

Close to half a billion people are living with diabetes worldwide and this number is expected to increase by 25% in 2030 (1). Despite diabetes being a global health burden, the mechanisms underlying normal β cell function and failure are not yet fully elucidated. This is partly due to limited availability of high quality human pancreatic islets for research purposes combined with variable quality and variation in relative cell type composition of islet preparations (2, 3). Moreover, primary human islets rapidly loose functional expression pattern in culture (4). To overcome these limitations, pancreatic islets from rodents have been used to study β cell biology for decades leading to significant advances in our understanding of the disrupted pathways causing impaired β cell function. However, the cellular architecture and secretory profile of rodent islets is substantially different from that of human islets (5, 6), manifesting the need for human β cell lines, which mirror the expression and secretory pattern of primary human β cells.

Several human β cell lines are currently available - 1.1B4, 1.4E7 and 1.1E7 (7), T6PNE (8), EndoC-βH1 (9), EndoC-βH2 (10), EndoC-βH3 (11) and ECN90 (12). The corresponding full text searches for these cells lines in Google Scholar retrieved 774 (22.5%), 993 (28.8%), 947 (27.5%), 13 (0.4%), 598 (17.4%), 67 (1.9%), 41(1.2%) and 11(0.3%) entries as of March 2021, respectively. Thus, EndoC-βH1 cells remain one of the most commonly used human β cell lines despite availability of next-generation cell lines EndoC-βH2 (10) and EndoC-βH3 (11) with inducible growth-arrest. Taken together, these three generations of EndoC-βH cells are used by over 150 research labs world-wide (https://www.humanbetacelllines.com/)(13).

Validity of EndoC-βH1 cells as a human β cell model has been previously established based on functional experiments such as glucose-stimulated insulin secretion (14–16) and marker expression profiling by qPCR (9, 16). Recently, EndoC-βH1 open chromatin, transcriptomics and miRNA landscapes were reported to be most similar to adult human β cells or islets compared to α cells, adipocytes, muscle and peripheral blood cells (17). Two proteomics studies investigated the response to interferons in EndoC-βH1 cells (18, 19), but did not validate EndoC-βH1 as a β cell model. It remains unclear what are the discrepancies of EndoC-βH1 with adult β cells and limitations (scope / applicability) of the model.

The aim of this study is to validate EndoC-βH1 as an *in vitro* human β cell model based on omics. We evaluate the EndoC-βH1 cell transcriptome and proteome against corresponding omics profiles of primary β cells and insulinoma samples. Due to their proliferative nature, EndoC-βH1 are more similar to human primary insulinomas than to adult human β cells which have an extremely low proliferation rate. We show that EndoC-βH1 cells belong to the β cell lineage and recapitulate features of adult primary β cells well. We observe that EndoC-βH1 exhibit ~90% similarity with reference adult human β cell transcriptome and proteome. Similarity exceeding 90% was noted for β cell markers, transcription factors and enzymes. We also extensively studied negative control markers of other cell and tissue lineages and confirm lack of major expression abnormalities.

Further, we address a major knowledge gap: β cells are a crucial endocrine cell type, yet little is known about the β cell secretome, especially in humans. Few studies reported secretome composition of rodent insulinoma cell lines (20–22), while the secretome of human immune-cell depleted islet tissue has been profiled using the SOMAscan 1,300 proteomics assay only this year (23). The SOMAscan study suggests that the human and murine secretomes may be quite distinct (23), making it unclear to what extent findings of secretome studies in murine β cells can be extrapolated to human. In this study, we present the first characterization of human β cell line EndoC-βH1 secretome by Data Independent Acquisition (DIA) proteomics. This novel data indicates that the secretome of β cells may be more extensive than previously thought, and highlights surprising abundance of exosome-associated proteins.

## EXPERIMENTAL PROCEDURES

### Experimental Design and Statistical Rationale

The experiment series was designed to investigate the effects of PCSK9 knockdown on β cell transcriptome, proteome and secretome. Non-targeting control siRNA samples were used to evaluate EndoC-βH1 as a translational β cell model and to characterize composition of the secretome, and are presented in this manuscript. Effects of PCSK9 knockdown are further explored in [**reference**]. For all experiments, the PCSK9 knockdown efficiency was analyzed at the mRNA level by RT-qPCR and at the protein level by Western blot in both cell extracts and conditioned media prior to conducting omics data acquisition. Pilot Tandem Mass Tag (TMT) proteomics cell lysate time course (PXD027898) contained untreated baseline and siRNA control samples and was used to verify lack of major impact of control siRNA treatment on proteome composition. Five other experiments included non-targeting control siRNA and siPCSK9 samples, had confirmed PCSK9 knockdown and supported both parts of the study. EndoC-βH1 cells treated with control siRNA and PCSK9 siRNA at 72h were profiled by RNA-seq (GSE182016), cell lysate DIA proteomics (PXD027911), cell lysate TMT proteomics (PXD027921) and secretome DIA proteomics (PXD027913). In addition, a time-course profiling of the secretome of PCSK9 siRNA vs control siRNA cells at 24, 48 and 72 hours was performed with DIA proteomics (PXD027920).

#### Sample size calculation

RNA-seq, cell lysate TMT proteomics and DIA secretome time course were unpowered pilot experiments with three replicates per condition. Sample size for cell lysate and secretome DIA proteomics at 72 hours (PXD027911 and PXD027913) was calculated based on the pilot cell lysate TMT experiment (PXD027921) as 12 non-targeting control siRNA and 12 siPCSK9 samples (power 80%, alpha 0.05, N proteins expected to be detected 9,000, alpha corrected for multiple testing 5.6e-6, Cohen d for PCSK9 knockdown −2.93, distribution of normalized protein intensities assumed to be approximately normal, two-tailed t-test).

### EndoC-βH1 cell culture

EndoC-βH1 cells were cultivated in low glucose DMEM medium (5,6 mmol/l; Sigma-Aldrich) supplemented with 2% BSA fraction V (Roche Diagnostics), 50 μmol/L 2-β-mercaptoethanol (Sigma-Aldrich), 10 mmol/L nicotinamide (Calbiochem), 5.5 μg/mL transferrin (Sigma-Aldrich), 6.7 ng/mL sodium selenite (Sigma-Aldrich), 100 units/mL penicillin, and 100 μg/mL streptomycin (ThermoFisher Scientific). The cells were seeded on plates previously coated with 1.2% Matrigel containing-3 μg/ml fibronectin (both from Sigma-Aldrich) and cultured at 37°C and 5% CO2 as previously described (9).

For siRNA transfection, EndoC-βH1 cells were passaged, seeded at 10^5^ cells /cm^2^. 24 hours later, cells were transfected in OptiMEM using Lipofectamine RNAiMAX (Life Technologies, Saint Aubin, France) with siRNA SMARTpools (Horizon Discovery LTD): non targeting Control (siCTRL, D-001810-01-20) or siRNA PCSK9 (M-005989-01-0005) at the final concentration of 80 nM as described (24). Medium was replaced 2.5 hours later with fresh EndoC-βH1 culture medium.

For proteome experiments, EndoC-βH1 cells were pre-incubated for 2 hours in BSA/phenol red-free culture medium before transfection. After transfection, the cells were grown in fresh BSA/phenol red-free culture medium for 3 days. Cell pellets were collected by trypsinization and washed 3 times in PBS. Conditioned media were collected, treated with protease and phosphatase inhibitors (Roche Diagnostics) and the debris eliminated by centrifugation at 1,500 rpm at 4 °C. All the samples were stored at −80°C.

### Fluorescence-activated cell sorting

EndoC-βH1 cells were trypsinized and washed three times in PBS. Cell death was assessed by incubating the cells with annexin-V antibody (APC, #*B187891*, Biolegend) for 15 minutes at room temperature, in the dark in annexin-V buffer (#*422201*, Biolegend). Propidium iodide was added before FACS analysis. For proliferation studies, cells were fixed, and permeabilized with a transcription factor staining buffer set (#*00-5523-00*, Thermo Fisher Scientific) according to the manufacturer’s instructions. Cells were then incubated with Ki67 antibody (PerCP/Cy5.5, #*B238642*, Biolegend) for 30 minutes at room temperature in the dark in permeabilization buffer, and then with DAPI (BD Biosciences) for 10 minutes at room temperature in the dark. FACS analysis was carried out using a FACS Aria III (BD Biosciences, San Jose, CA, USA). Data were analyzed using FlowJo 10.7 software (RRID:SCR_008520).

### RNA-sequencing

RNeasy Micro Kit (Qiagen) was used to extract total RNA from EndoC-βH1 cells. Genomic DNA was removed by DNAse treatment following the RNeasy Micro Kit protocol. RNA quality (RNA integrity number) was determined by electrophoresis using the Agilent 2100 Bioanalyzer (Agilent Technologies, Palo Alto, CA, USA) as per the manufacturer’s instructions. To construct the libraries, 600 ng of high quality total RNA sample (RIN >8.5) was processed using TruSeq Stranded mRNA kit (Illumina) according to manufacturer’s instructions. Briefly, after purification of poly-A containing mRNA molecules, mRNA molecules were fragmented and reverse-transcribed using random primers. Replacement of dTTP by dUTP during the second strand synthesis enabled strand specificity. The addition of a single A base to the cDNA was followed by ligation of Illumina adapters. Then, the libraries were quantified with Qubit fluorometer DNA HS kit (Invitrogen). The library profiles were assessed using the DNA High Sensitivity LabChip kit on an Agilent Bioanalyzer. Libraries were sequenced on an Illumina Nextseq 500 instrument using 75 base-length reads V2.5 chemistry in a paired-end mode. After sequencing, a primary analysis based on AOZAN software (Ecole Normale Supérieure, Paris) was applied to demultiplex and control the quality of the raw data (based on FastQC modules / version 0.11.9). Raw reads were aligned to human genome GRCh38 release 96 using STAR version 2.7.1a (25). Transcript quantification was performed with RSEM (26). Quality control metrics were generated with Picard [“Picard Toolkit.” 2019. Broad Institute, GitHub Repository. http://broadinstitute.github.io/picard/; Broad Institute] and STAR (25) and examined visually in addition to clustering and PCA. All samples passed quality control. TPM (transcript per million) normalization was applied for descriptive analysis in this study.

### Cell Proteome Sample Preparation

Cell pellets were thawed on ice and subjected to sample preparation with the PreOmics iST NHS or iST kit (PreOmics, Planegg, Germany) according to manufacturer's protocols. Briefly, iST or iST-NHS lysis buffer was added directly to the cell pellet and incubated at 95°C for 10 min for cell lysis, reduction and alkylation of proteins. Cell lysate was normalized using BCA assay (Thermo Fisher Scientific) and 50 μg from each condition was subjected to enzymatic cleavage for 3 hours by adding equal amounts of endoproteinase Lys-C and trypsin in a 1:50 (wt/wt) enzyme:protein ratio. For experiments utilizing Tandem Mass Tags™ (TMT™), TMT™ reagents were reconstituted in anhydrous acetonitrile (ACN) and added to peptides at a 4:1 reagent:peptide. De-salting and purification were performed according to the PreOmics iST or PreOmics iST-NHS protocol on a styrene divinylbenzene reversed-phase sulfonate (SDB-RPS) sorbent. IST Purified peptides were vacuum-centrifuged to dryness and reconstituted in double-distilled water with 2 vol% ACN and 0.1 vol% formic acid (FA) for single-run LC-MS analysis. IST-NHS TMT™ labeled peptides were reconstituted in pH 10.0, 20mM ammonia hydroxide and combined evenly to create a single sample.

### HPRP Fractionation

TMT™ labeled peptides were fractionated into 96 fractions across a 96 well plate, using high pH reverse phase chromatography on an Agilent 1100 system and Phenomenex Gemini 5μm C18 250mm × 2mm column. The linear gradient profile consisted of 2% to 90% mobile phase B in 70 minutes at flow rate of 0.20mL/min. Mobile phase A consisted of pH 10.0, 20mM ammonia hydroxide in water and mobile phase B consisted of 20mM ammonia hydroxide in ACN. The 96 fractions were combined within each column orthogonally into a total of 12 concatenated analytical samples. Samples were vacuum-centrifuged to dryness and were then reconstituted with double-distilled water with 2 vol% ACN and 0.1 vol% FA for LC-MS analysis.

### Secretome Sample Preparation

Cell conditioned media was thawed on ice and was concentrated down to ~100 μL using Amicon® Ultra-15 3k MWCO centrifugal filter units (Millipore Sigma). Buffer exchange into PreOmics iST lysis buffer was performed, followed by incubation at 95°C for 10 min for reduction and alkylation of proteins. Protein amounts were normalized using BCA assay (Thermo Fisher Scientific) and equal amounts from each condition was subjected to enzymatic cleavage for 3 hours by adding equal amounts of endoproteinase Lys-C and trypsin in a 1:50 (wt/wt) enzyme:protein ratio. De-salting and purification were performed according to the PreOmics iST protocol on a SDB-RPS sorbent. Purified peptides were vacuum-centrifuged to dryness and reconstituted in double-distilled water with 2 vol% acetonitrile (ACN) and 0.1 vol% formic acid (FA) for single-run LC-MS analysis.

### LC-MS/MS Measurement

#### Orbitrap Exploris 480 Mass Spectrometer (PXD027911 and PXD027920)

Peptides were loaded onto a 25cm IonOpticks Aurora Series emitter Column (25cm × 75u ID, 1.6um C18; IonOpticks, Australia) performed by a Dionex Ultimate 3000 coupled online to an Exploris 480 Mass Spectrometer equipped with a Nanospray Flex Ion Source, integrated with a column oven (PRSO-V1, Sonation, Biberach, Germany) maintained at 50°C. Peptides were separated using a non-linear gradient. Mobile Phase A was 0.1 vol% FA, 3 vol% ACN in water, while Mobile Phase B was 90 vol% ACN, 0.1 vol % FA. The gradient was operated at 400nl/min flowing 3 vol% B for 25min, 5-17 vol% B over 72min, 17-24 vol % B over 18min, 24-30 vol % B over 10min, 30-85 vol % B over 3min, hold at 85 vol% B for 7min, 85-3 vol% B over 0.1min, and hold at 3 vol% B for 15min. Orbitrap Exploris 480 was operated in BoxCar Data Independent Acquisition (DIA) positive mode where spray voltage was set to 1600V, funnel RF at 40%, and heated capillary temperature at 275°C. Method Timeline experiment consisted of 1 MS1 Scan, 1 tSIM scan and 1 tMS2 scan. MS1 scan was operated at 120k resolution, 400-1200 m/z scan range, 40% RF lens, 300% AGC target, and 54ms IT. MS1 tSIM was operated with Multiplexed Ions enabled (12 ions), at 120k resolution, 300% AGC target, 20ms IT with a set loop control of 2N (number of spectra). BoxCar windows spanned 400-1200m/z space. A total of 48 MS2 DIA variable windows were operated at 15k resolution with normalized collision energy of 28%, 1000% AGC target with 22ms IT spanning 400-1200m/z range taken into account heavily dense peptide regions.

#### Orbitrap Fusion Lumos Mass Spectrometer (PXD027913)

Peptides were loaded onto a ReproSil-Pur 120 C18AQ 1.9um in-house packed to a 5um tip 75u ID × 360u × 50 cm column using a Thermo EASY-nLC1200 connected through a Nanospray Flex Ion Source, integrated with a Column Oven (PRSO-V1, Sonation, Biberach, Germany) maintained at 50°C. Mobile Phase A was 0.1 vol% FA, 3 vol% ACN in water, while Mobile Phase B was 90 vol% ACN, 0.1 vol% FA. The gradient which had a flow of 300nl/min consisted of a non-linear ramp of 3-24 vol% B in 85min, 24-30 vol% B for 15min, 30-95 vol% B in 5 min, 95 vol% B hold for 5min, and re-equilibration to 3 vol% B for 10min. The LC was connected to an Orbitrap Fusion Lumos Mass Spectrometer operated in DIA Positive mode. Briefly, spray voltage was set to 2500V, Ion transfer tube set to 300°C, MS1 resolution at 120k, MS2 resolution at 30k, scan range of 350-1650 m/z for MS1 and 350-1650 m/z for MS2. RF lens was set to 30%, MS1 IT to 20ms, MS2 IT to 60ms. MS1 AGC target set to 3e6, MS2 AGC target set to 1.5e6 with 28% HCD collision energy. DIA variable windows covered 400-1650 m/z space taken into account heavily dense peptide regions.

#### Orbitrap Fusion Lumos Mass Spectrometer (PXD027898 and PXD027921)

Peptides were loaded onto a ReproSil-Pur 120 C18AQ 1.9um in-house packed to a 5um tip 75u ID × 360u × 50 cm bed volume column using a Thermo EASY-nLC1200 connected through a Nanospray Flex Ion Source, integrated with a Column Oven (PRSO-V1, Sonation, Biberach, Germany) maintained at 50°C. Mobile Phase A was 0.1 vol% FA, 3 vol% ACN in water, while Mobile Phase B was 90 vol% ACN, 0.1 vol% FA. The gradient which had a flow of 250nl/min consisted of non-linear ramp of 3-5 vol% B in 5min, 5-20 vol% B for 80min, 20-32 vol% B in 30 min, 32-95 vol% B in 1min, and hold at 95 vol% B for 14 min. The LC was connected to an Orbitrap Fusion Lumos Mass Spectrometer operated in Data Dependent Acquisition Positive mode. Briefly, spray voltage was set to 2200V, Ion transfer tube set to 275°C, MS1 resolution at 60k, MS2 resolution at 15k, scan range was set to 375-2000 m/z for MS1 and first mass of 100 m/z for MS2. MS1 IT to 50ms, MS2 IT to 22ms. MS1 AGC target set to 1e6, MS2 AGC target set to 1e6 with 38% HCD collision energy. Quadrupole isolation window was set to 1m/z with 30 second exclusion duration. MIPS filter was set to peptide, filter intensity threshold was activated with Maximum intensity set to 1E20, minimum intensity set to 4E4, intensity filter type set to intensity threshold.

### Mass Spectrometry Analysis

DIA experiments were analyzed using Spectronaut™ V14.10 (Biognosys AG, Switzerland) direct DIA analysis using a Swiss-Prot human canonical database downloaded on 2/28/2020; the database contained 20,350 entries and utilized Pulsar as the search engine. Analysis settings were maintained to factory settings where identification was set to 1% FDR for precursor and protein level and quantification was conducted on MS2 level for specific digest type of Trypsin/P. Static modifications of Carbamidomethyl (+57.021 Da) and dynamic modification N-terminal acetylation (+42.011 Da).

TMT experiments were analyzed using Thermo Proteome Discoverer 2.4. Briefly, the raw files were searched through Sequest HT using canonical database as above with the following parameters: Trypsin (full), max missed cleavage of 2, minimum peptide length of 6, max peptide length of 144, precursor mass tolerance of 10ppm, fragment mass tolerance of 0.02Da, dynamic modification: oxidation (+15.995 Da) of methionine, N-terminal acetylation (+42.011 Da), static modification: TMT (+229.163 Da) on peptide N-terminus and lysine, PreOmics cystine modification (+113.084 Da). Target/decoy was performed using Percolator: concatenated, q-value, with FDR on peptide and protein level of 1%. TMT reporter ion quantification was carried out using 50% co-isolation threshold, and average reporter S/N threshold of 10.

### Down-stream analysis

TMT proteomics data were normalized using scaling factor normalization and log2 transformed. DIA proteomics data were normalized using quantile normalization as implemented in limma (27) and log2-transformed. DIA proteomics data contained small amount of missing values: 0.7 (0.2-1.8)% missing protein intensity values per sample in cell lysate PXD027911 and 1.5 (0.9-2.3)% in secretome PXD027913 experiments. Missing values were imputed using sequential imputation method (28, 29). Approximately normal distribution of normalized protein intensities were confirmed by visual examination of quantile-quantile plots. Ambiguously annotated proteins (e,g., tryptic peptides aligning to several immunoglobulin kappa chain proteins) were excluded prior to statistical analysis. Differential protein expression analysis of siRNA control vs untreated cells was performed with mixed-effect linear model with condition as fixed effect term and TMT plex as random intercept. Functional classification of the proteins (30) was evaluated using AmiGO2 (31) and PANTHER (32). Proteins may have more than one functional annotation (e.g., be involved in several pathways), therefore the sum of percentages of proteins with different annotations may exceed 100%. Figures were made in R version 4.0.2(33) with ggplot2 (34) and ggVennDiagram (35).

## RESULTS

### Quality control of the biological material

One of the biggest challenges in secretome studies is determining whether the proteins are truly secreted by viable cells or are observed as a result of cell death or leakage. Therefore, we verified that treatment with non-targeting control siRNA did not drastically affect cell viability, cell type identity or functional state. We confirmed that treatment with control siRNA did not affect cell proliferation (**Fig 1A**) or viability (**Fig 1B** and **C)**, which are important for secretome assessment as these factors could affect membrane permeability. We assessed the differences in protein expression between untreated baseline and control non-targeting siRNA treated EndoC-βH1 cells. Treatment with control siRNA altered expression of 3.3% of proteins. The cells treated with control siRNA displayed signs of altered lipid metabolism and stress. However, the proteins quantified at baseline were a complete subset of the proteins quantified at 72h after transfection with control siRNA (**Fig 1D**). Thus, treatment with control siRNA did not change the repertoire of proteins expressed by the cells and cell type identity.

**Fig. 1.**
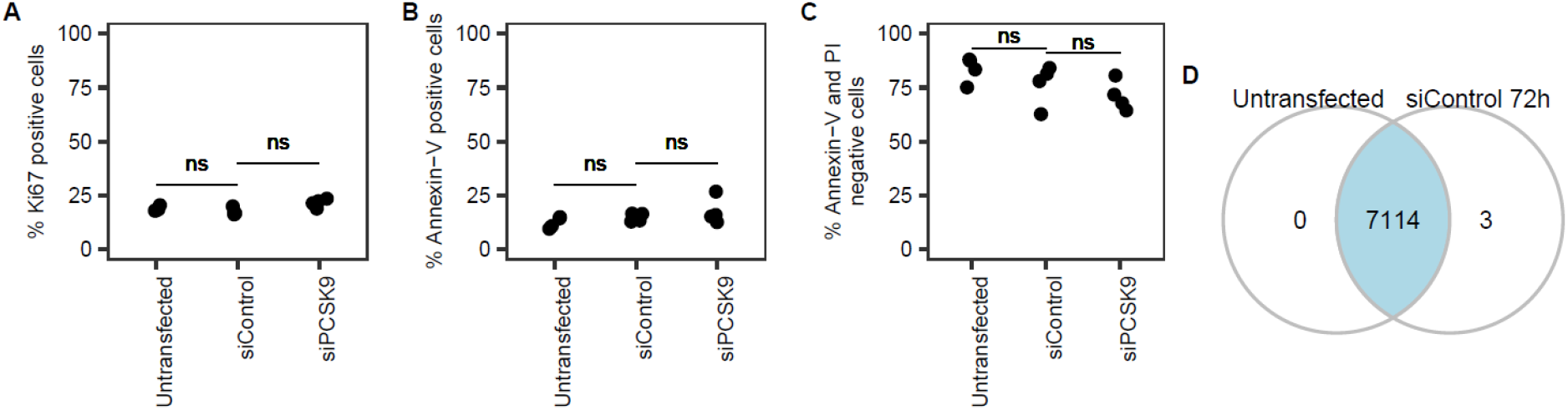
Non-targeting control siRNA did not affect EndoC-βH1 cells proliferation, viability and composition of the proteome. Ki67/Dapi staining for cell proliferation (**A**) and Annexin-V/PI staining for apoptosis (**B**) and live cells (**C**) assessed by FACS in untransfected cells compared to 72 hours post transfection with control siRNA or siPCSK9. (**D**) Proteome composition of untransfected cells and cells transfected with control siRNA at 72 hours in pilot TMT cell lysate experiment. Expression cut-off for proteins: quantified in 3 samples and unambiguously annotated.

### Validation of EndoC-βH1 as a translational human β cell model

We compared the transcriptome and proteome of EndoC-βH1 both globally and across selected positive control and negative control marker categories to published human primary β cells and insulinomas. The reference human β cell preparations were selected based on purity of the tissue: RNA-seq of adult primary human β cells from 7 non-diabetic donors, isolated by FACS sorting with at least 97% β cell purity in GSE67543 (36) and proteomics of 4 non-diabetic human islet cultures on nanostructured zirconia with 80% ± 10% purity of β cells in PXD007569 (37). The reference insulinoma samples were selected from tumors with typical presentation and without metastases: RNA-seq samples WT_MK27 and WT_MK8 in GSE118014 (38) and proteomics by Song and colleagues (39). The reference transcriptome and proteome contained less molecular species than the EndoC-βH1 transcriptome and proteome acquired in this study. Hence, we calculated percentage of the reference transcriptome/proteome that was recapitulated by EndoC-βH1 cells.

First, we compared EndoC-βH1 cells to primary human adult β cells. Overall, transcriptome and proteome of EndoC-βH1 cells resembled adult human β cells (**Fig 2A** and **2B**). EndoC-βH1 recapitulated 90.7% of β cell protein-coding transcriptome and 87.7% of the β cell - enriched islet proteome. The reference human β cell proteomics data originated from an older study and contained limited number of protein species. Therefore, the agreement between marker gene sets was calculated based on transcriptomics. Good agreement between 69% and 91% vs adult β cells was observed across individual gene/protein categories such as β cell markers, genes associated with diabetes, transcription factors, enzymes, ion channels and lncRNA (**Table S1, Fig S1-S6, Fig S8).** The lowest agreement was obtained for G-protein coupled receptors (GPCRs) with 52% of β cell GPCRs detected on mRNA level and only a few species confirmed by proteomics (**Fig S7**). In particular, median GLP1R expression in control siRNA treated EndoC-βH1 cells was 0.8 TPM whereas median expression in primary adult β cells was 61.7 TPM. We also verified low GLP1R expression with median 5 TPM in an external experiment with untransfected EndoC-βH1 cells sequenced at 200 million reads per sample in GSE133218 (18).

**Fig. 2.**
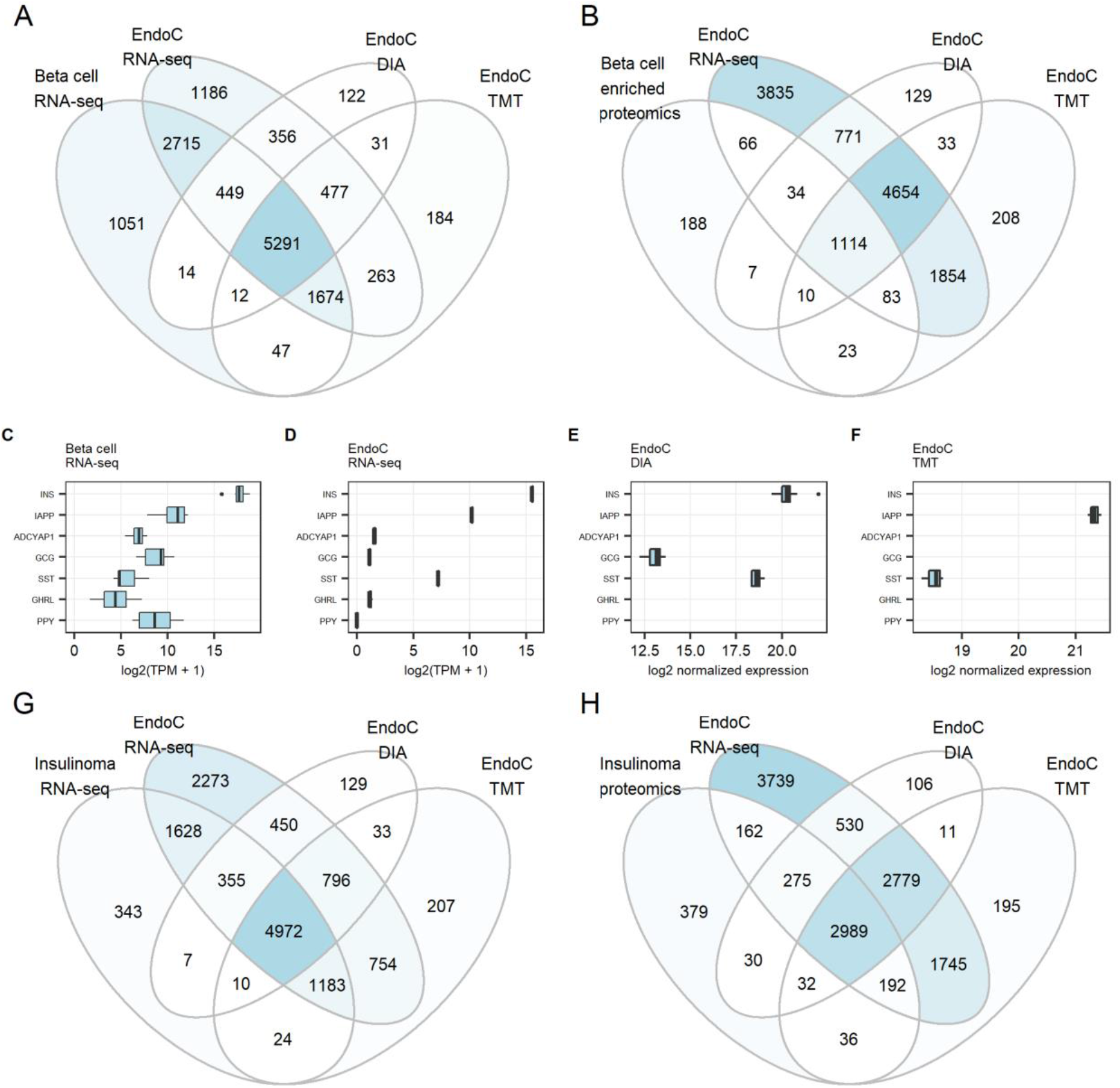
Transcriptome and proteome of EndoC-βH1 cells compared to adult human β cells and insulinomas. (**A**) Venn diagram showing the number of quantified protein-coding genes and proteins in EndoC-βH1 cells vs adult non-diabetic β cells (GSE67543). (**B**) Comparison to human β cell-enriched islet proteomics (PXD007569). (**C-F**) Hormone expression. (**G**) Comparison to protein-coding genes with expression > 1 TPM in human low-grade insulinomas without lymph node and distant metastases (GSE118014). (**H**) Comparison to proteome of human insulinomas with typical tumor presentation and without lymph node and distant metastases (39). EndoC-βH1 cells were treated with control non-targeting siRNA and harvested at 72 h. Expression cut-off for genes: median expression >= 1 TPM. Expression cut-off for proteins: quantified in >= 3 samples and unambiguously annotated. Sample sizes in EndoC-βH1 experiments: RNA-seq N = 3, TMT proteomics = 3, cell lysate DIA proteomics = 12 replicates.

Next, we investigated possible ectopic expression patterns. EndoC-βH1 expressed β cell hormones INS and IAPP at gene and protein levels and ADCYAP1 (also known as PACAP) at mRNA level. (**Fig 2C-F**). EndoC-βH1 expressed mRNA of other islet endocrine hormones (**Fig 2D**) which can also be observed in primary β cells (**Fig 2C, Fig S9**). However, detection of proglucagon and somatostatin on protein level (**Fig 2E**, **Fig 2F**) is atypical for adult human β cells. Somatostatin expression in EndoC-βH1 cells has been noted previously (16) and is observed in insulinomas (39). Proglucagon protein was detected based on a single peptide DFINWLIQTK aligning to amino acid positions 166-175 and corresponding to glucagon-like peptide 2. Proglucagon co-expression with insulin is a feature of insulinomas (39, 40). No ectopic expression was observed for other α cell markers (**Fig S10**). EndoC-βH1 were derived from fetal islet cells at 9 to 11 weeks of gestation (9). Interestingly, expression of endocrine progenitor markers was considerably reduced in EndoC-βH1 cells compared to human fetal pancreas at 9 weeks gestation (**Fig S11**). Expression of NEUROG3 was lost. Markers of fetal β cell at 12 to 18 weeks gestation were not expressed (**Fig S12**). We also observed no ectopic expression of markers of exocrine pancreas acinar (**Fig S13**) and ductal cells (**Fig S14**) and markers of other tissues (**Fig S15**).

Overall, the genes and proteins quantified in EndoC-βH1 cells but not in primary β cells were annotated with cell cycle, DNA repair, proliferation, ATP generation and mitochondrial function Gene Ontology (GO) terms. This ectopic expression pattern was consistent with proliferative phenotype. EndoC-βH1 cells double in number every 174 hours (14) whereas proliferation of adult human β cells is an extremely rare event (41). Accordingly, the EndoC-βH1 transcriptome and proteome were more similar to insulinomas than to primary human β cells (**Fig 2G-H**). EndoC-βH1 cells recapitulated 96% of the reference insulinoma transcriptome and 90.7% of the reference insulinoma proteome. Good agreement was observed across individual protein categories (**Fig 3**).

**Fig. 3.**
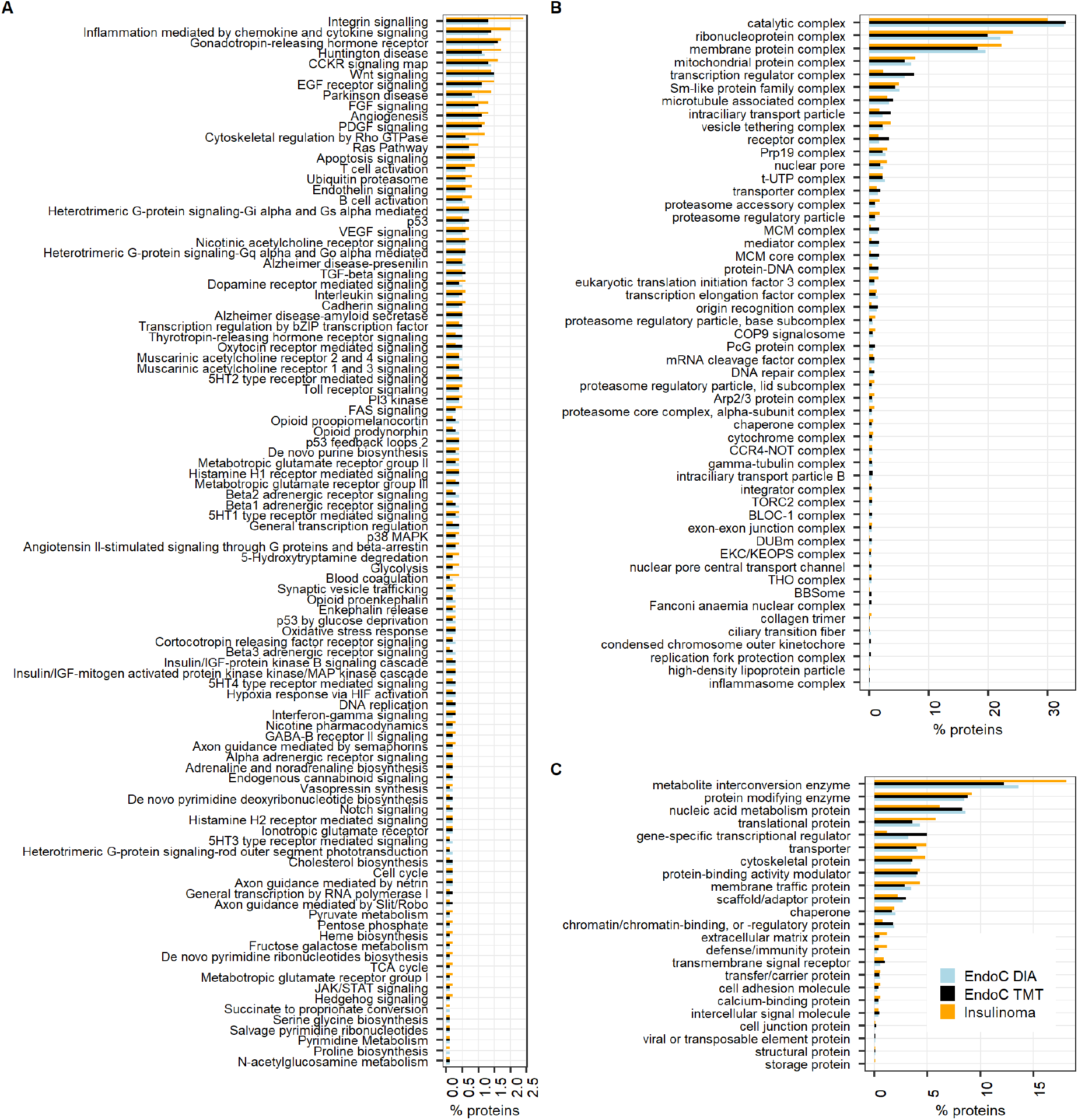
PANTHER classification of EndoC-βH1 cell proteome compared to insulinoma proteome. (**A**) Top 100 pathway by the number of annotated proteins. (**B**) Protein-containing complexes. (**C**) Protein classes. Percentages indicate the number of proteins annotated with a given functional term vs the total number of proteins quantified in the experiment. EndoC-βH1 cells were treated with control non-targeting siRNA and harvested at 72 h. Insulinoma samples from (39). Expression cut-off for proteins: quantified in >= 3 samples and unambiguously annotated.

### The secretome of EndoC- βH1 cells

In order to quality-control the secretome, we cross-referenced coordinates of peptides detected in the secretome experiments and mapped to membrane proteins with topology of proteins in the human surfaceome (42). If a secretome preparation is high-quality, peptides should map to extracellular regions that can be cleaved-off or shed. By contrast, contamination with peptides that map to transmembrane or intracellular regions indicates lysis or mechanical cell damage. The ratio of combined lengths of extracellular vs intracellular regions in the reference database was 2.4:1. This expected ratio was preserved in the cell lysate DIA experiment PXD027911 with 65.6% peptides mapped to extracellular regions and 30.9% peptides mapped to intracellular regions. By contrast, 94.4% peptides mapped to extracellular regions in PXD027913 and 93.2% in PXD027920 secretome experiments. Cell lysate proteome in PXD027911 and secretome in PXD027913 were generated based on the same EndoC-βH1 cell cultures. Expression abundances of proteins with cytoplasmic subcellular location according to UniProt (43) did not correlate between cell lysate and the corresponding secretome samples - median IQR Spearman rho 0.02 (−0.12 to 0.16), - indicating that leakage was unlikely. Proglucagon was not detected in the secretome experiments. Somatostatin was secreted by cells treated with control siRNA but not by untransfected EndoC-βH1 cells.

Higher number of proteins were detected in the secretome of EndoC-βH1 cells treated with control non-targeting siRNA than in untreated cells at baseline (**Fig 4A**).

**Fig. 4.**
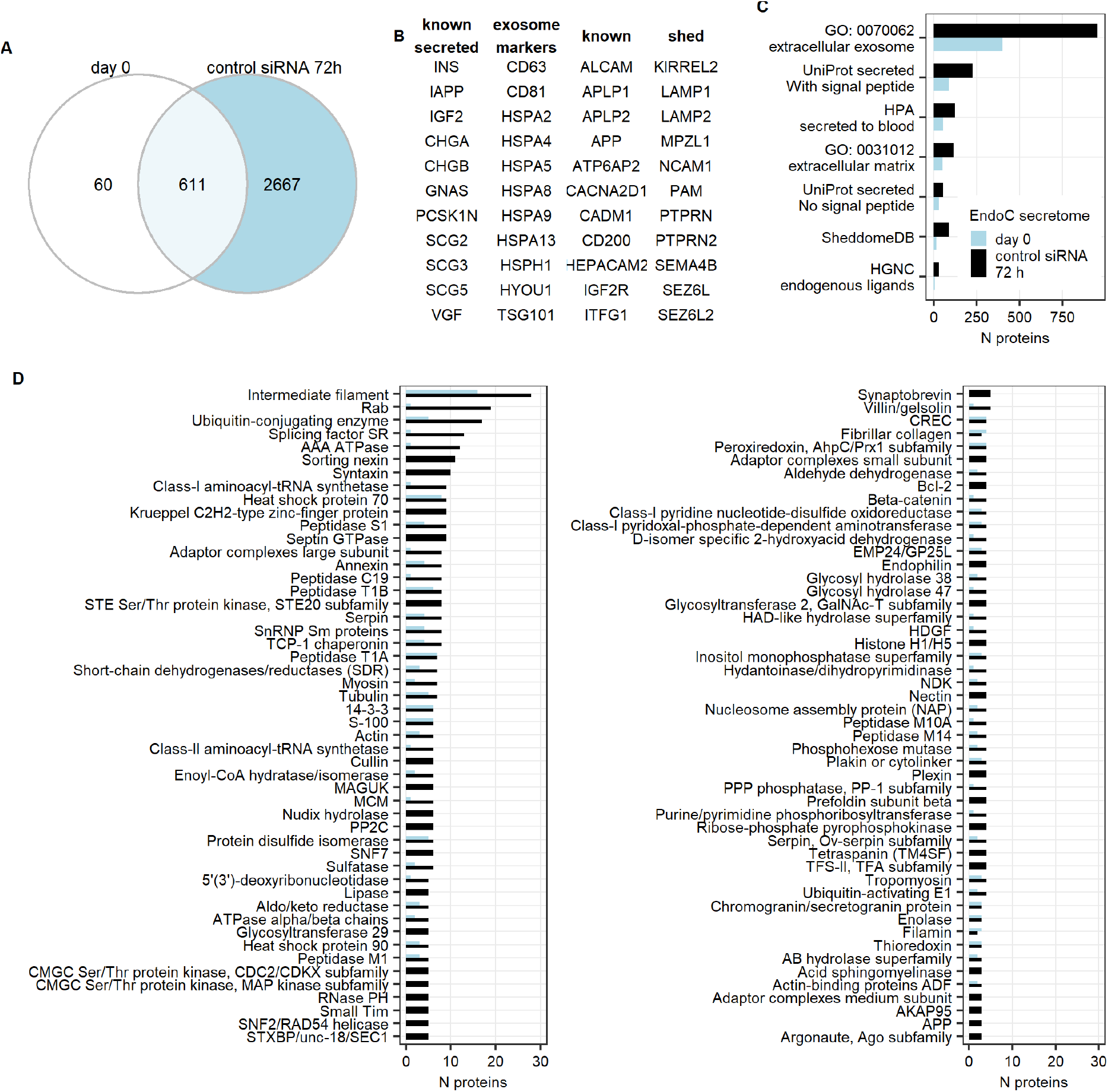
Composition of the EndoC-βH1 cell secretome. **(A)** Secretome of untreated cells compared to secretome of cells treated with control non-targeting siRNA for 72 hours. (**B**) Known positive control proteins that were quantified in EndoC cell secretome. (**C**) Number of proteins by mode of secretion. Proteins can belong to more than one category (e.g., secreted based on both UniProt and HPA annotation). (**D**) Top 50 protein families. Expression cut-off for proteins: quantified in min 2 out of 3 baseline samples, quantified in min 3 out of 12 samples at 72 hours.

EndoC-βH1 cells secreted known β cell hormones INS and IAPP and autocrine regulator IGF2 (44) (**Fig 4B**). The secretome contained proteins known to generate bioactive peptides such as all 8 proteins of the granin family (**Fig 4B**), among which chromogranins A and B, secretogranins 2 and 3 and VGF have been previously reported in insulinomas (45). Secretome proteomics indicated potential presence of several bioactive peptides (**Fig S16-S50**) including chromogranin A peptides pancreastatin and vasostatins, chromogranin B peptides GAWK and CCB, secretogranin-2 peptide manserin and VGF peptides. Some peptides were established β cell secretion products: IGF2 peptide preptin is co-secreted with insulin (46) while VGF peptides neuroendocrine regulatory peptide-2 (47) and TLQP-62 (48) regulate glucose-induced-insulin-secretion. The secretome also contained CPE, PAM and proprotein convertases PCSK1 and PCSK2 - the enzymes involved in generation of bioactive peptides from prohormones. Interestingly, EndoC-βH1 cells also secreted PCSK9, a member of proprotein convertase family without known enzymatic function but involved in LDL receptor clearance in hepatocytes. We further investigated the function of PCSK9 in EndoC-βH1 cells in a separate follow-up study [**reference**].

We quantified exosome markers (49) in the secretome samples - CD63, CD81, heat shock 70kDa proteins and TSG101 (**Fig 4B**) and members of the Rab family (**Fig 4D**). Exosomal proteins were a major component of the secretome (**Fig 4C**). Secretion of exosomes is increased in stressed β cells (50). Therefore, higher number of protein species quantified in EndoC-βH1 cells treated with control siRNA than in untreated cells was consistent with exosomes being the major “mode of secretion”.

The EndoC-βH1 cell secretome after 72 hours of incubation with control siRNA contained proteins implicated in β cell proliferation: NR3C1 (glucocorticoid receptor), GSK3B and cell cycle-regulating kinases CDK1, CDK2, CDK4 and CDK6. These proteins can be detected in exosomes based on ExoCarta database (51), although most of the referenced research works were conducted in cancer tissues or cell lines rather than non-cancerous tissues. Hence, it might be theoretically possible to infer β cell state based on exosome markers that are normally intracellular and inaccessible for measurement in plasma or serum.

The second most common category of proteins in the secretome were classically secreted proteins containing signal peptide (**Fig 4C**). For example, both untreated and control siRNA treated EndoC-βH1 cells secreted factors regulating cell differentiation and survival NENF, MANF, MYDGF and CREG1. EndoC-βH1 cells also secreted proteins that are measurable in circulation (52) such as annexins, apolipoproteins, serpins, IGF-binding proteins and VEGFA.

In addition, we captured known proteins with shed extracellular domain in islets or rodent cell lines (21, 53, 54) in the EndoC-βH1 cell secretome (**Fig 4B**). Sheddases ADAM10, ADAM17, ADAM22 and ADAM9 were quantified in the EndoC-βH1 secretome. Sheddases BACE1 and BACE2 were not quantified in the secretome samples, but were quantified in the DIA and TMT cell lysate experiments.

Finally, we investigated the most abundantly secreted proteins in the untreated (**Table S2**) and control siRNA treated samples (**Table 1**). β cell hormones were among the most abundant secreted proteins confirming β cell line identity.

**Table 1.**
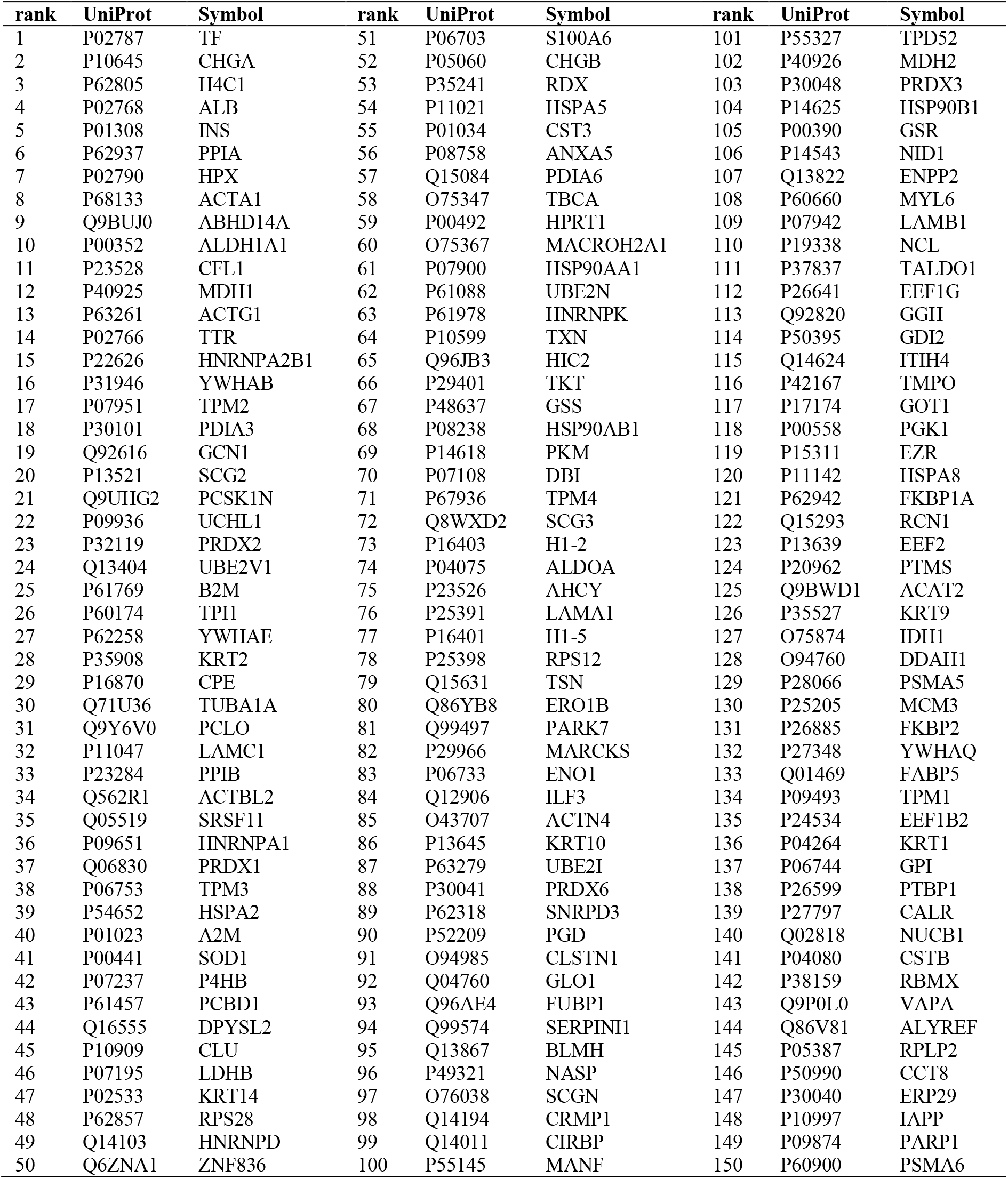
Top 150 secreted proteins. Proteins are ordered by decreasing median normalized expression abundance in EndoC-βH1 cells treated with control non-targeting siRNA at 72 hours in the DIA secretome experiment (PXD027913). N=12.

## DISCUSSION

EndoC-βH1 cells are among the most commonly used human β cell lines for early-stage diabetes research, yet EndoC-βH1 cells have not been validated as an *in vitro* human β cell model based on omics. Here, we confirm that EndoC-βH1 are cells of β cell lineage using transcriptomics and proteomics data. The observed discrepancies with adult human β cells and increased similarity with insulinomas were due to elevated proliferation rate (14). Therefore, inducible growth arrest will likely ameliorate the discrepancies in gene and protein expression pattern. In fact, next generation EndoC-βH2 and EndoC-βH3 cells display dramatic increase in insulin content and secretion capacity after the immortalization cassette is excised (10, 11). Increased expression of β cell markers post-excision has also been characterized in second-generation EndoC cells (10).

Limited detection of GPCRs in the proteomics experiments might be of technical nature. In general, membrane proteins are hydrophobic. Much harsher lysis buffers than the buffers were used in these proteomics experiments are required for solubilizing GPCRs. In addition, the hydrophobic nature of GPCRS leads to low frequency of polar aminoacids (i.e., very few lysin and arginine residues), which affects tryptic digestion and subsequent detection by mass-spectrometry (55, 56). By contrast, low GLP1R expression appeared to be a true biological finding. GLP1R expression may also be remedied by using next-generation cell lines. For example, dose-dependent response to GLP-1 mimetics has been reported by vendors in Endoc-βH5 cells (https://www.humancelldesign.com/human-beta-cells-endoc-bh5/).

In summary, growth-arrested next-generation EndoC cells may be more suitable for some research questions such as studies of factors influencing proliferation of adult β cells, GLP1R-mediated drug delivery or effects GLP-1 mimetics.

As EndoC-βH1 cells proved to be a reasonable translational human β cell model, we used EndoC-βH1 cells to uncover the β cell secretome. EndoC-βH1 cells secreted β cell hormones INS and IAPP. Both adult human β cells and insulinoma cells secrete low amounts of insulin under basal conditions and not only after glucose or KCl challenge (57, 58). Hence, presence of insulin and co-secreted products IAPP and IGF2/preptin in the secretome of unstimulated EndoC-βH1 is not abnormal. Albumin was not quantified in the EndoC-βH1 RNA-seq samples. Therefore, albumin in the secretome samples likely represents the traces of albumin derived from the culture medium where the cells had been previously grown.

Our secretome proteomics workflow was aimed at identification of secreted proteins and contained a tryptic digestion step. Thus, we could not establish full-length sequences of endogenously cleaved peptides. However, we could make inference based on the spectra and prior knowledge of protein families generating bioactive peptides. Our secretome data indicated that β cells may be capable of producing and secreting bioactive peptides. In fact, we confirmed the presence of preptin, neuroendocrine regulatory peptide-2 and TLQP-62 (46–48) in conditioned medium from EndoC-βH1 cells. We also found that EndoC-βH1 cells secrete proteins of the granin family and possess the enzymatic machinery necessary to produce bioactive peptides, which suggests that the endogenous bioactive peptide repertoire of β cells might be richer than previously anticipated. Bioactive peptides may play an important role in both autocrine and endocrine signaling as well as cross-talk between β cells and other islets cell types. We believe that presence of full-length endogenous peptides and their secretion dynamics and function warrant further investigation in subsequent studies with specialized proteomics methods.

The secretome of EndoC-βH1 encompassed classically secreted proteins with signal peptide. However, we speculate that they may play a limited role as biomedically relevant biomarkers. Only a few proteins in the secretome had enriched expression in β cells (INS, IAPP) or nervous system and β cells (e.g. PTPRN, VGF) (59). In addition, the relative contribution of β cells to the plasma pool of broadly expressed proteins in human is unknown, and deconvoluting tissue-of-origin for such blood-borne biomarkers can be extremely challenging. By contrast, classically secreted proteins may play an autocrine or paracrine function. For example, the functions of NENF, MANF, MYDGF and CREG1 as regulators of cell survival and differentiation were described in a different tissue context (43) but they may play similar roles in islets. In fact, MANF has been recently characterized as a β cell protecting factor (60).

Abundance of exosome-associated proteins in untreated and control siRNA treated cells was an unexpected and potentially biomedically relevant finding. Exosomes may offer additional opportunities to discover and non-invasively monitor biomarkers reflecting β cell state. Another important observation was quantification of CD63 and CD81 in the EndoC-βH1 cell secretome. Antibodies recognizing these exosome surface markers are used in exosome isolation protocols, and our findings indicate that standard protocols could be applicable to isolate β cell derived exosomes in *in vitro* studies. Future studies need to further elucidate composition of β cell exosomes and their secretion dynamics, and to establish if it is possible to enrich β cell derived exosomes from plasma, serum or fecal samples based on additional biomarkers.

### Data availability

#### Data sets will be made public upon publication of the manuscript in a peer-reviewed journal

RNA-seq data have been deposited to NCBI GEO (61) with accession GSE182016

The mass spectrometry proteomics data have been deposited to the ProteomeXchange Consortium via the PRIDE (62) partner repository with the following identifiers: PXD027898, PXD027921, PXD027920, PXD027911 and PXD027913. Each data set contains raw files, annotated spectra files in .sne or .msf format, files with parameters for Proteome Discoverer or Spectronaut software run and a processed_data.zip. Each processed_data.zip files contain raw protein quantification table, log2 normalized and imputed protein quantification table, sample description and analysis log file. Peptide quantification tables and the corresponding log files are included in PXD027911, PXD027913 and PXD027920.

Gene/protein lists corresponding to Figure 2 and Figure 4 are provided in the supplementary excel file

The previously published data sets used as reference primary human biological material were: GTEx (59), Human Protein Atlas (HPA) (52), GSE67543 (36), GSE57973 (63), GSE118014 (38), GSE84133 (64), GSE81608 (65), E−MTAB−5061 (66), GSE86469 (67), PXD007569 (37). Marker lists were obtained from HGNC (68), OMIM (69), GENECODE v36 (70) and supplementary data sets in (39, 71–74).

## Abbreviations

ACN: acetonitrile
DIA: Data Independent Acquisition
FA: formic acid
FACS: Fluorescence-activated cell sorting
FDR: False Discovery Rate
GLP1R: glucagon like peptide 1 receptor
GO: Gene Ontology
GPCR: G-protein coupled receptor
HCD: Higher-energy C-trap dissociation
HPRP Fractionation: High pH Reversed-Phase Fractionation
LC: liquid chromatography
MS/MS: tandem mass spectrometry
PBS: Phosphate Buffered Saline
SDB-RPS: styrene divinylbenzene reversed-phase
T2D: Type 2 Diabetes
TMT: Tandem Mass Tag
TPM: transcripts per million

## Author contribution statements

R.S., S.A., K.Se., G.H. - design and planning of the study. K.S. - cell culture, RNA-seq and FACS experiments. C.B. - FACS experiments. G.H., A.J. - mass-spectrometry data acquisition, pre-processing. M.R. - down-stream analysis, first draft. All authors - data interpretation, scientific feedback, revisions of the manuscript and approval of the submitted version.

