## Supplemental Figures for "Characterization of the secretome, transcriptome and proteome of human β cell line EndoC-βH1"

**Table S1. Detectability of expression across gene categories.** The table shows the number of markers with median expression >= 1 TPM in adult β cells (GSE67543, N=7, FACS sorted, at least 97% purity) and EndoC-βH1 cells. Percentage is given as the number of markers expressed above the threshold in both EndoC-βH1 cells and adult β cells vs in adult β cells. Total N markers is the total number of genes in the gene category. E.g., total number of human enzymes according to HGNC (December 2020).

| Category | Total N markers | Adult β cells | EndoC | Both | Percent |
|---|---|---|---|---|---|
| β cell markers | 128 | 128 | 116 | 116 | 90.62 |
| OMIM MODY and/or familial hyperinsulinemic hypoglycemia | 19 | 10 | 10 | 9 | 90 |
| GWAS T2D or glycemic traits | 56 | 41 | 37 | 32 | 78.05 |
| Transcription factors | 1639 | 1035 | 1105 | 932 | 90.05 |
| Enzymes | 1797 | 1240 | 1340 | 1132 | 91.29 |
| Ion channels | 328 | 126 | 134 | 99 | 78.57 |
| GPCRs | 1421 | 120 | 115 | 62 | 51.67 |
| Polyadenylated lncRNA | 2042 | 198 | 442 | 137 | 69.19 |

**Fig. S1. $\beta$ cell markers.** Markers were defined based on elevated expression Z-score in $\beta$ cells vs other cell types in at least two out of four single-cell RNA-seq studies GSE84133, GSE81608, E-MTAB-5061 and GSE86469. **(A)** Gene expression in adult $\beta$ cells (GSE67543, N=7, FACS sorted, at least 97% purity). Markers are ordered by decreasing median expression. **(B)** Gene expression in EndoC-$\beta$H1 cells treated with control siRNA at 72 hours (N=3). **(C)** Protein expression in DIA proteomics on cell lysate of EndoC-$\beta$H1 cells treated with control siRNA at 72 hours (N=12). **(D)** Protein expression in TMT proteomics on cell lysate of EndoC-$\beta$H1 cells treated with control siRNA at 72 hours (N=3). No boxplot for a protein in panels C and D indicates that the protein was either not detected or quantified in $< 3$ samples.

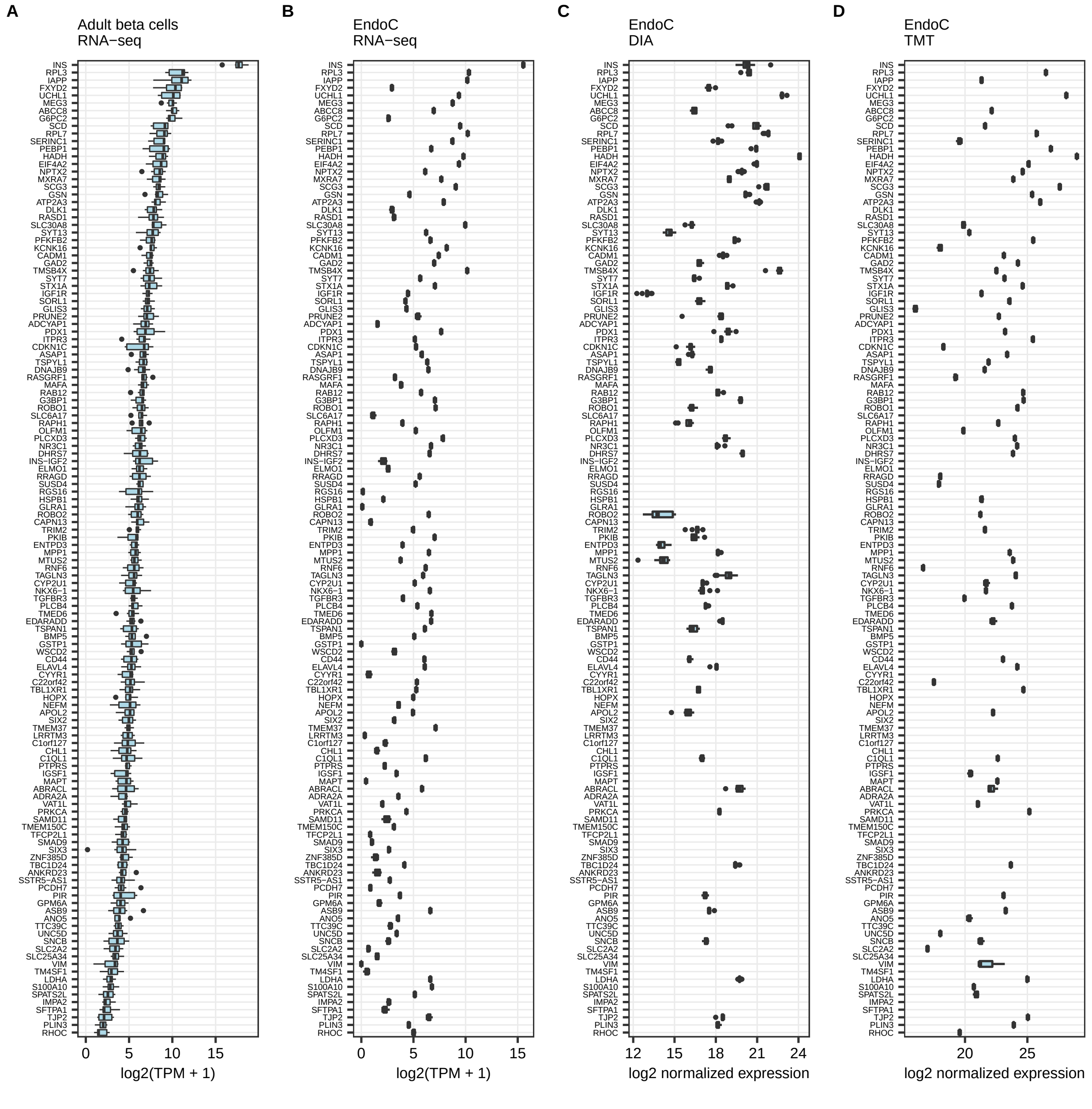

**Fig. S2. Genes implicated in maturity-onset diabetes of the young and/or familial hyperinsulinemic hypoglycemia (OMIM). (A)** Gene expression in adult $\beta$ cells (GSE67543, N=7, FACS sorted, at least 97% purity). **(B)** Gene expression in EndoC-$\beta$H1 cells treated with control siRNA at 72 hours (N=3). **(C)** Protein expression in DIA proteomics on cell lysate of EndoC-$\beta$H1 cells treated with control siRNA at 72 hours (N=12). **(D)** Protein expression in TMT proteomics on cell lysate of EndoC-$\beta$H1 cells treated with control siRNA at 72 hours (N=3). No boxplot for a protein in panels C and D indicates that the protein was either not detected or quantified in $<$ 3 samples.

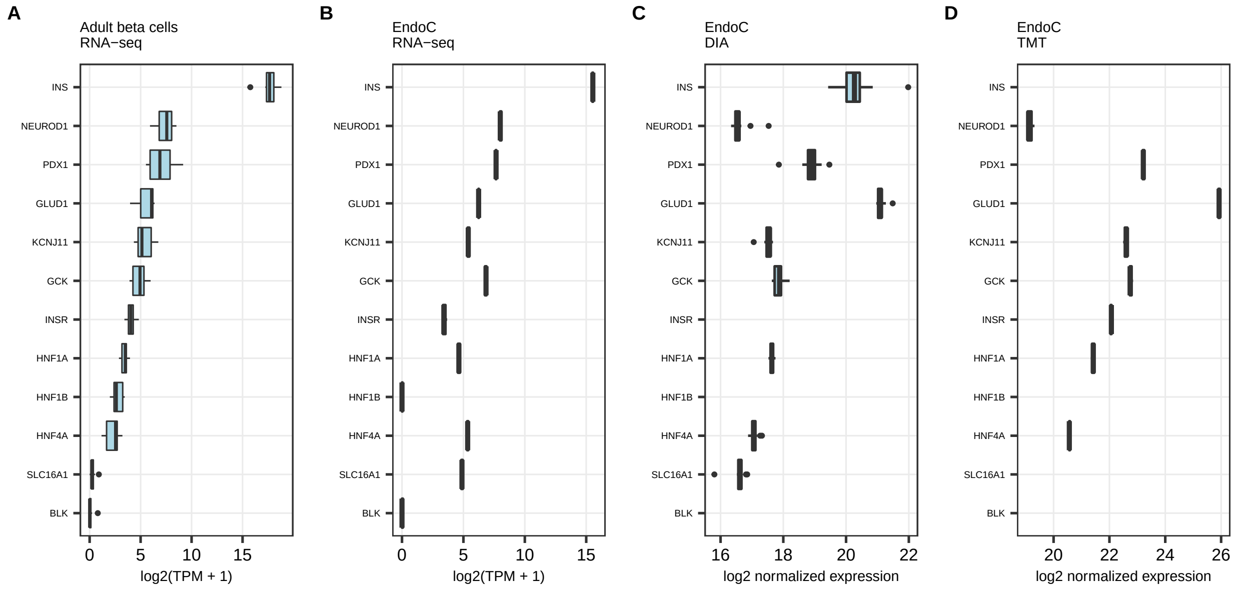

**Fig. S3. GWAS genes associated with T2D and glycemic traits.** Markers were extracted from Supplementary Data 15 in Viñuela et al. 2020 (GWAS-to-eQTL co-localization in islets). **(A)** Gene expression in adult $\beta$ cells (GSE67543, N=7, FACS sorted, at least 97% purity). **(B)** Gene expression in EndoC-$\beta$H1 cells treated with control siRNA at 72 hours (N=3). **(C)** Protein expression in DIA proteomics on cell lysate of EndoC-$\beta$H1 cells treated with control siRNA at 72 hours (N=12). **(D)** Protein expression in TMT proteomics on cell lysate of EndoC-$\beta$H1 cells treated with control siRNA at 72 hours (N=3). No boxplot for a protein in panels C and D indicates that the protein was either not detected or quantified in $<$ 3 samples.

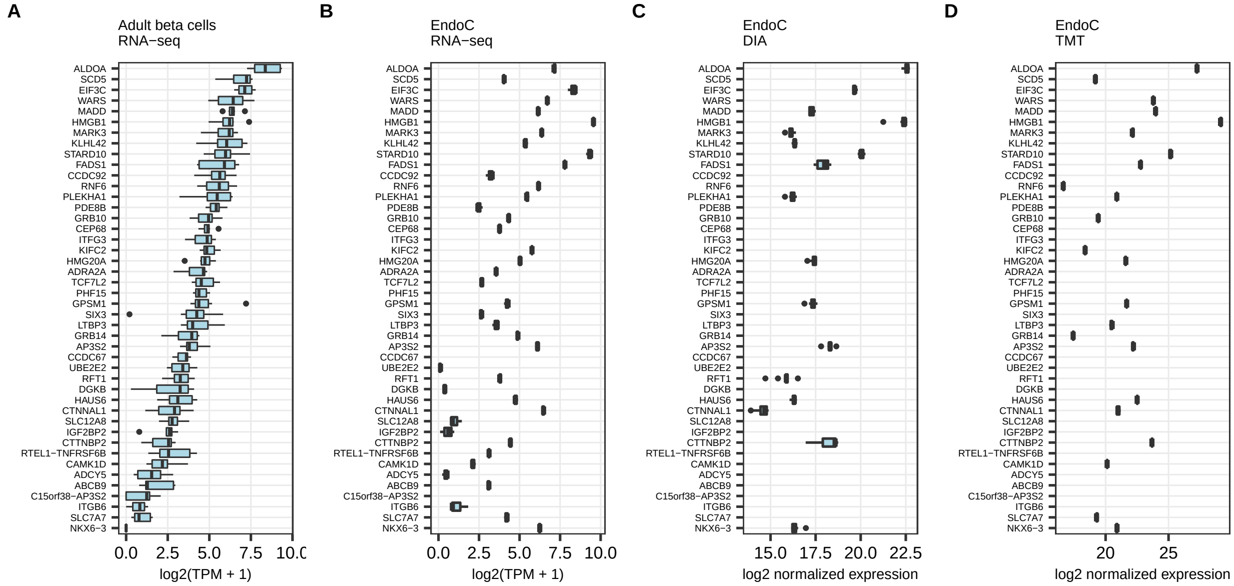

**Fig. S4. Transcription factors.** Genes were extracted from supplementary data in Lambert et al. 2018. **(A)** Gene expression in adult $\beta$ cells (GSE67543, N=7, FACS sorted, at least 97% purity). Top 130 genes by median expression. **(B)** Gene expression in EndoC-$\beta$H1 cells treated with control siRNA at 72 hours (N=3). **(C)** Protein expression in DIA proteomics on cell lysate of EndoC-$\beta$H1 cells treated with control siRNA at 72 hours (N=12). **(D)** Protein expression in TMT proteomics on cell lysate of EndoC-$\beta$H1 cells treated with control siRNA at 72 hours (N=3). No boxplot for a protein in panels C and D indicates that the protein was either not detected or quantified in $<$ 3 samples.

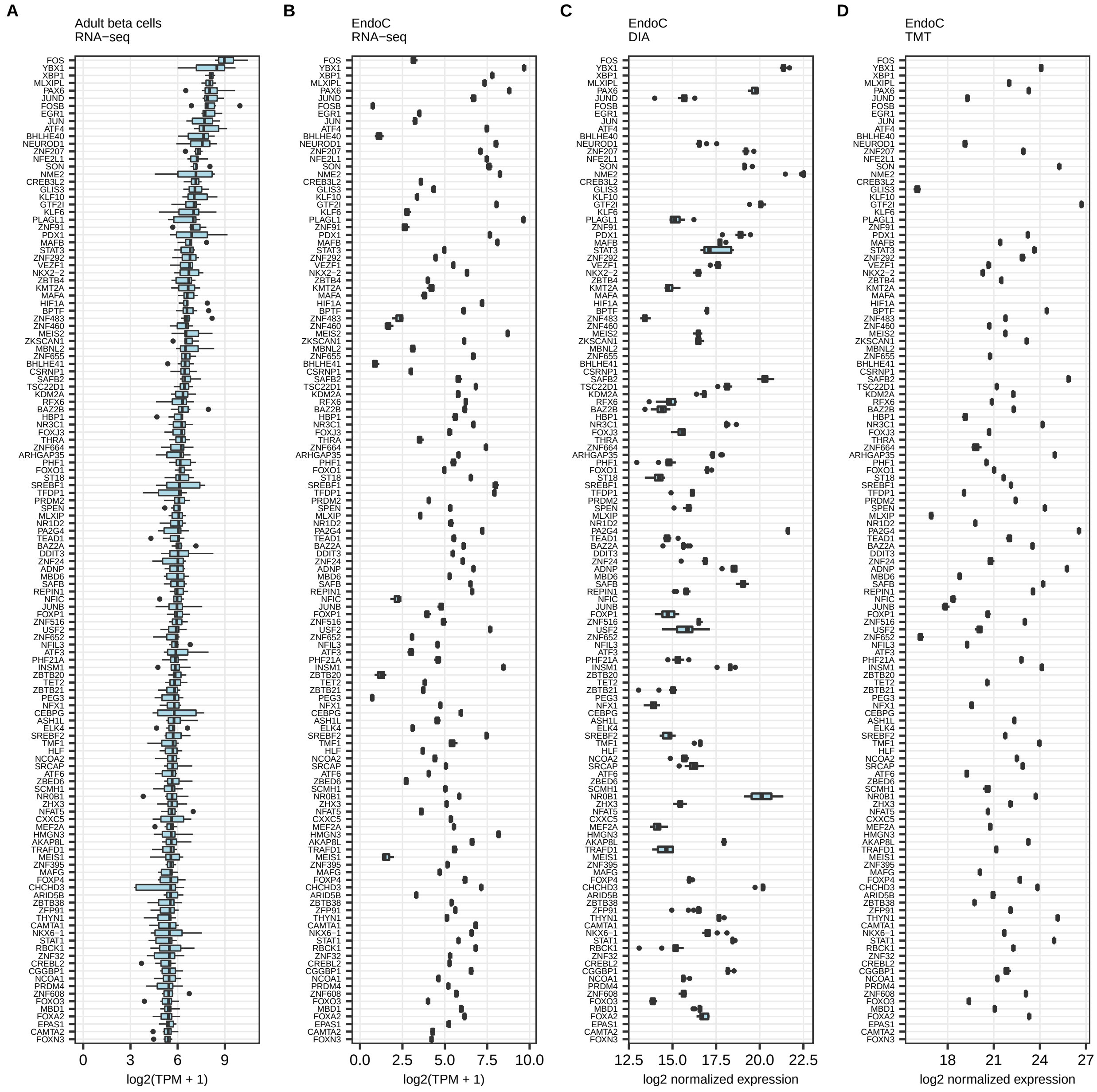

**Fig. S5. Enzymes.** Gene list was obtained from HGNC. **(A)** Gene expression in adult $\beta$ cells (GSE67543, N=7, FACS sorted, at least 97% purity). Top 130 genes by median expression. **(B)** Gene expression in EndoC-$\beta$H1 cells treated with control siRNA at 72 hours (N=3). **(C)** Protein expression in DIA proteomics on cell lysate of EndoC-$\beta$H1 cells treated with control siRNA at 72 hours (N=12). **(D)** Protein expression in TMT proteomics on cell lysate of EndoC-$\beta$H1 cells treated with control siRNA at 72 hours (N=3). No boxplot for a protein in panels C and D indicates that the protein was either not detected or quantified in $<$ 3 samples.

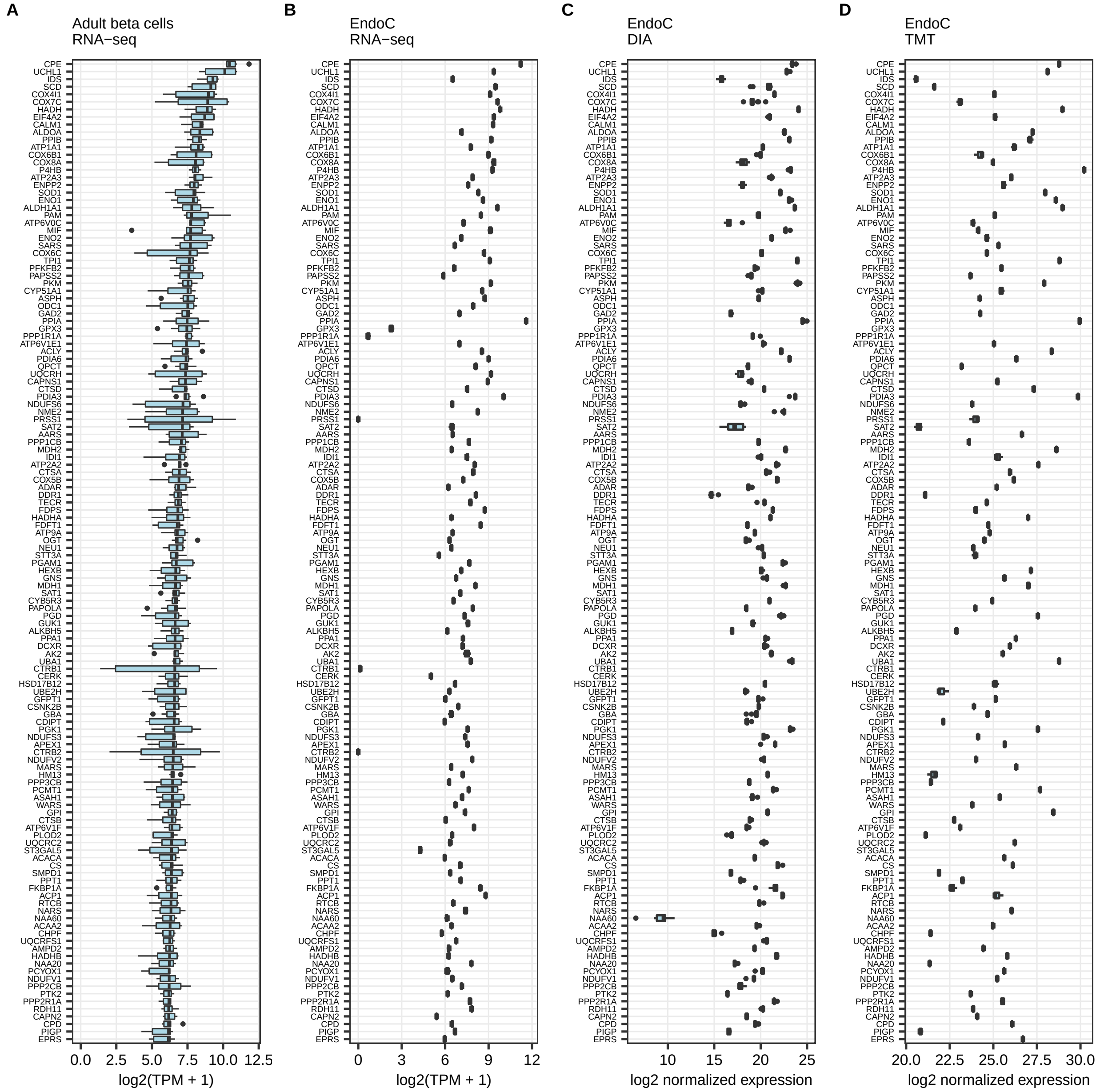

**Fig. S6. Ion channels.** Gene list was obtained from HGNC. **(A)** Gene expression in adult $\beta$ cells (GSE67543, N=7, FACS sorted, at least 97% purity). **(B)** Gene expression in EndoC-$\beta$H1 cells treated with control siRNA at 72 hours (N=3). **(C)** Protein expression in DIA proteomics on cell lysate of EndoC-$\beta$H1 cells treated with control siRNA at 72 hours (N=12). **(D)** Protein expression in TMT proteomics on cell lysate of EndoC-$\beta$H1 cells treated with control siRNA at 72 hours (N=3). No boxplot for a protein in panels C and D indicates that the protein was either not detected or quantified in $< 3$ samples.

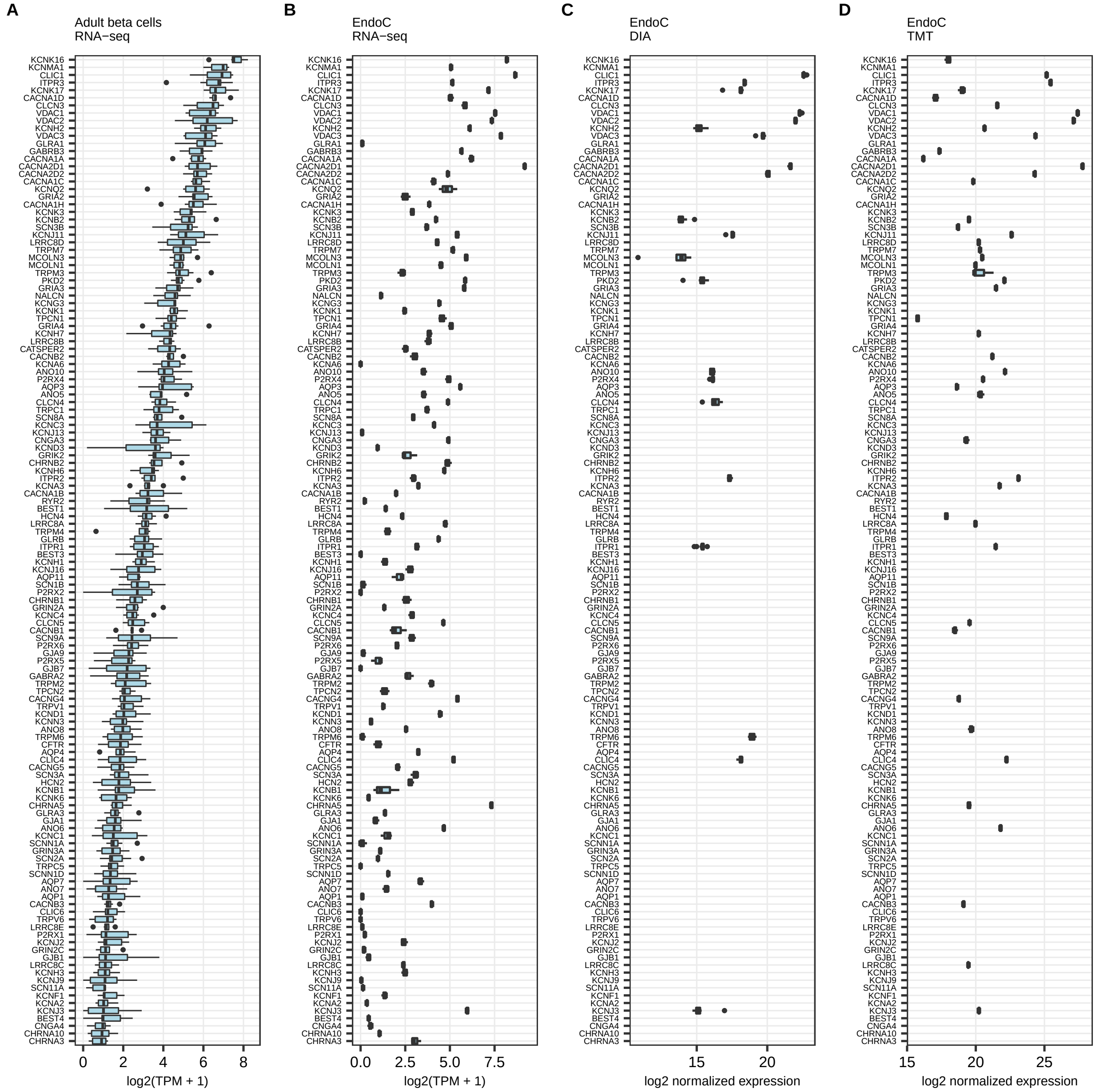

**Fig. S7. G protein-coupled receptors.** Gene list was obtained from HGNC. **(A)** Gene expression in adult $\beta$ cells (GSE67543, N=7, FACS sorted, at least 97% purity). **(B)** Gene expression in EndoC-$\beta$H1 cells treated with control siRNA at 72 hours (N=3). **(C)** Protein expression in DIA proteomics on cell lysate of EndoC-$\beta$H1 cells treated with control siRNA at 72 hours (N=12). **(D)** Protein expression in TMT proteomics on cell lysate of EndoC-$\beta$H1 cells treated with control siRNA at 72 hours (N=3). No boxplot for a protein in panels C and D indicates that the protein was either not detected or quantified in $<$ 3 samples.

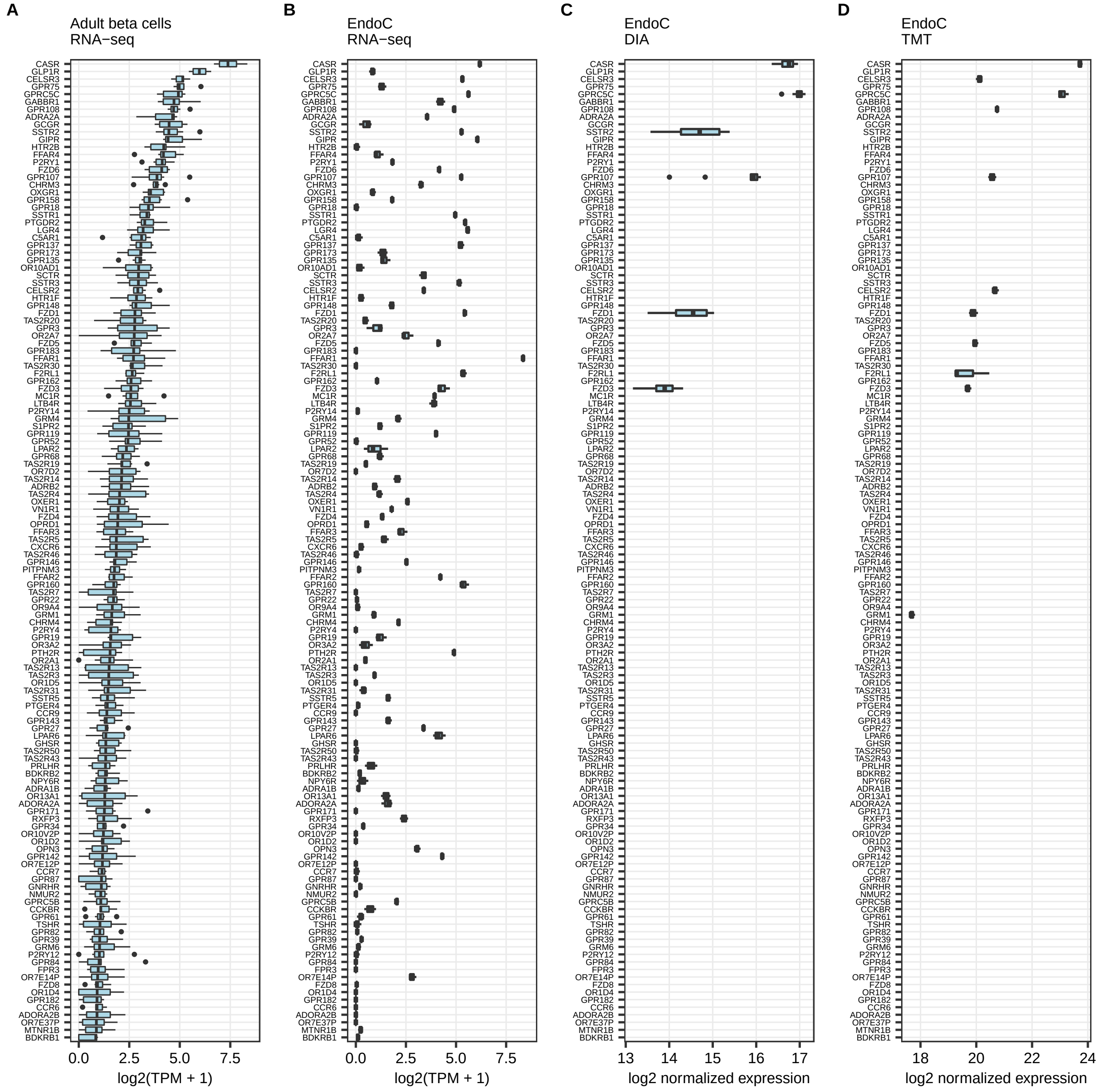

**Fig. S8. Polyadenylated lncRNA.** Gene list was obtained by cross-referencing HGNC with GENECODE v36 polyadenylation features. **(A)** lncRNA with median expression >=1 TPM in adult $\beta$ cells (GSE67543, N=7, FACS sorted, at least 97% purity). **(B)** Expression of the corresponding genes in EndoC-$\beta$H1 cells treated with control siRNA at 72 hours (N=3).

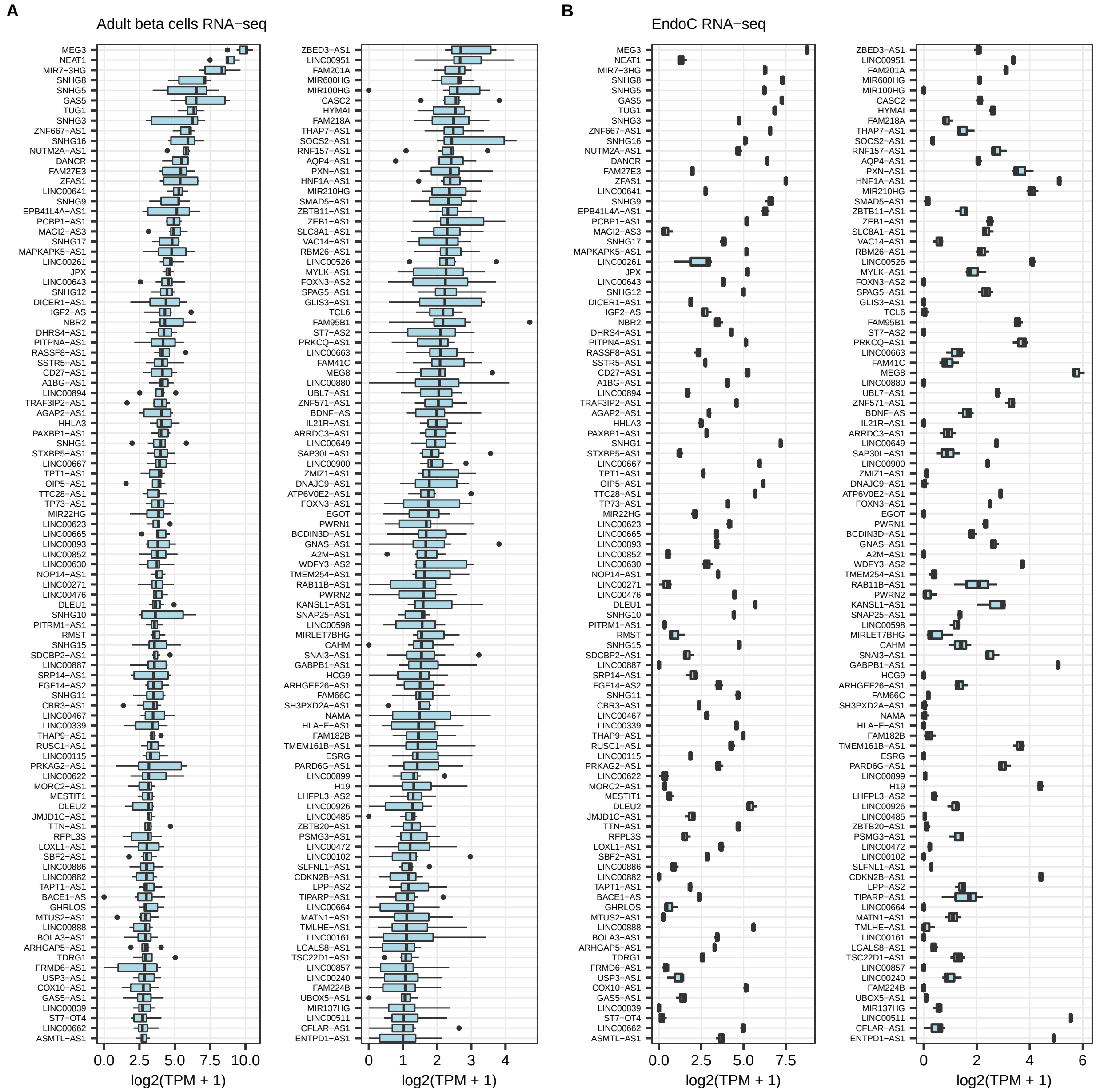

**Fig. S9. Glucagon and somatostatin expression compared to insulin in single-cell RNA seq studies on human islets.** Cells from non-diabetic donors in **(A)** GSE84133, **(B)** GSE81608, **(C)** E-MTAB-5061 and **(D)** GSE86469.

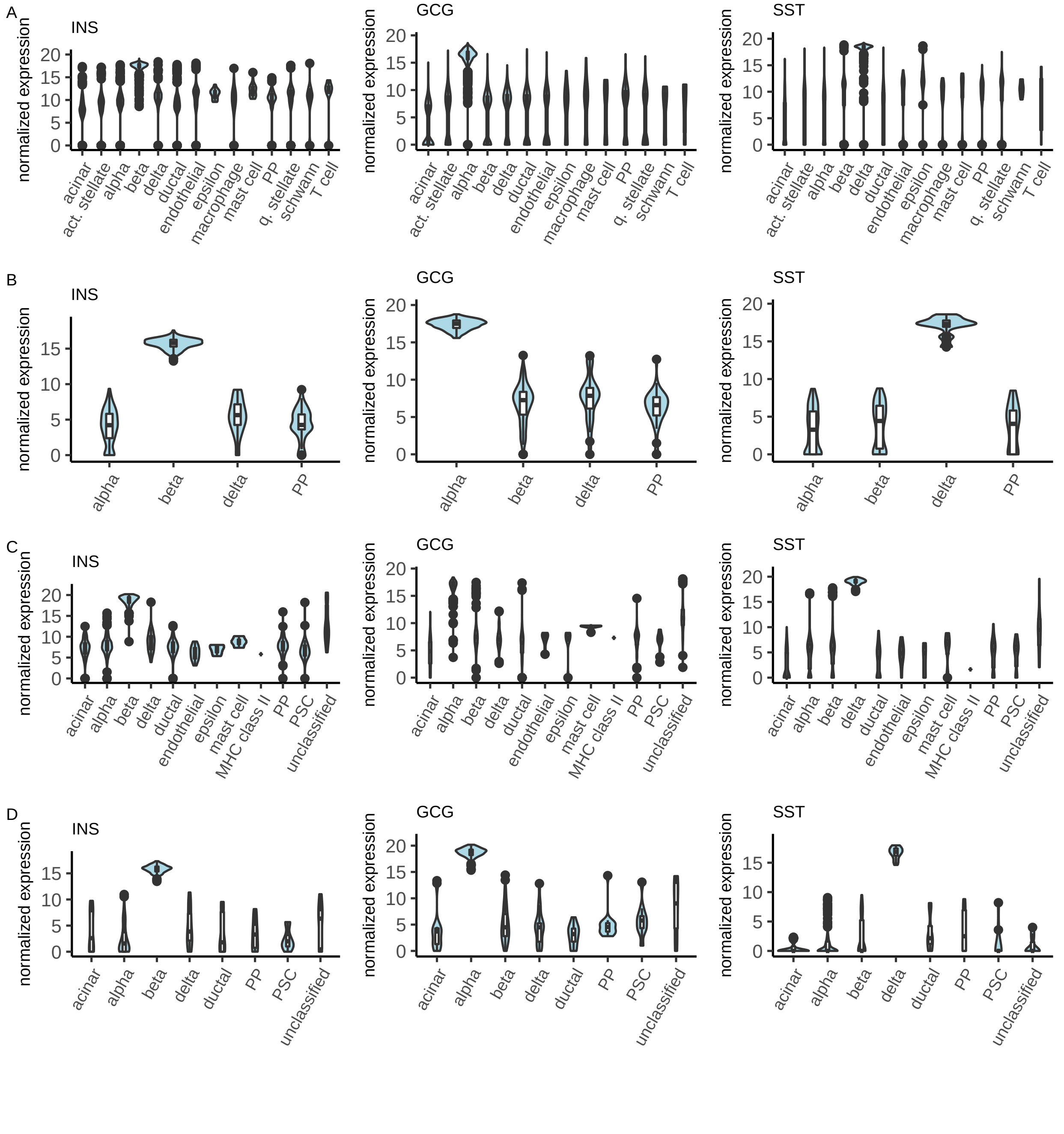

**Fig. S10. EndoC-$\beta$H1 cells and adult $\beta$ cells expressed $\alpha$ cell markers but at lower levels than $\alpha$ alpha cells. (A)** Gene expression in adult $\alpha$ cells (GSE67543, N=7, FACS sorted, at least 97% purity). **(B)** Gene expression in adult $\beta$ cells (GSE67543, N=7, FACS sorted, at least 97% purity). **(C)** Gene expression in EndoC-$\beta$H1 cells treated with control siRNA at 72 hours (N=3). **(D)** Protein expression in DIA proteomics on cell lysate of EndoC-$\beta$H1 cells treated with control siRNA at 72 hours (N=12). **(E)** Protein expression in TMT proteomics on cell lysate of EndoC-$\beta$H1 cells treated with control siRNA at 72 hours (N=3). No boxplot for a protein in panels C and D indicates that the protein was either not detected or quantified in $<$ 3 samples.

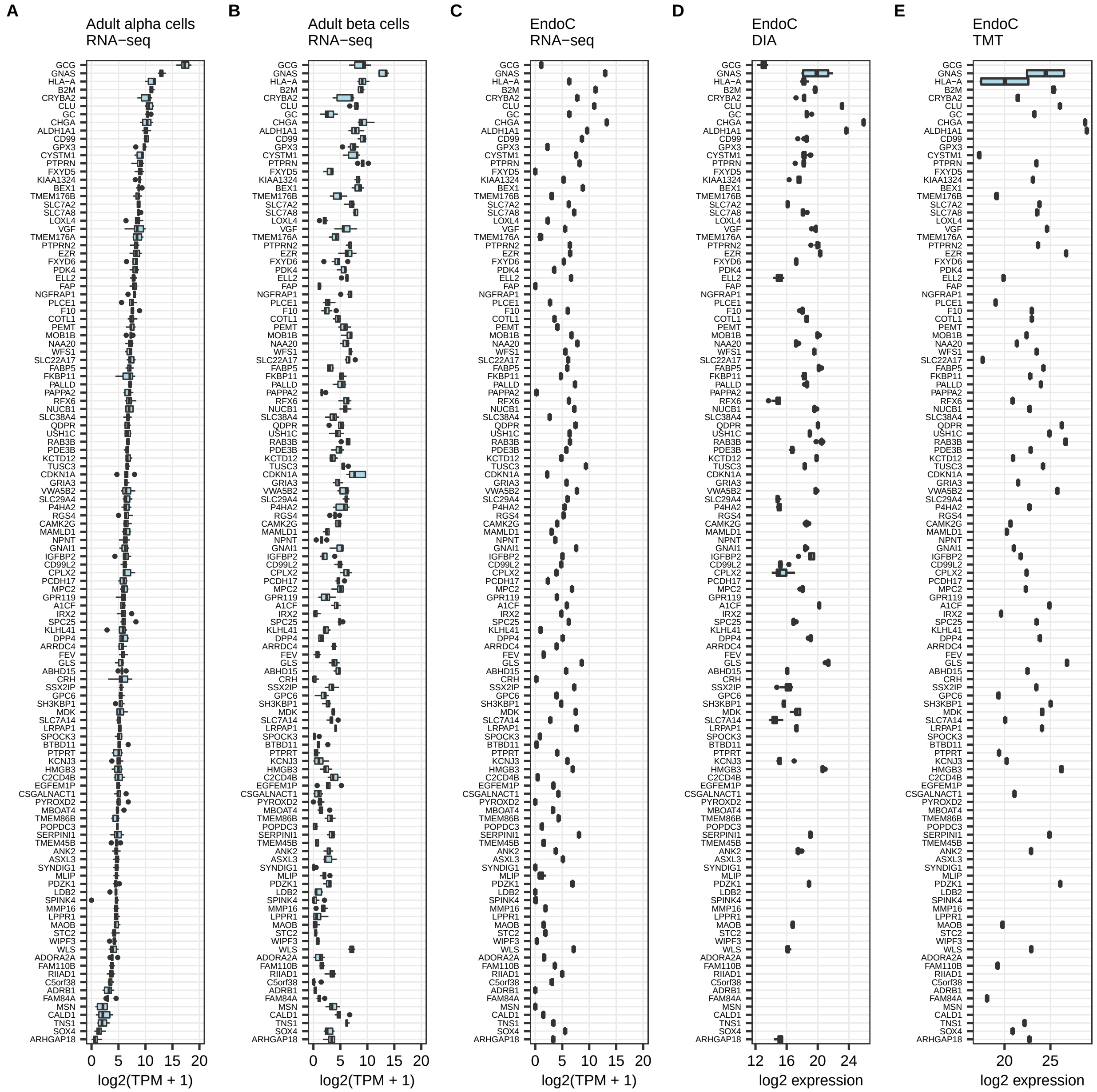

**Fig. S11. EndoC-$\beta$H1 cells had reduced expression of progenitor markers.** Endocrine progenitor markers were defined as markers of ECAD$^{low}$CD142$^{neg}$SUSD2$^{pos}$ and ECAD$^{low}$CD142$^{neg}$SUSD2$^{neg}$ in fetal pancreas at 9 weeks of gestation from Ramond et al. 2018 and median expression $<$ 1 TPM in adult $\beta$ cells (GSE67543). Marker mRNA expression in **(A)** fetal pancreas 9 weeks, **(B)** adult $\beta$ cells, **(C)** EndoC-$\beta$H1 cells treated with control siRNA at 72 hours (N=3). **(D)** Protein expression in DIA proteomics on cell lysate of EndoC-$\beta$H1 cells treated with control siRNA at 72 hours (N=12). **(E)** Protein expression in TMT proteomics on cell lysate of EndoC-$\beta$H1 cells treated with control siRNA at 72 hours (N=3). No boxplot for a protein in panels C and D indicates that the protein was either not detected or quantified in $<$ 3 samples.

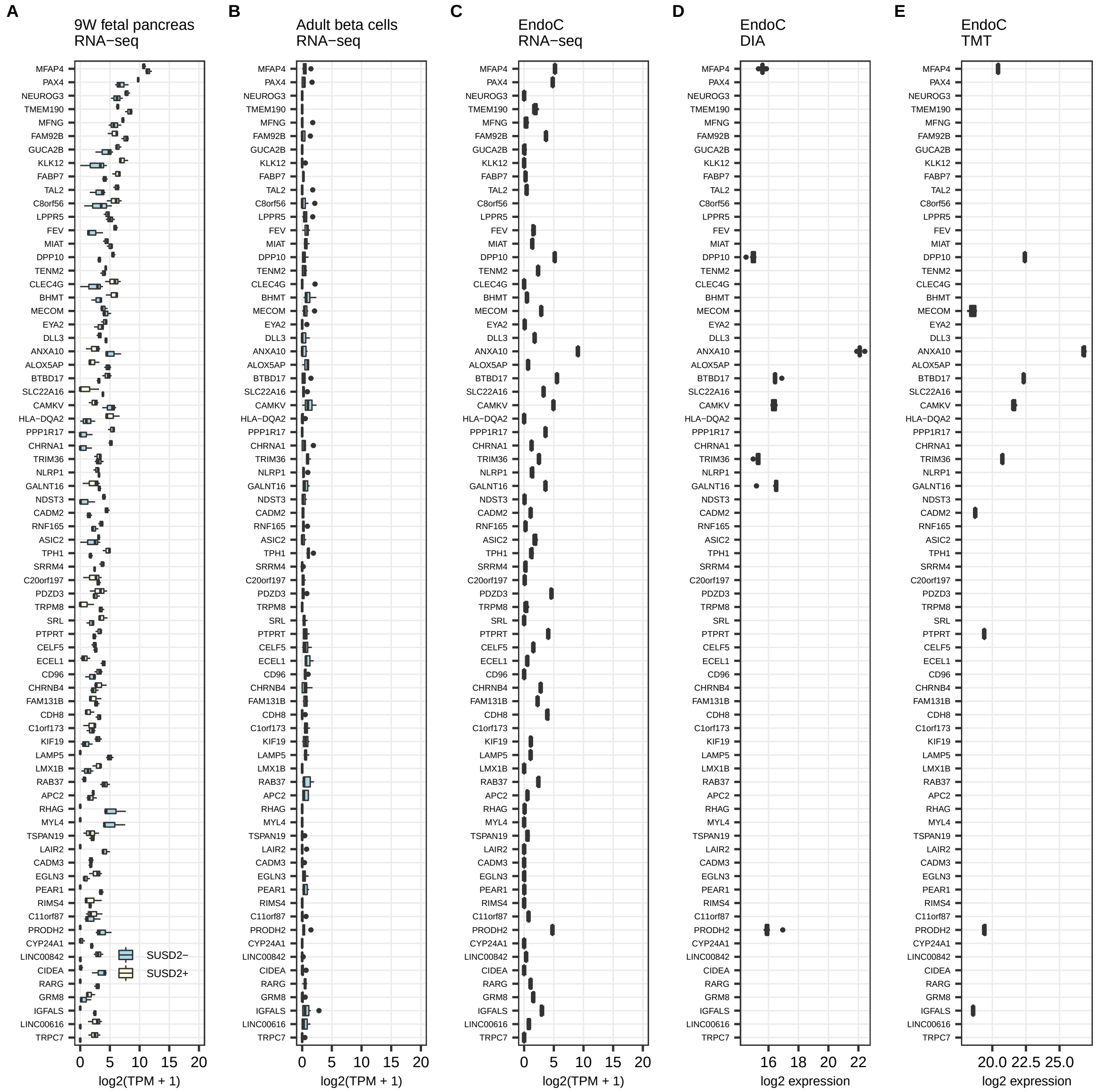

**Fig. S12. EndoC-$\beta$H1 cells and adult $\beta$ cells did not express fetal $\beta$ cell markers. (A)** Gene expression in fetal $\beta$ cells (GSE67543, 12 to 18 weeks gestation, N=6). **(B)** Gene expression in adult $\beta$ cells (GSE67543, N=7, FACS sorted, at least 97% purity). **(C)** Gene expression in EndoC-$\beta$H1 cells treated with control siRNA at 72 hours (N=3). **(D)** Protein expression in DIA proteomics on cell lysate of EndoC-$\beta$H1 cells treated with control siRNA at 72 hours (N=12). **(E)** Protein expression in TMT proteomics on cell lysate of EndoC-$\beta$H1 cells treated with control siRNA at 72 hours (N=3). No boxplot for a protein in panels C and D indicates that the protein was either not detected or quantified in $<$ 3 samples.

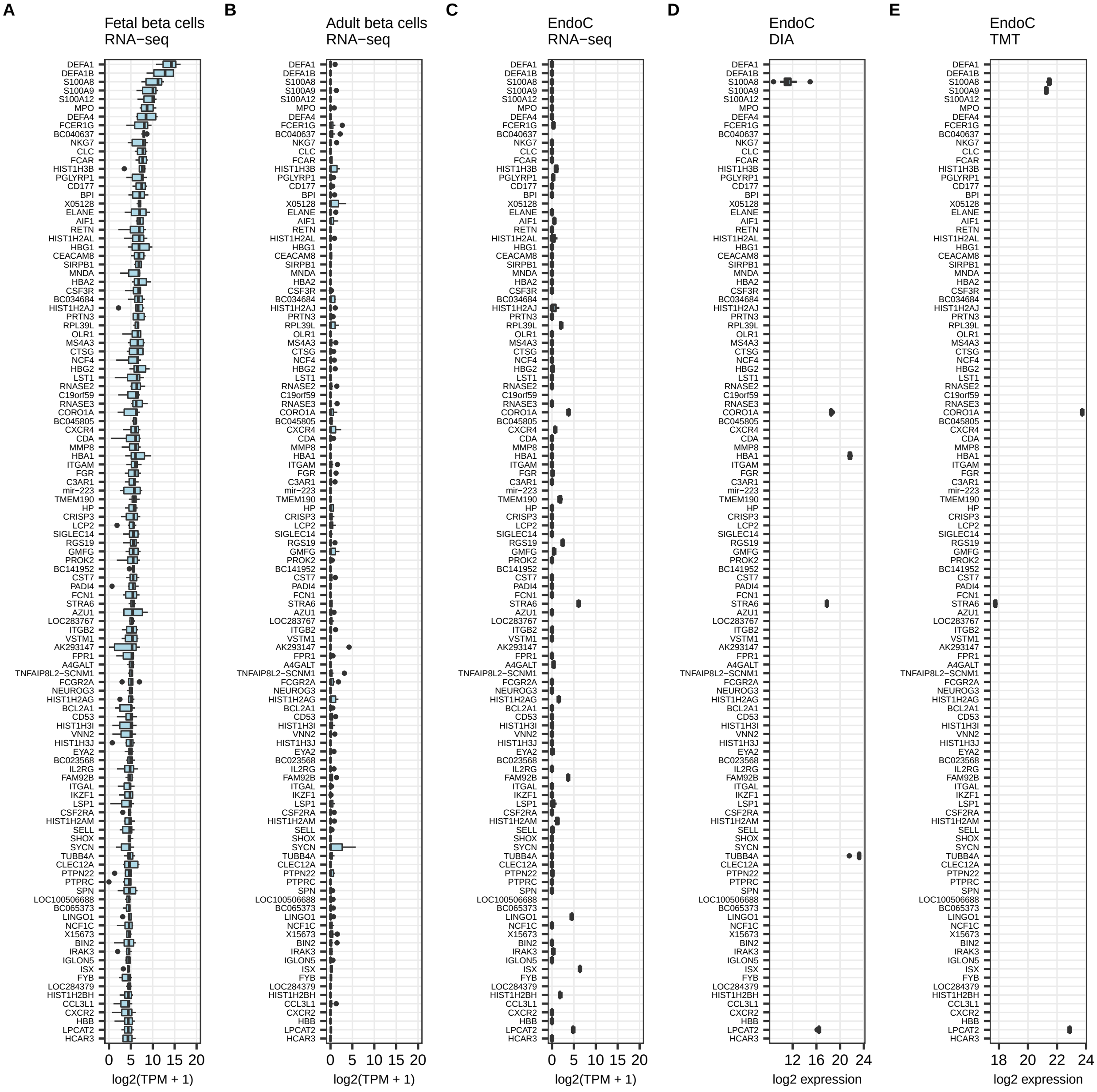

**Fig. S13. EndoC-$\beta$H1 cells did not express acinar cell markers. (A)** Gene expression in adult acinar cells (GSE57973, N= 2). **(B)** Gene expression in adult $\beta$ cells (GSE57973, N=5). **(C)** Gene expression in EndoC-$\beta$H1 cells treated with control siRNA at 72 hours (N=3). **(D)** Protein expression in DIA proteomics on cell lysate of EndoC-$\beta$H1 cells treated with control siRNA at 72 hours (N=12). **(E)** Protein expression in TMT proteomics on cell lysate of EndoC-$\beta$H1 cells treated with control siRNA at 72 hours (N=3). No boxplot for a protein in panels C and D indicates that the protein was either not detected or quantified in $<$ 3 samples.

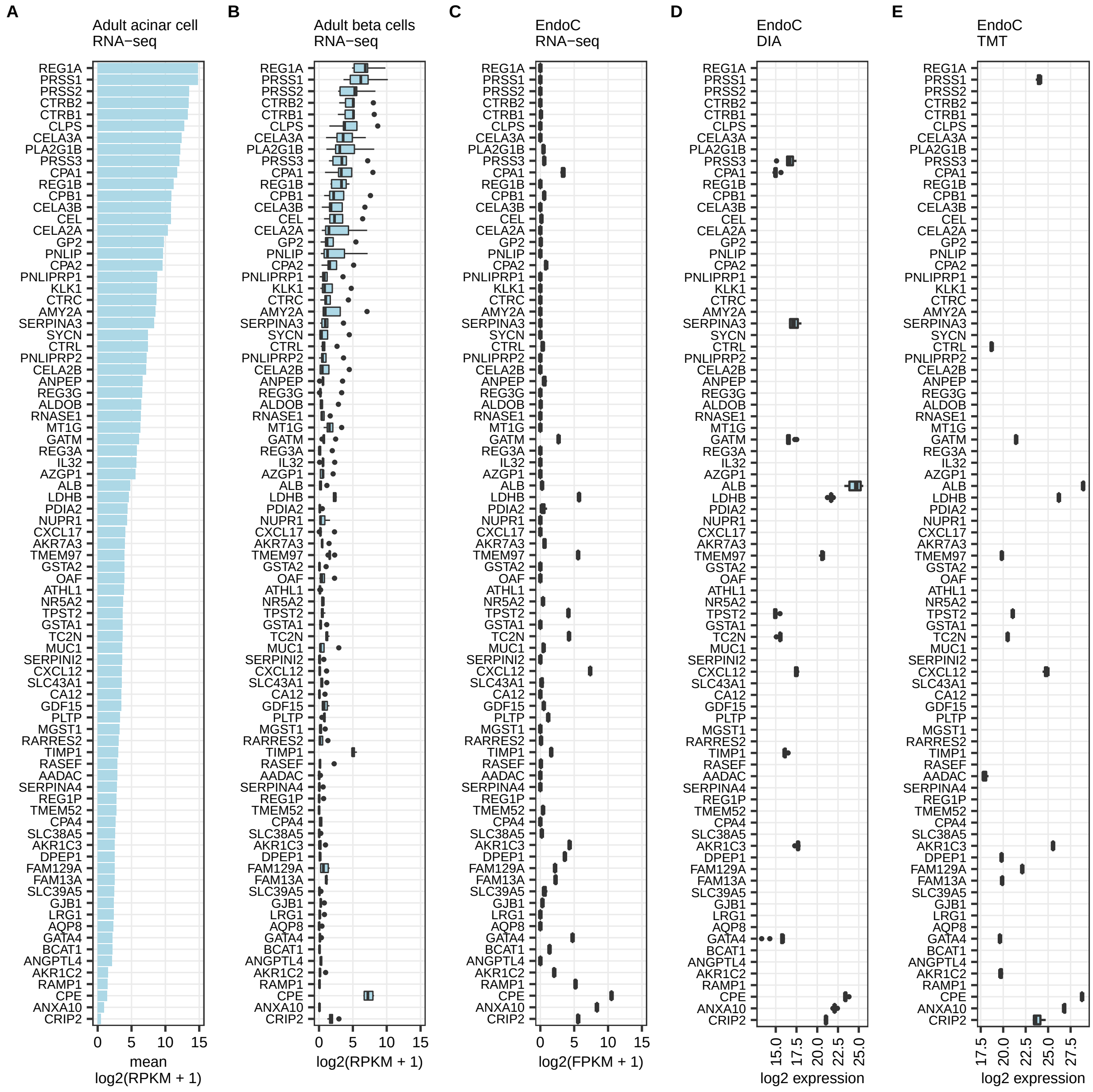

**Fig. S14. EndoC-$\beta$H1 cells did not express ductal cell markers. (A)** Gene expression in adult ductal cells (GSE57973, N= 2). **(B)** Gene expression in adult $\beta$ cells (GSE57973, N=5). **(C)** Gene expression in EndoC-$\beta$H1 cells treated with control siRNA at 72 hours (N=3). **(D)** Protein expression in DIA proteomics on cell lysate of EndoC-$\beta$H1 cells treated with control siRNA at 72 hours (N=12). **(E)** Protein expression in TMT proteomics on cell lysate of EndoC-$\beta$H1 cells treated with control siRNA at 72 hours (N=3). No boxplot for a protein in panels C and D indicates that the protein was either not detected or quantified in $<$ 3 samples.

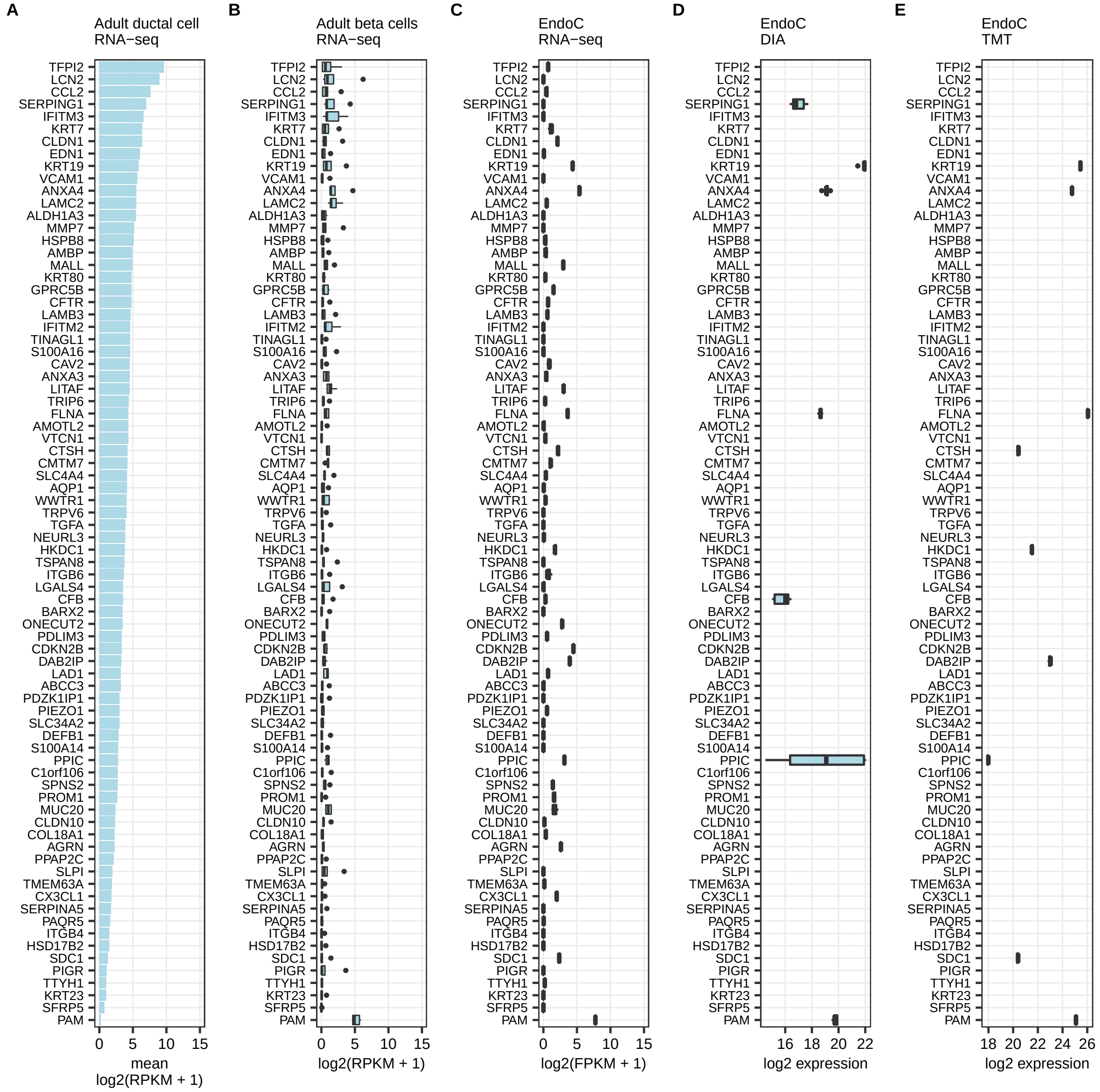

**Fig. S15. Ectopic expression of extra-islet tissue markers was not observed in EndoC-$\beta$H1 cells.** Tissue-specific genes (x=10) were extracted from supplementary data of Ryaboshapkina et al. 2019. Pancreas-specific genes were excluded from the analysis. **(A)** Median gene expression in the corresponding tissue with the highest expression in GTEx. N markers = 509. **(B)** Gene expression in adult $\beta$ cells (GSE67543, N=7, FACS sorted, at least 97% purity). **(C)** Gene expression in EndoC-$\beta$H1 cells treated with control siRNA at 72 hours (N=3). **(D)** Protein expression in DIA proteomics on cell lysate of EndoC-$\beta$H1 cells treated with control siRNA at 72 hours (N=12). **(E)** Protein expression in TMT proteomics on cell lysate of EndoC-$\beta$H1 cells treated with control siRNA at 72 hours (N=3). No boxplot for a protein in panels C and D indicates that the protein was either not detected or quantified in $<$ 3 samples.

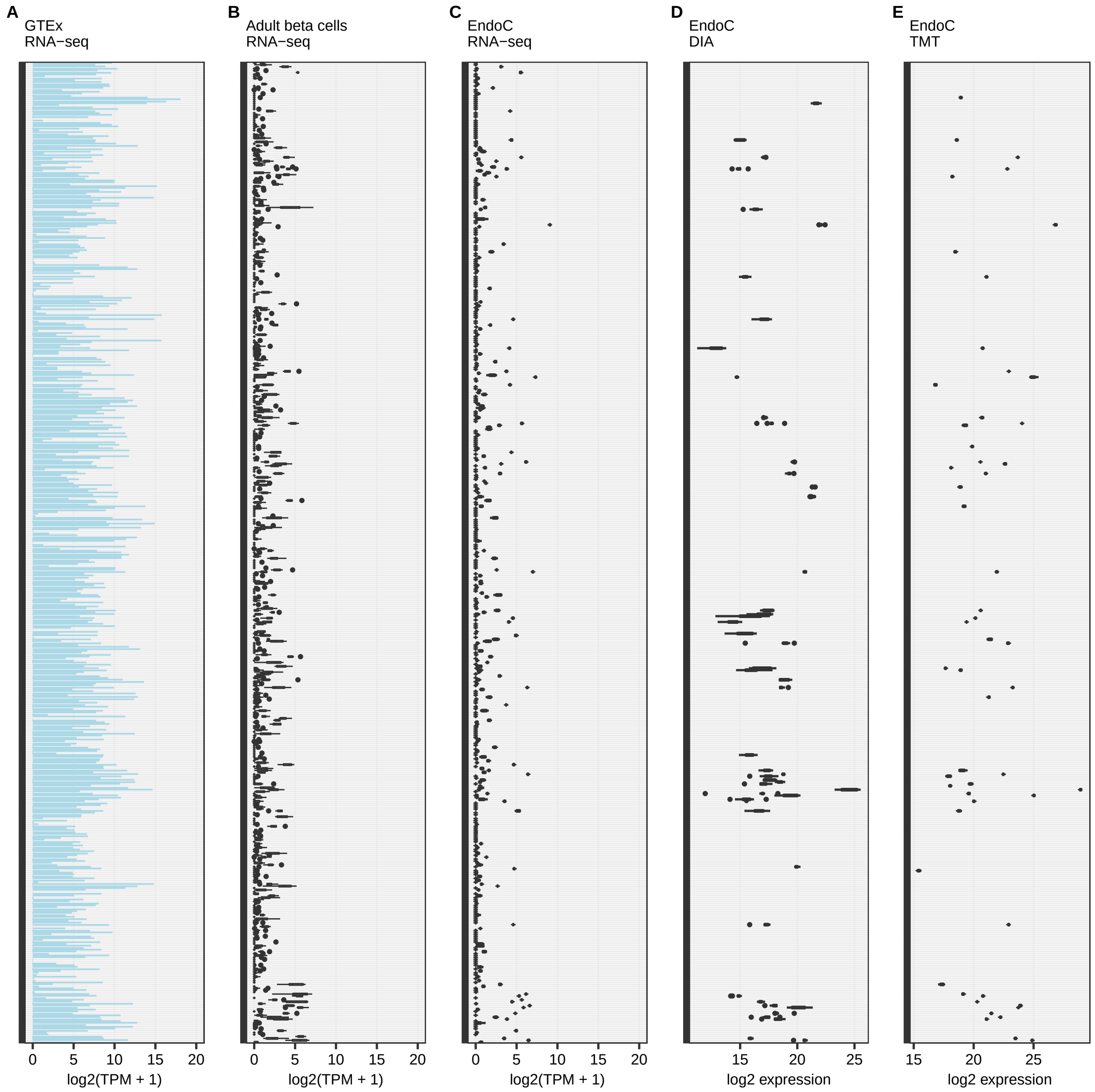

**Fig. S16. Potential chromogranin A bioactive peptides in secretome of EndoC-βH1 cells.** CLUSTAL O(1.2.4) multiple sequence alignment of bioactive peptide sequences (yellow) vs peptide sequences detected in the DIA secretome experiment at 72h. Alignment was performed online with UniProt (https://www.uniprot.org/align/).

### (A) EA-92

```
Reference EDSKEAEKSGEATDGARPQALPEPMQESKAEGNNQAPGEEEEEEEEATNTHPPASLPSQKYPGPQAEGDSEGLSQGLVDREKGLSAEPGWQA
Peptide 1 ------------------------------------------------------------YPGPQAEGDSEGLSQGLVDREK----------
Peptide 2 ------------------------------------------------------------YPGPQAEGDSEGLSQGLVDR------------
Peptide 3 -----------------------------AEGNNQAPGEEEEEEEEATNTHPPASLPSQK--------------------------------
Peptide 4 --------SGEATDGARPQALPEPMQESK---------------------------------------------------------------
Peptide 5 -----------------PQALPEPMQESK---------------------------------------------------------------
Peptide 6 ----EAEKSGEATDGARPQALPEPMQESK---------------------------------------------------------------
Peptide 7 -----------------------------AEGNNQAPGEEEEEEEEATNTHPPASLPSQKYPGPQAEGDSEGLSQGLVDR------------
Peptide 8 --------SGEATDGAR---------------------------------------------------------------------------
```

### (B) Pancreastatin

```
Reference SEALAVDGAGKPGAEEAQDPEGKGEQEHSQQKEEEEEMAVVPQGLFRG
Peptide 1 SEALAVDGAGK-------------------------------------
Peptide 2 SEALAVDGAGKPGAEEAQDPEGK-------------------------
Peptide 3 --------------------------------EEEEEMAVVPQGLFR-
Peptide 4 -----------PGAEEAQDPEGK-------------------------
Peptide 5 SEALAVDGAGKPGAEEAQDPEGKGEQEHSQQK----------------
Peptide 6 -----------------------GEQEHSQQKEEEEEMAVVPQGLFR-
Peptide 7 -----------PGAEEAQDPEGKGEQEHSQQK----------------
Peptide 8 -----------------------GEQEHSQQK----------------
```

### (C) SS-18

```
Reference SGELEQEEERLSKEWEDS
Peptide 1 SGELEQEEER--------
Peptide 2 SGELEQEEERLSK-----
```

### (D) LF-19

```
Reference LEGQEEEEDNRDSSMKLSF
Peptide 1 LEGQEEEEDNR--------
Peptide 2 LEGQEEEEDNRDSSMK---
```

### (E) GE-25

```
Reference GWRPSSREDSLEAGLPLQVRGYPEE
Peptide 1 -------EDSLEAGLPLQVR-----
Peptide 2 ---PSSREDSLEAGLPLQVR-----
```

### (E) Vasostatins

```
Vasostatin-2 LPVNSPMNKGDTEVMKCIVEVISDTLSKPSPMPVSQECFETLRGDERILSILRHQNLLKELQDLALQGAKERAHQQKKHSGFEDELSEVLENQSSQAELKEAVEEPSSKDVME
Vasostatin-1 LPVNSPMNKGDTEVMKCIVEVISDTLSKPSPMPVSQECFETLRGDERILSILRHQNLLKELQDLALQGAKERAHQQ-------------------------------------
Peptide 1    ------------------------------------------------------------------------------HSGFEDELSEVLENQSSQAELK-------------
Peptide 2    -----------------------------------------------------------ELQDLALQGAK-------------------------------------------
Peptide 3    ----------------------------------------------------------------------------------------------------EAVEEPSSK----
Peptide 4    ------------------------------------------------------------------------------HSGFEDELSEVLENQSSQAELKEAVEEPSSK----
Peptide 5    -----------------------------------------------------------------------------KHSGFEDELSEVLENQSSQAELK-------------
Peptide 6    -----------------------------------------------------------ELQDLALQGAKER-----------------------------------------
Peptide 7    ---------GDTEVMK-------------------------------------------------------------------------------------------------
Peptide 8    -----------------------------------------------------HQNLLKELQDLALQGAK-------------------------------------------
```

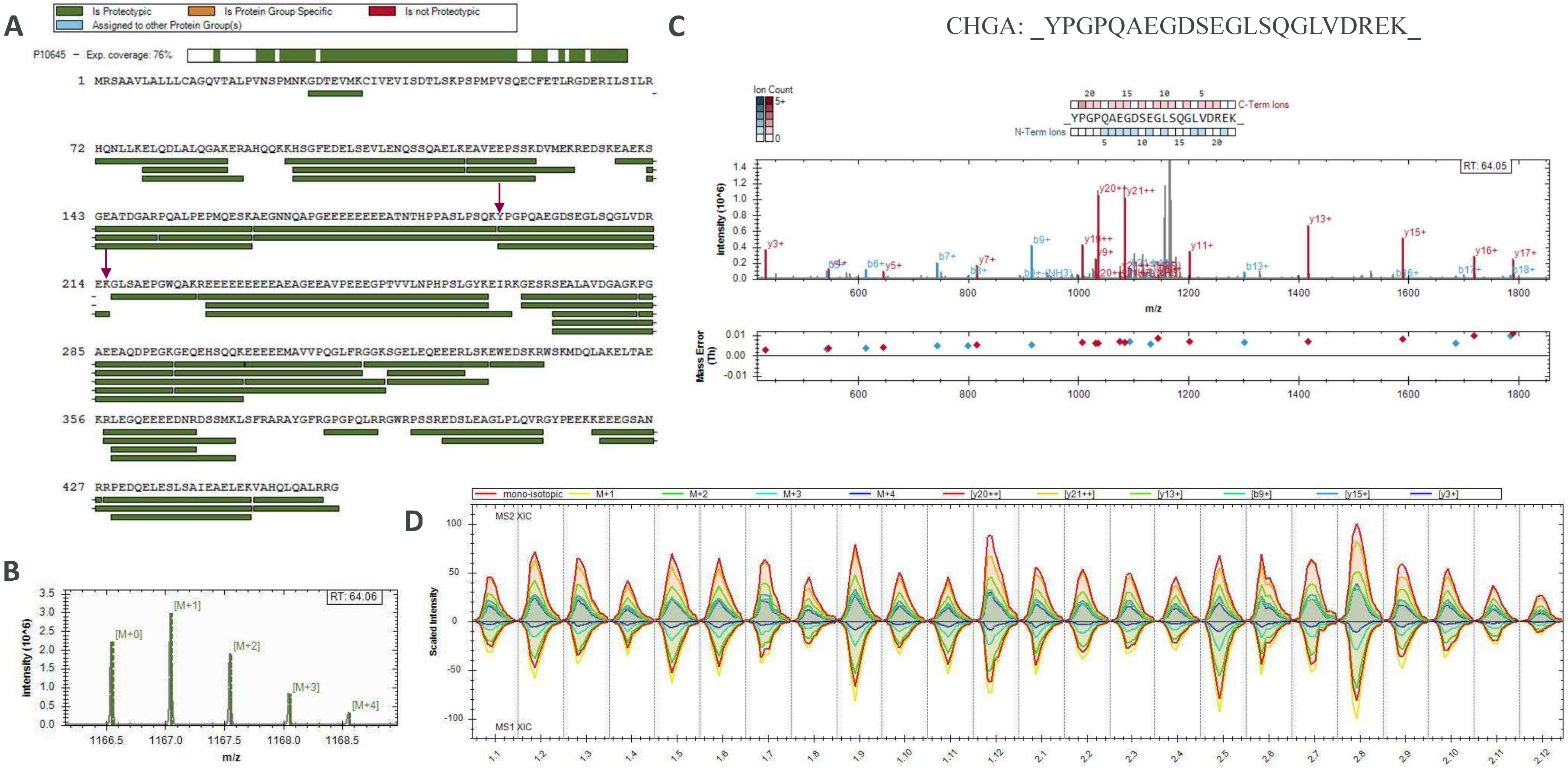


**Fig. S17. Chromogranin A proteotypic peptides identified by DIA secretome proteomics experiment at 72h.** Total protein coverage (**A**), MS1 isotopic envelope (**B**), MS/MS fragmentation match and mass error (**C**), and XIC of MS1 & MS2 for the conditions (**D**) are shown for each given peptide identified.

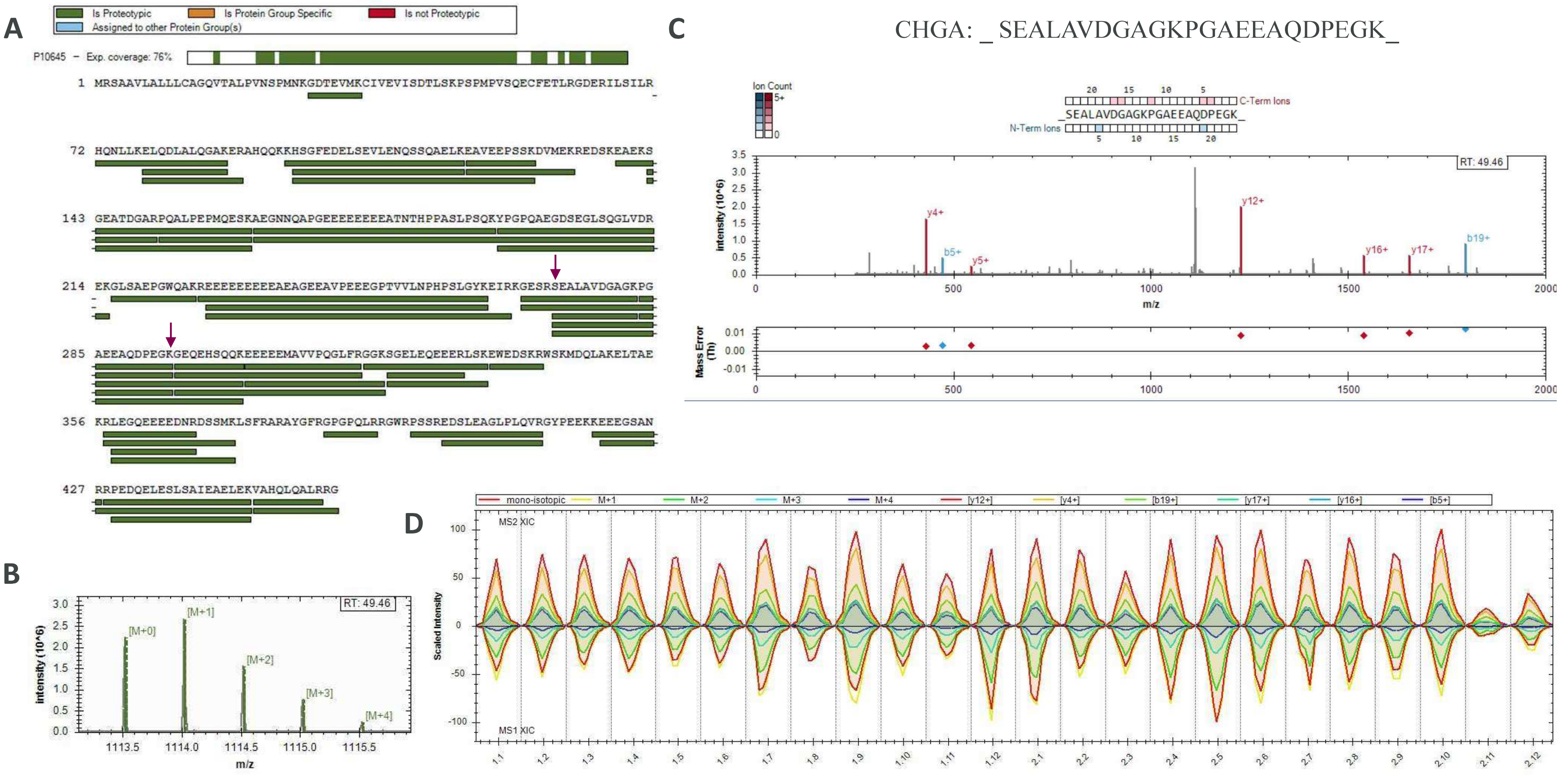


**Fig. S18. Chromogranin A proteotypic peptides identified by DIA secretome proteomics experiment at 72h.** Total protein coverage (**A**), MS1 isotopic envelope (**B**), MS/MS fragmentation match and mass error (**C**), and XIC of MS1 & MS2 for the conditions (**D**) are shown for each given peptide identified.

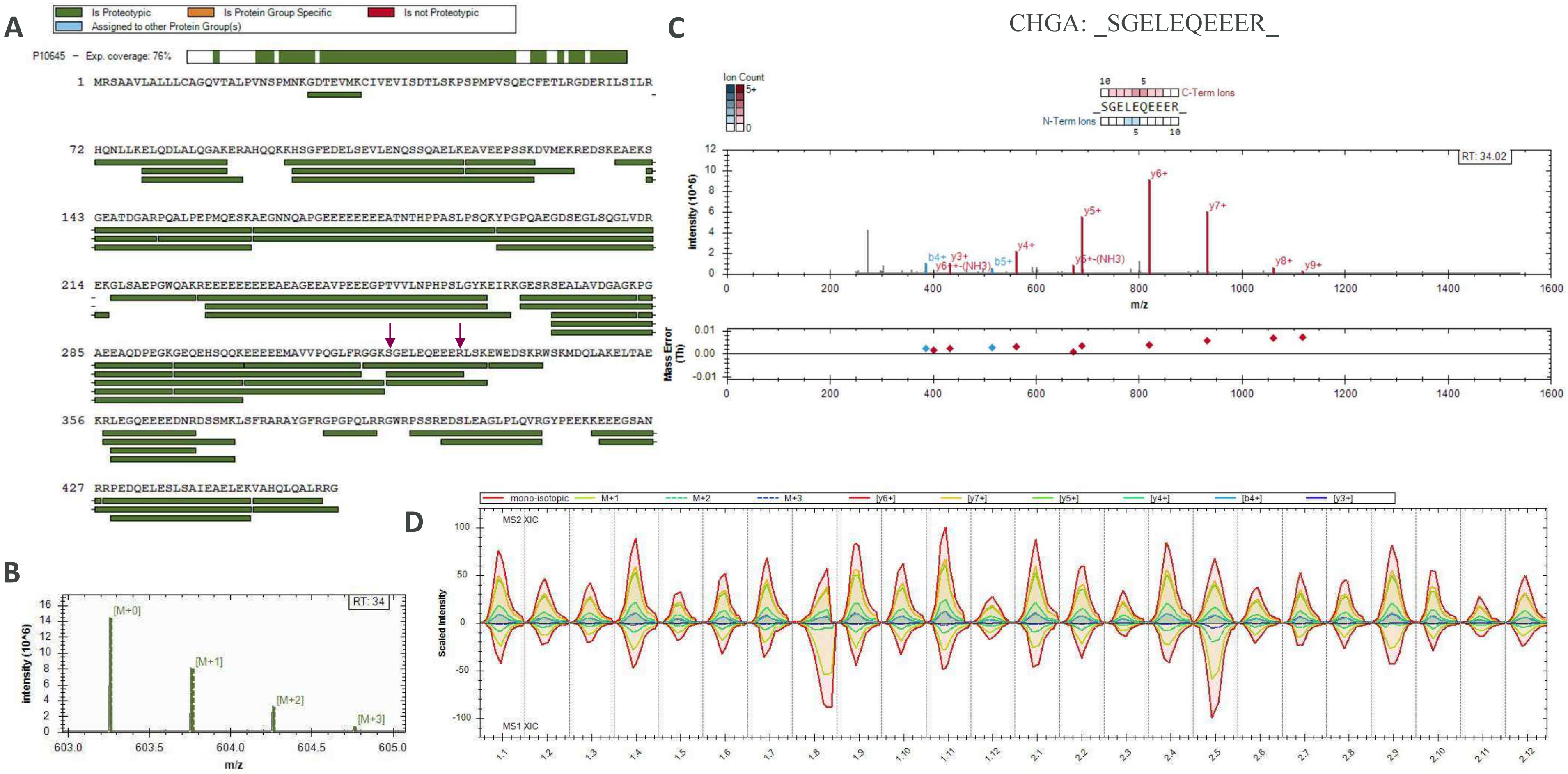


**Fig. S19. Chromogranin A proteotypic peptides identified by DIA secretome proteomics experiment at 72h.** Total protein coverage (**A**), MS1 isotopic envelope (**B**), MS/MS fragmentation match and mass error (**C**), and XIC of MS1 & MS2 for the conditions (**D**) are shown for each given peptide identified.

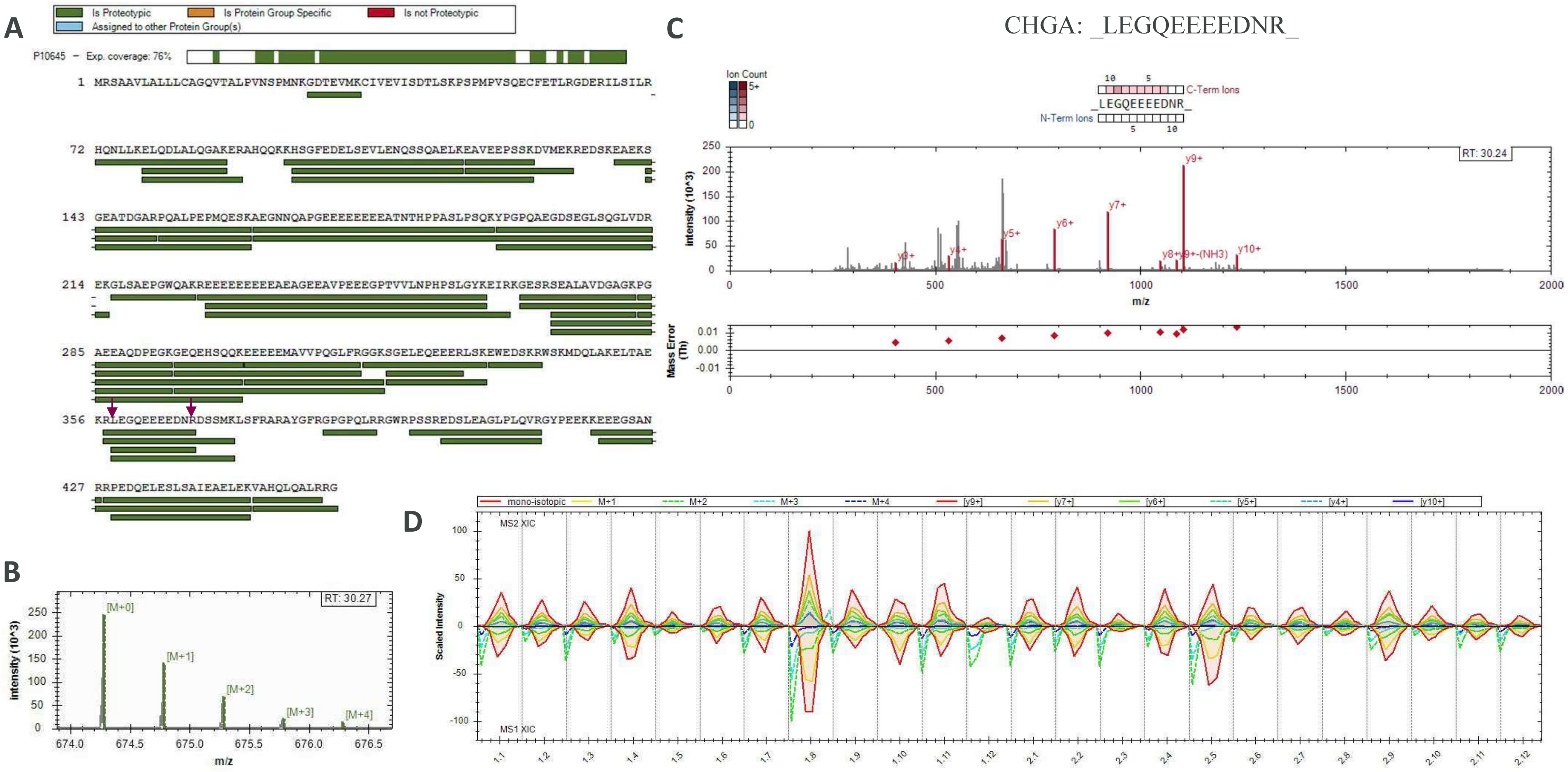


**Fig. S20. Chromogranin A proteotypic peptides identified by DIA secretome proteomics experiment at 72h.** Total protein coverage (**A**), MS1 isotopic envelope (**B**), MS/MS fragmentation match and mass error (**C**), and XIC of MS1 & MS2 for the conditions (**D**) are shown for each given peptide identified.

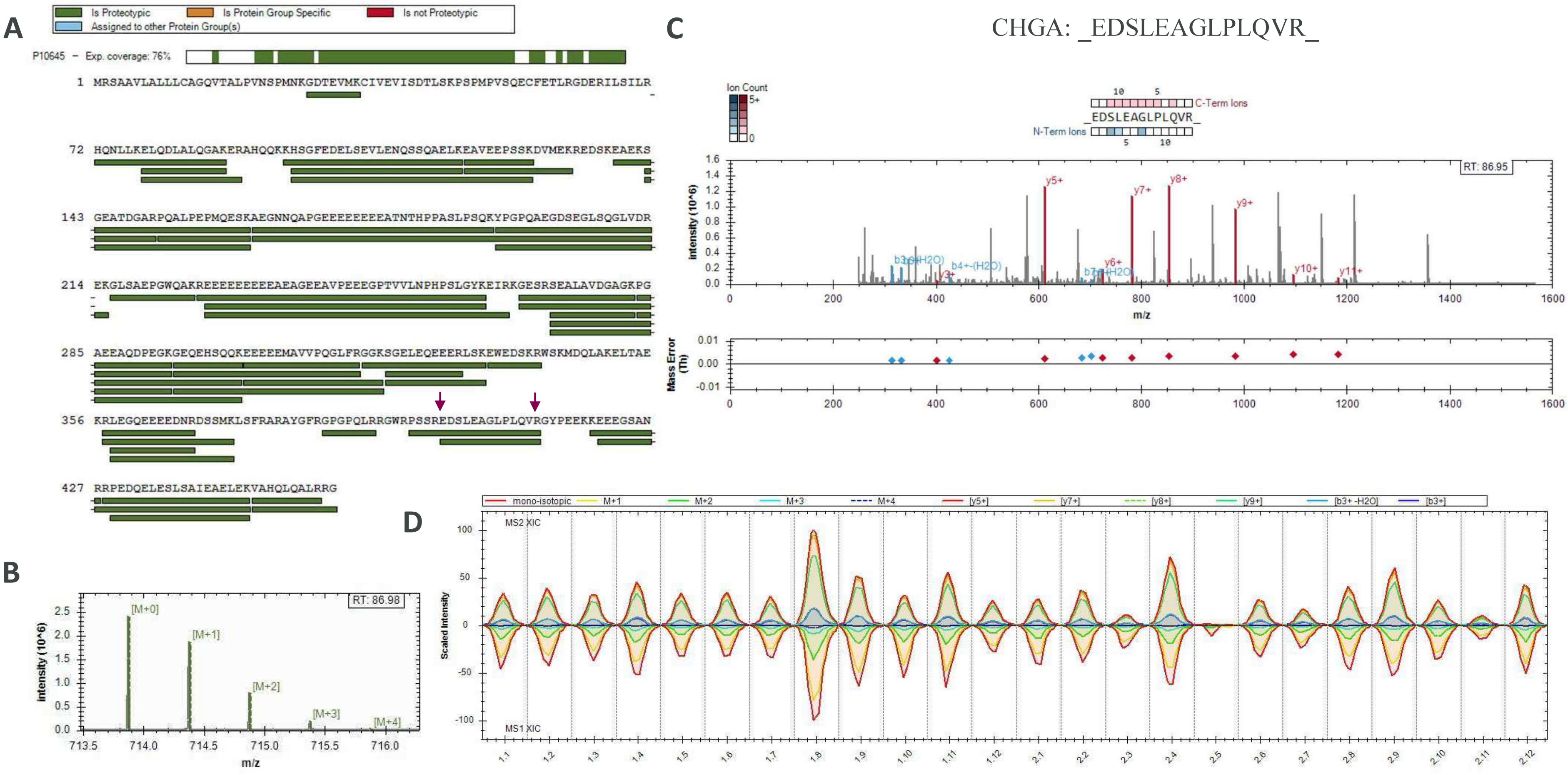


**Fig. S21. Chromogranin A proteotypic peptides identified by DIA secretome proteomics experiment at 72h.** Total protein coverage (**A**), MS1 isotopic envelope (**B**), MS/MS fragmentation match and mass error (**C**), and XIC of MS1 & MS2 for the conditions (**D**) are shown for each given peptide identified.

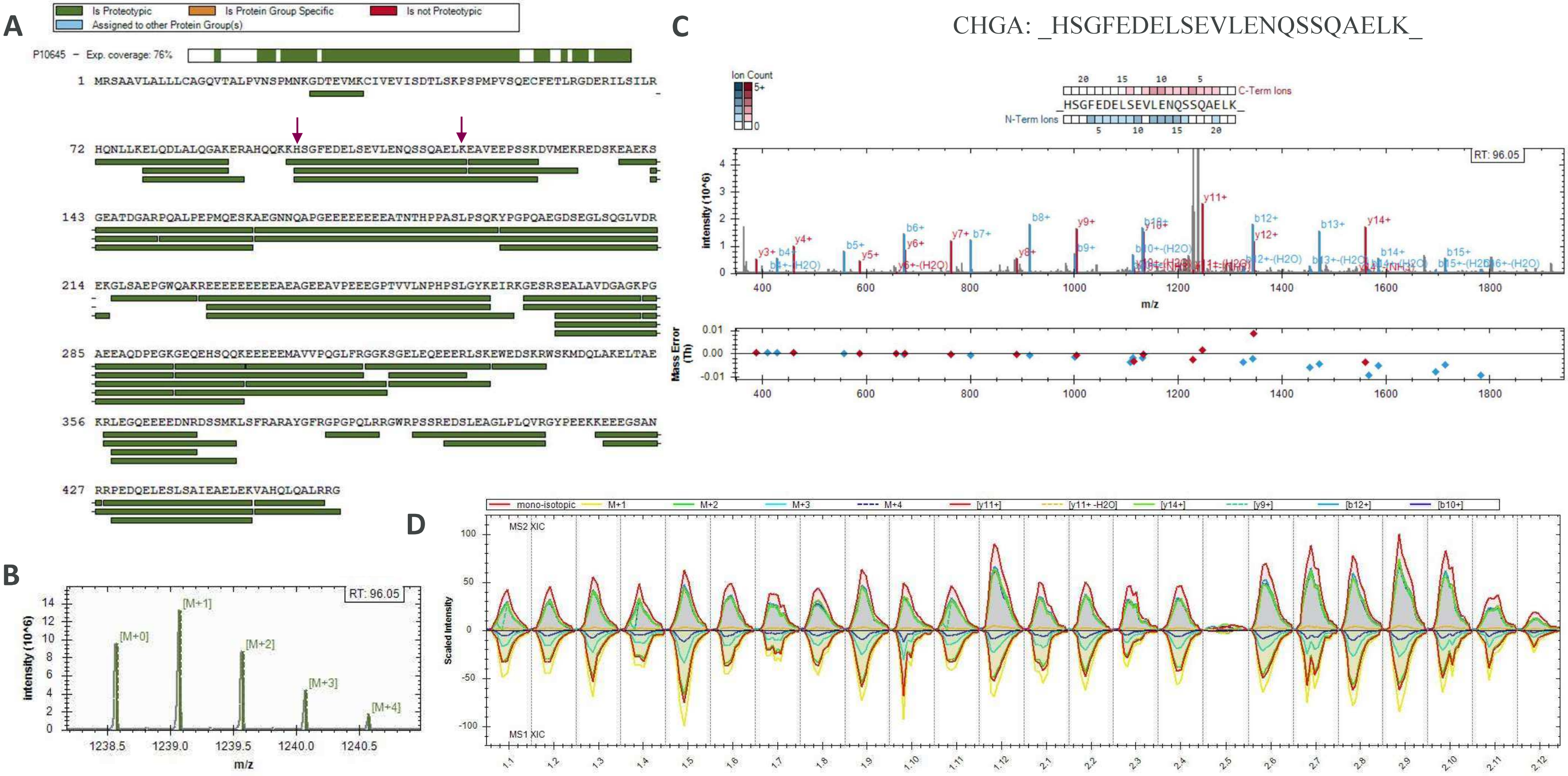


**Fig. S22. Chromogranin A proteotypic peptides identified by DIA secretome proteomics experiment at 72h.** Total protein coverage (**A**), MS1 isotopic envelope (**B**), MS/MS fragmentation match and mass error (**C**), and XIC of MS1 & MS2 for the conditions (**D**) are shown for each given peptide identified.

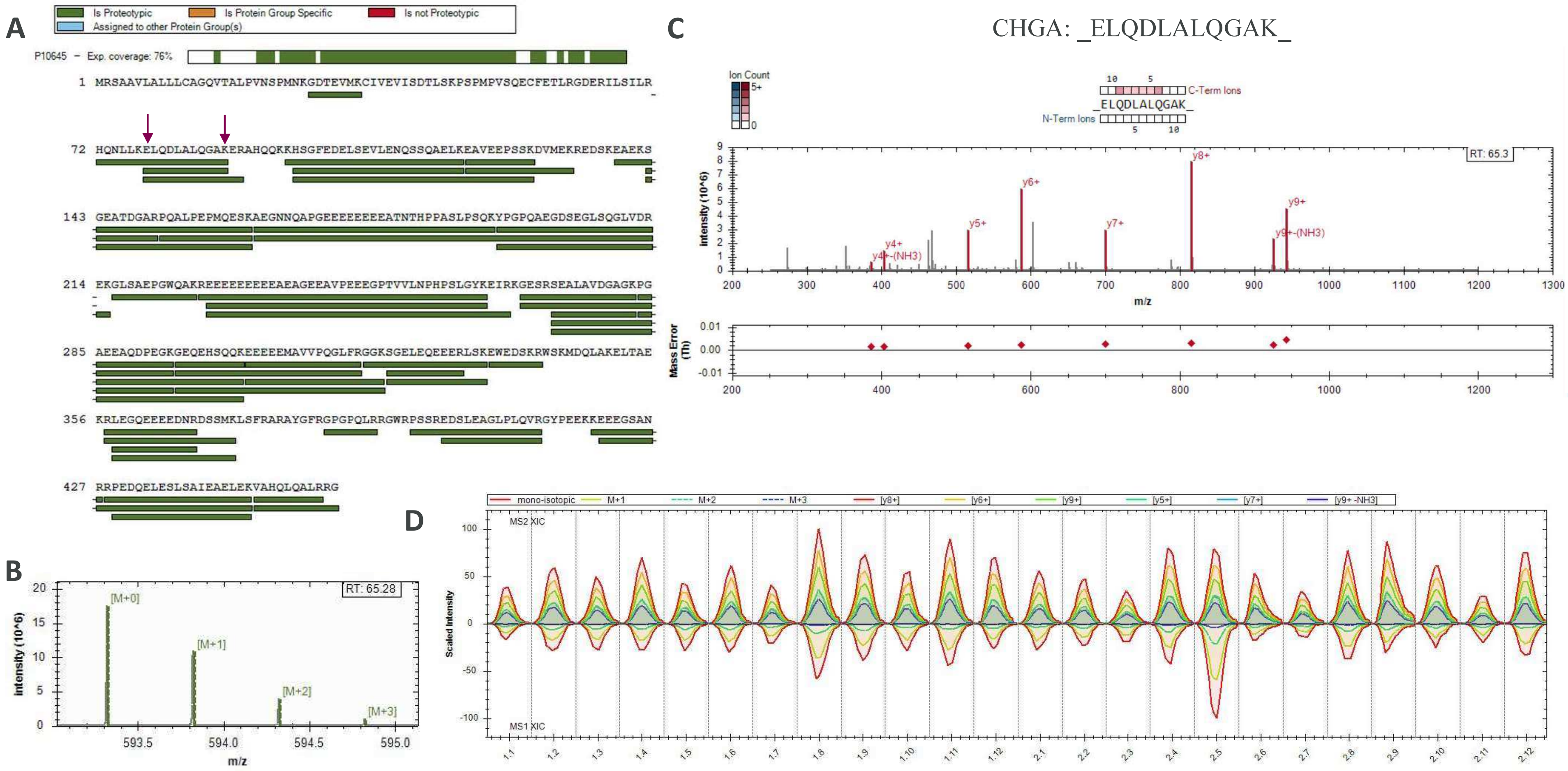


**Fig. S23. Chromogranin A proteotypic peptides identified by DIA secretome proteomics experiment at 72h.** Total protein coverage (**A**), MS1 isotopic envelope (**B**), MS/MS fragmentation match and mass error (**C**), and XIC of MS1 & MS2 for the conditions (**D**) are shown for each given peptide identified.

**Fig. S24. Potential chromogranin B bioactive peptides in secretome of EndoC-βH1 cells.** CLUSTAL O(1.2.4) multiple sequence alignment of bioactive peptide sequences (yellow) vs peptide sequences detected in the DIA secretome experiment at 72h. Alignment was performed online with UniProt (https://www.uniprot.org/align/).

**(A) CCB peptide**

```
Reference SAEFPDFYDSEEPVSTHQEAENEKDRADQTVLTEDEKKELENLAAMDLELQKIAEKF
Peptide 1 ------------------------DRADQTVLTEDEK--------------------
Peptide 2 --------------------------ADQTVLTEDEK--------------------
Peptide 3 --------------------------------------ELENLAAMDLELQK-----
Peptide 4 ------------------------DRADQTVLTEDEKK-------------------
Peptide 5 SAEFPDFYDSEEPVSTHQEAENEK---------------------------------
```

**(B) GAWK peptide**

```
Reference FLGEGHHRVQENQMDKARRHPQGAWKELDRNYLNYGEEGAPGKWQQQGDLQDTKENREEARFQDKQYSSHHTAE
Peptide 1 -------------------------------------------WQQQGDLQDTK--------------------
Peptide 2 ------------------------------NYLNYGEEGAPGK-------------------------------
Peptide 3 --------------------------ELDRNYLNYGEEGAPGK-------------------------------
Peptide 4 FLGEGHHR------------------------------------------------------------------
```

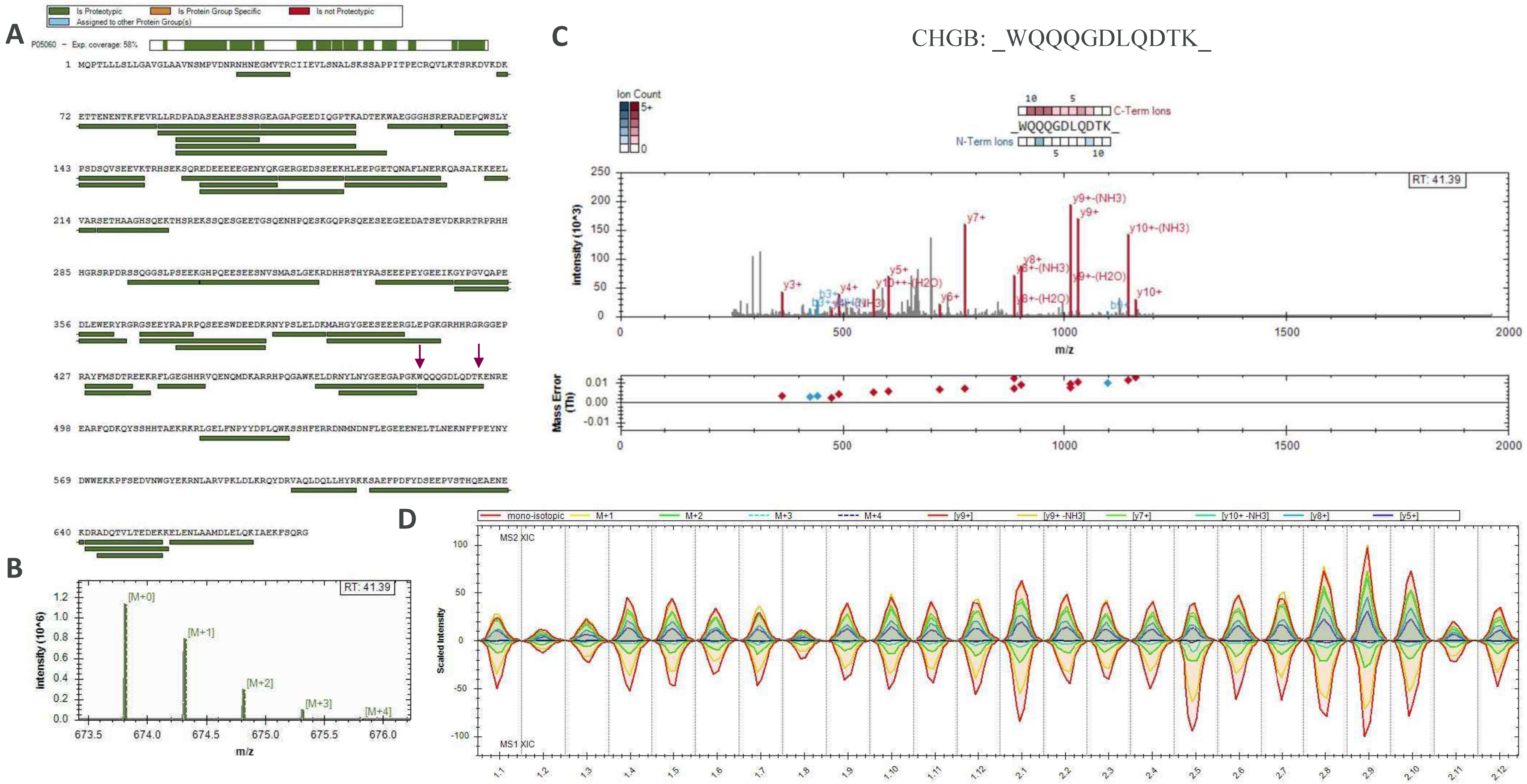


**Fig. S25. Chromogranin B proteotypic peptides identified by DIA secretome proteomics experiment at 72h.** Total protein coverage (**A**), MS1 isotopic envelope (**B**), MS/MS fragmentation match and mass error (**C**), and XIC of MS1 & MS2 for the conditions (**D**) are shown for each given peptide identified.

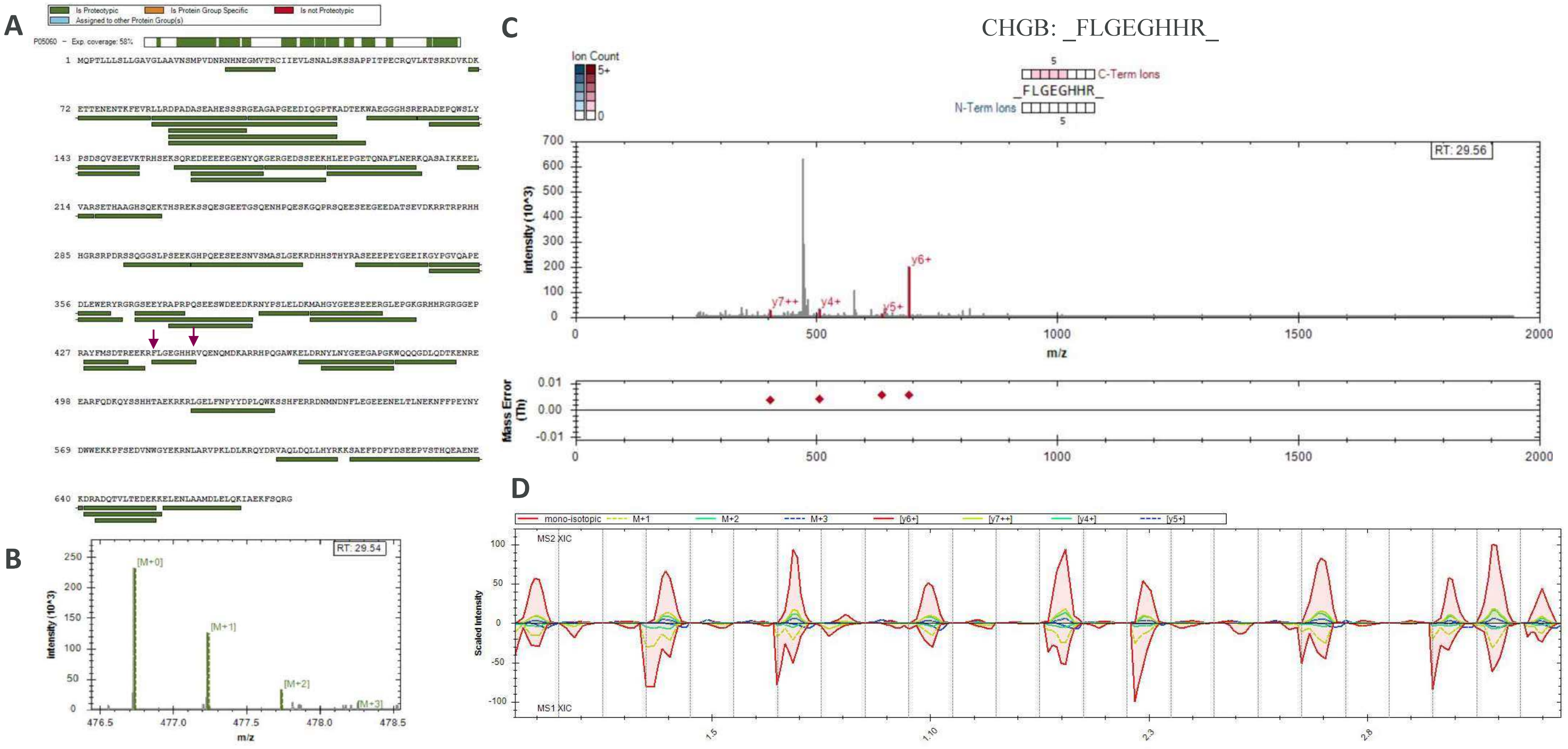


**Fig. S26. Chromogranin B proteotypic peptides identified by DIA secretome proteomics experiment at 72h.** Total protein coverage (**A**), MS1 isotopic envelope (**B**), MS/MS fragmentation match and mass error (**C**), and XIC of MS1 & MS2 for the conditions (**D**) are shown for each given peptide identified.

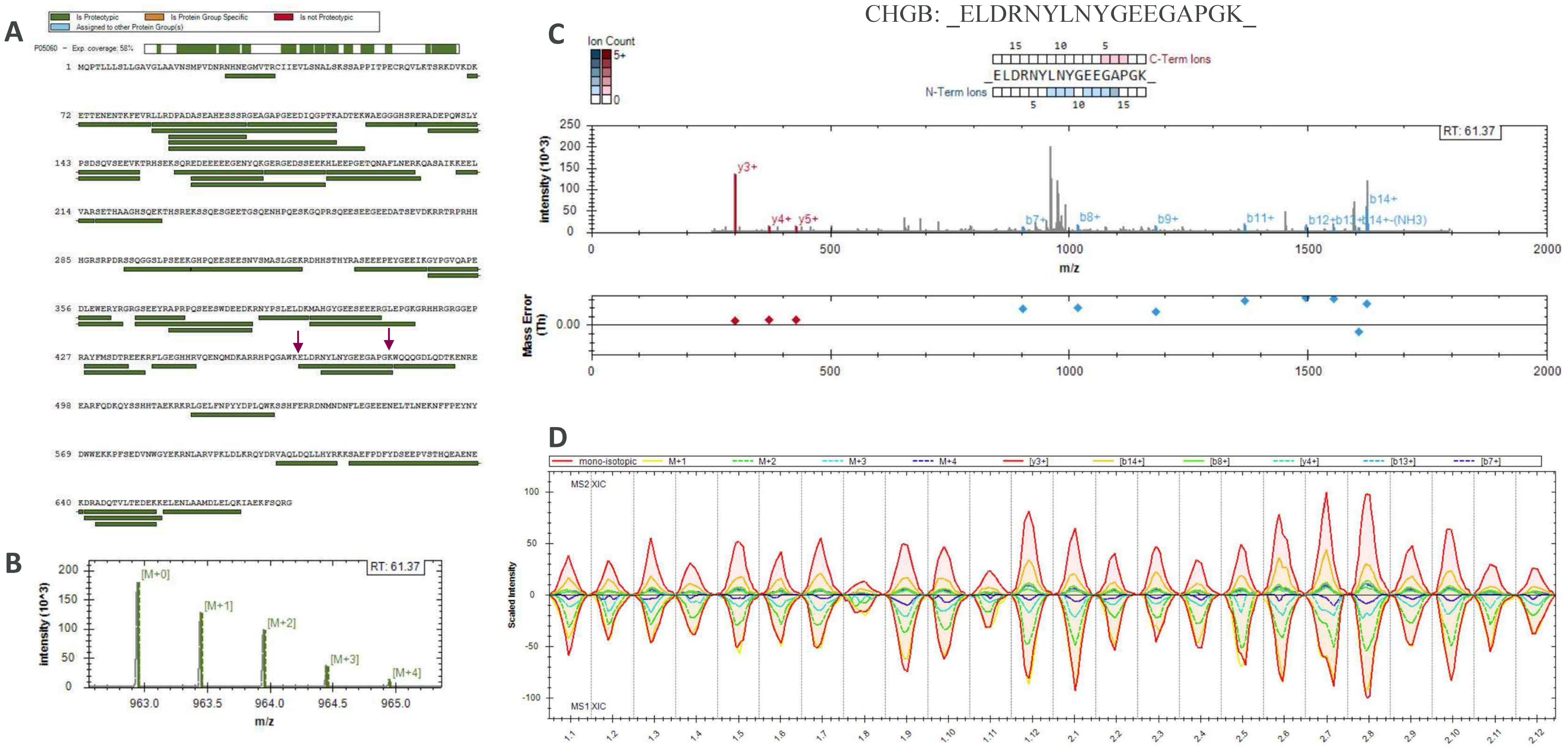


**Fig. S27. Chromogranin B proteotypic peptides identified by DIA secretome proteomics experiment at 72h.** Total protein coverage (**A**), MS1 isotopic envelope (**B**), MS/MS fragmentation match and mass error (**C**), and XIC of MS1 & MS2 for the conditions (**D**) are shown for each given peptide identified.

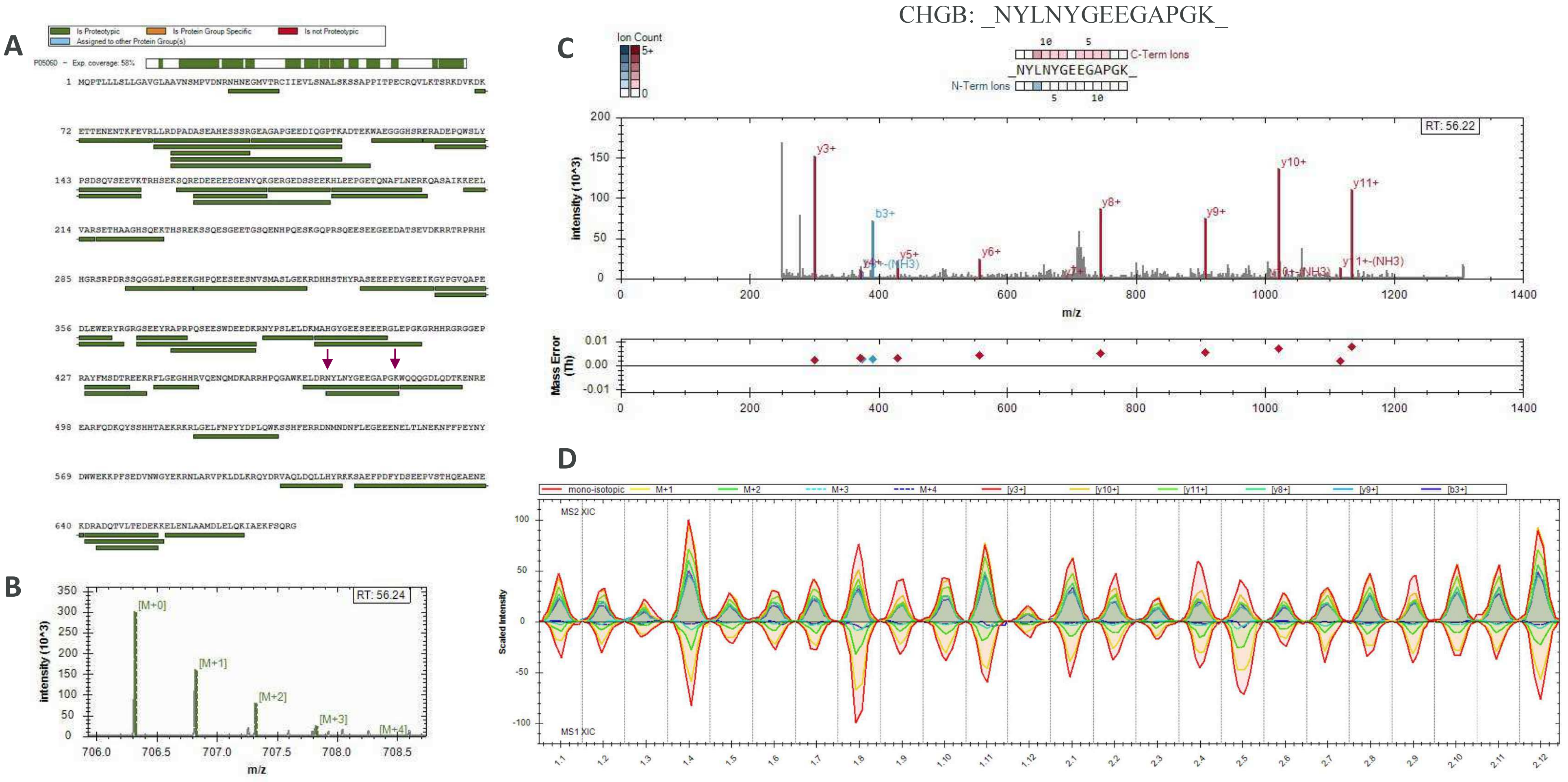


**Fig. S28. Chromogranin B proteotypic peptides identified by DIA secretome proteomics experiment at 72h.** Total protein coverage (**A**), MS1 isotopic envelope (**B**), MS/MS fragmentation match and mass error (**C**), and XIC of MS1 & MS2 for the conditions (**D**) are shown for each given peptide identified.

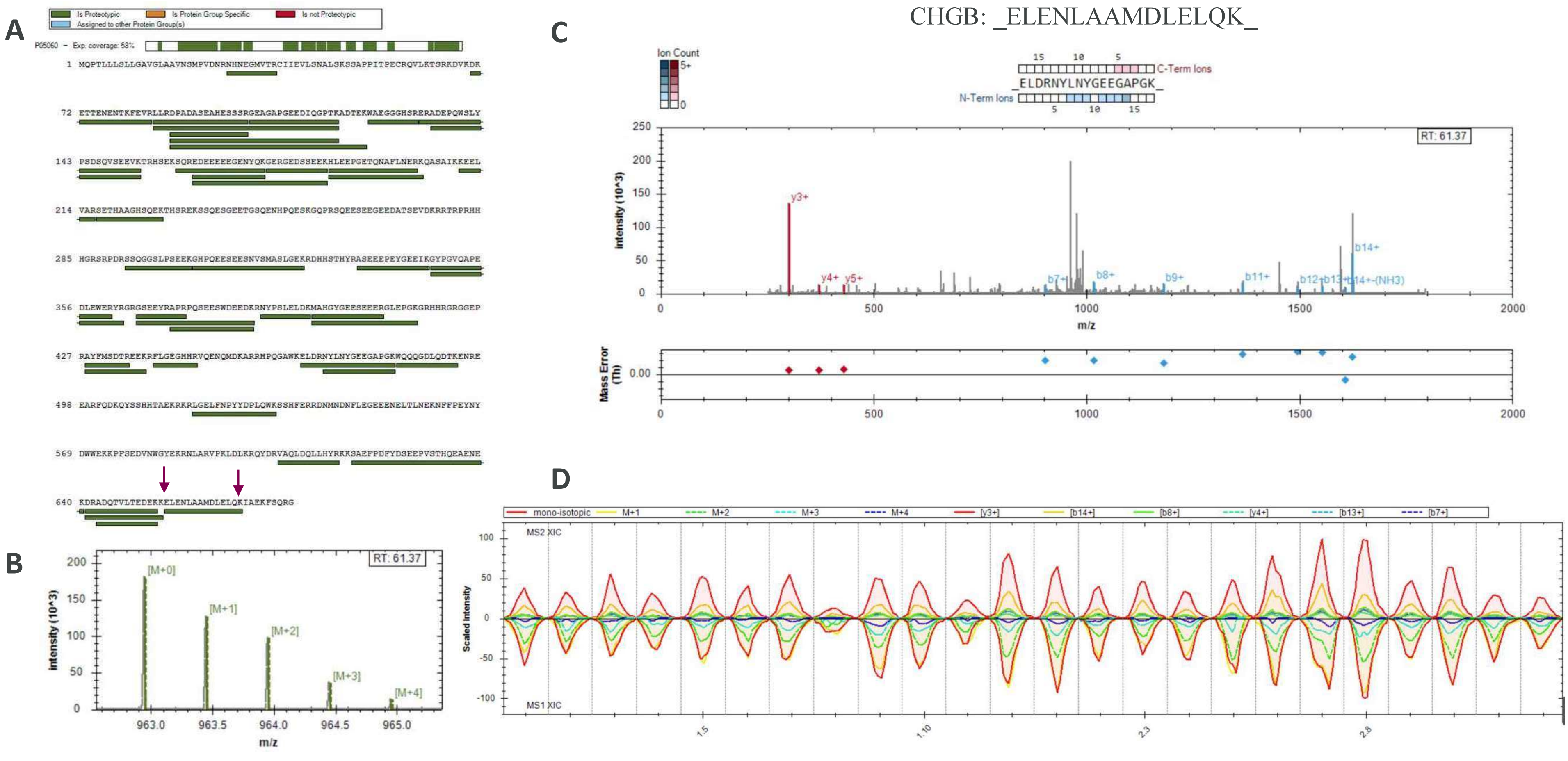


**Fig. S29. Chromogranin B proteotypic peptides identified by DIA secretome proteomics experiment at 72h.** Total protein coverage (**A**), MS1 isotopic envelope (**B**), MS/MS fragmentation match and mass error (**C**), and XIC of MS1 & MS2 for the conditions (**D**) are shown for each given peptide identified.

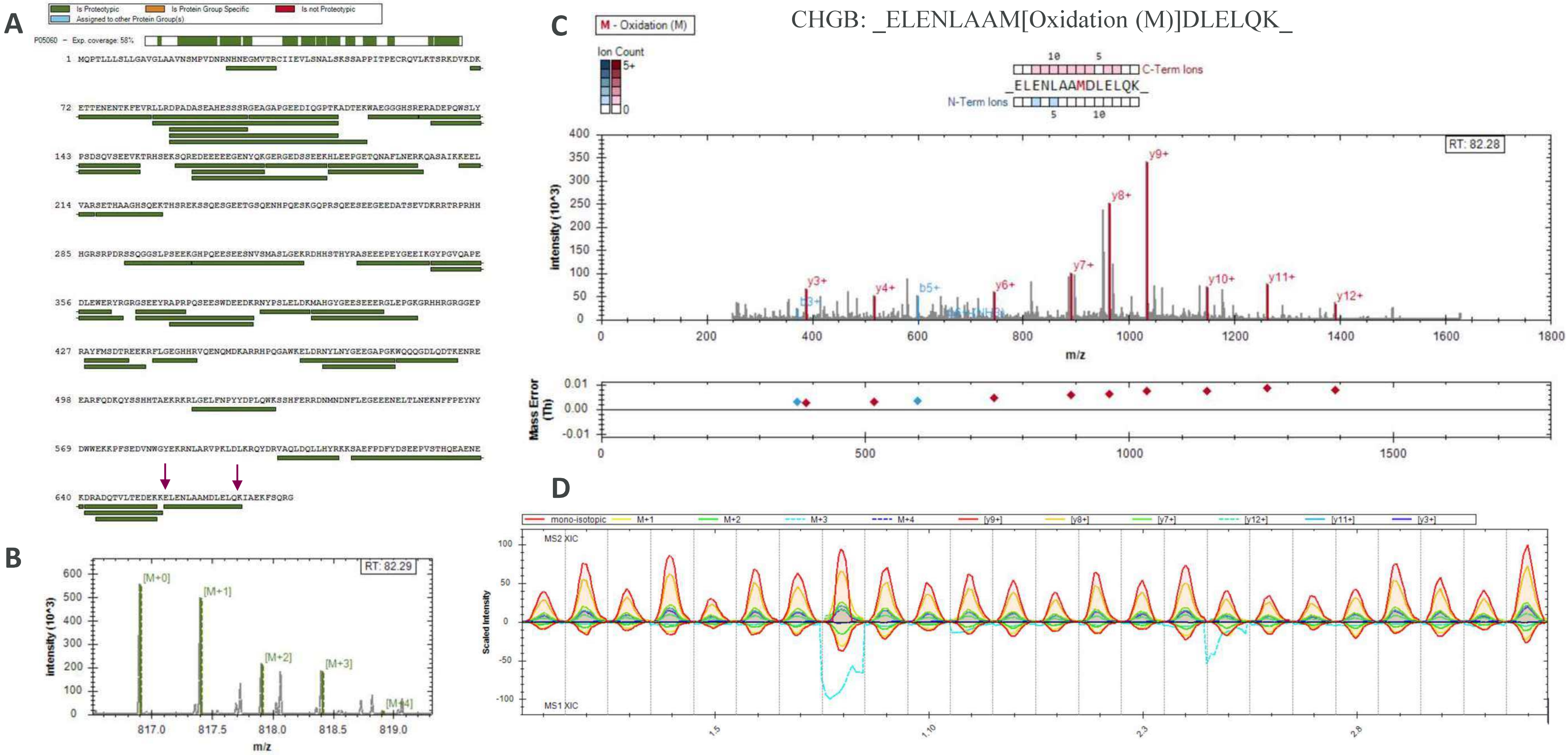


**Fig. S30. Chromogranin B proteotypic peptides identified by DIA secretome proteomics experiment at 72h.** Total protein coverage (**A**), MS1 isotopic envelope (**B**), MS/MS fragmentation match and mass error (**C**), and XIC of MS1 & MS2 for the conditions (**D**) are shown for each given peptide identified.

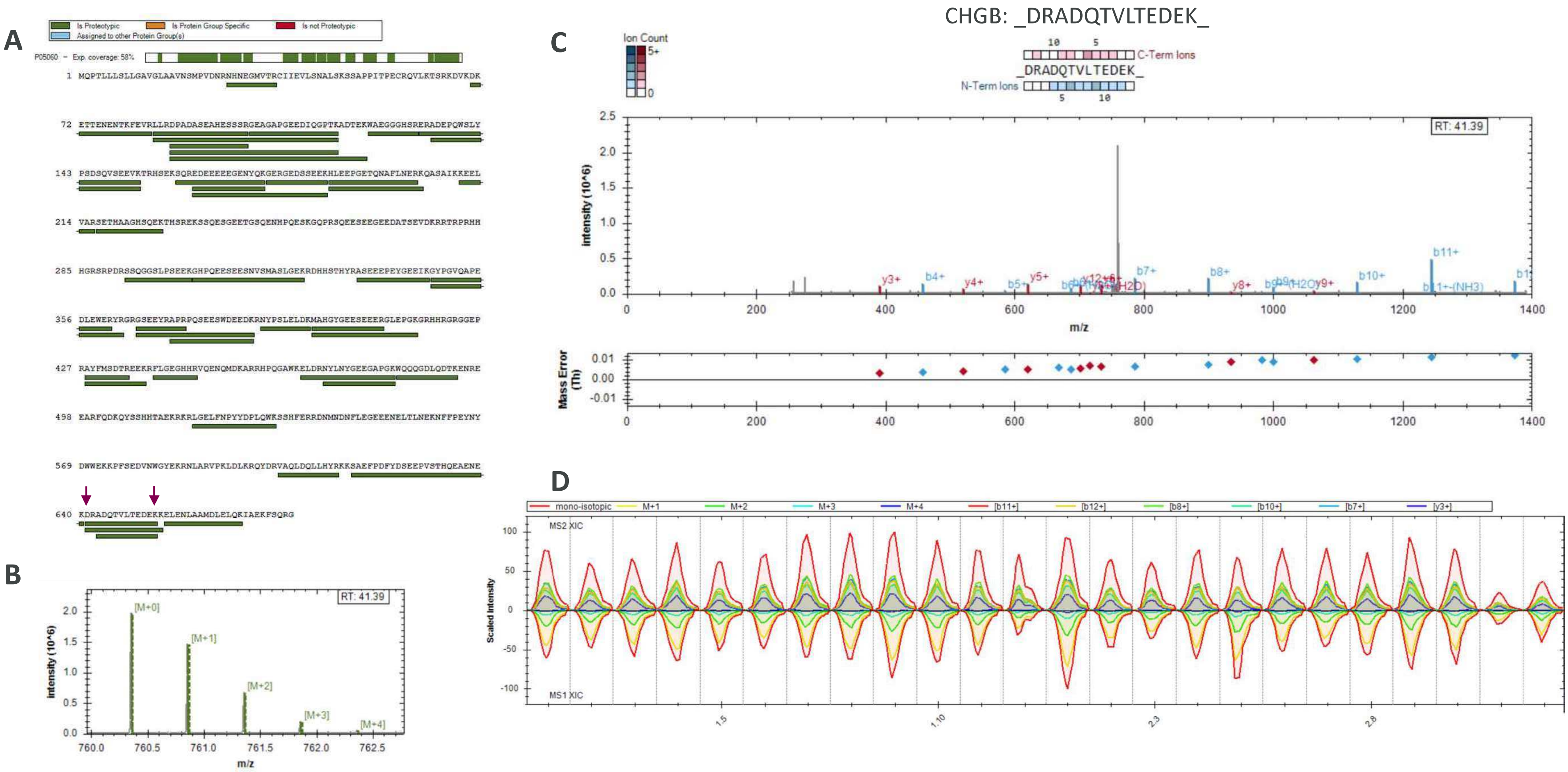


**Fig. S31. Chromogranin B proteotypic peptides identified by DIA secretome proteomics experiment at 72h.** Total protein coverage (**A**), MS1 isotopic envelope (**B**), MS/MS fragmentation match and mass error (**C**), and XIC of MS1 & MS2 for the conditions (**D**) are shown for each given peptide identified.

**Fig. S32. Potential VGF bioactive peptides in secretome of EndoC-βH1 cells.** CLUSTAL O(1.2.4) multiple sequence alignment of bioactive peptide sequences (yellow) vs peptide sequences detected in the DIA secretome experiment at 72h. Alignment was performed online with UniProt (https://www.uniprot.org/align/).

### (A) Neuroendocrine regulatory peptide-1

```
Reference RPESALLGGSEAGERLLQQGLAQVEA
Peptide 1 RPESALLGGSEAGER-----------
Peptide 2 -PESALLGGSEAGER-----------
```

### (B) Neuroendocrine regulatory peptide-2

```
Reference QAEATRQAAAQEERLADLASDLLLQYLLQGGARQRGLG
Peptide 1 --------------LADLASDLLLQYLLQGGAR-----
```

### (C) TLQP-62

```
Reference TLQPPSALRRRHYHHALPPSRHYPGREAQARRAQEEAEAEERRLQEQEELENYIEHVLLRRP
Peptide 1 ------------------------------------------RLQEQEELENYIEHVLLR--
Peptide 2 -------------------------------RAQEEAEAEER--------------------
Peptide 3 --------------------------------AQEEAEAEER--------------------
Peptide 4 -------------------------------------------LQEQEELENYIEHVLLR--
Peptide 5 --------------------------------AQEEAEAEERR-------------------
Peptide 6 -------------------------------------------LQEQEELENYIEHVLLRR-
Peptide 7 -------------------------------RAQEEAEAEERR-------------------
```

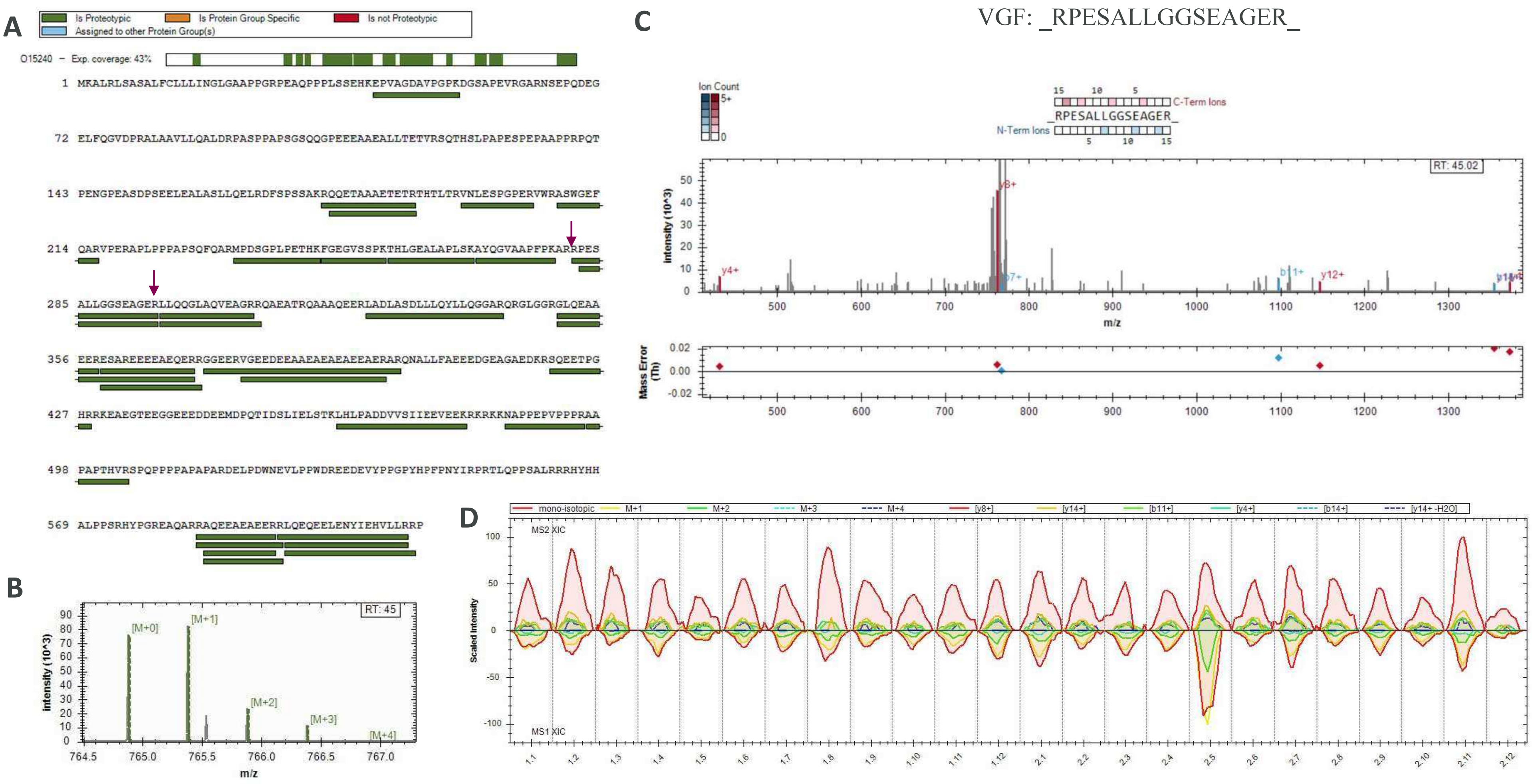


**Fig. S33. VGF proteotypic peptides identified by DIA secretome proteomics experiment at 72h.** Total protein coverage (**A**), MS1 isotopic envelope (**B**), MS/MS fragmentation match and mass error (**C**), and XIC of MS1 & MS2 for the conditions (**D**) are shown for each given peptide identified.

VGF: _LADLASDLLLQYLLQGGAR_

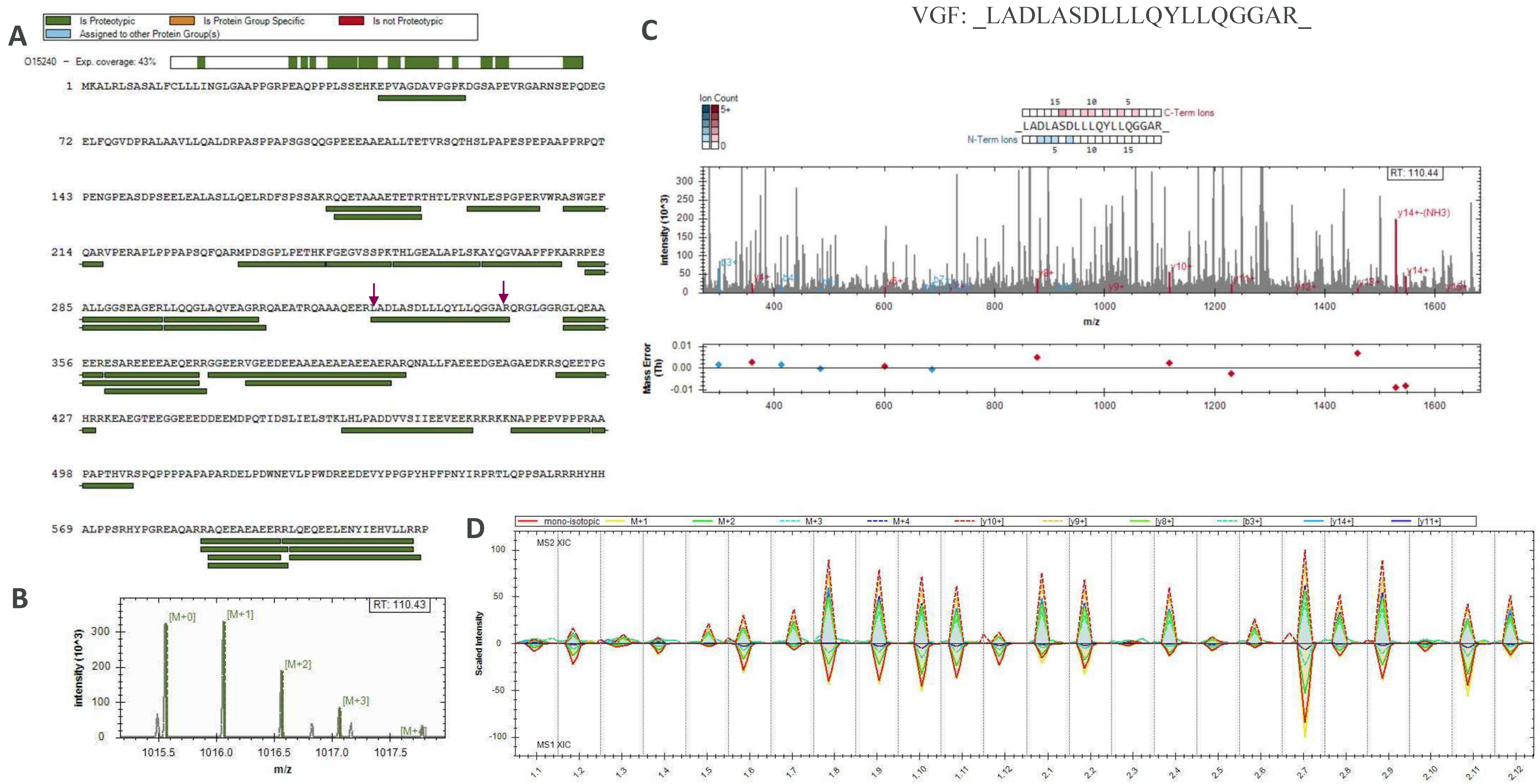


**Fig. S34. VGF proteotypic peptides identified by DIA secretome proteomics experiment at 72h.** Total protein coverage (**A**), MS1 isotopic envelope (**B**), MS/MS fragmentation match and mass error (**C**), and XIC of MS1 & MS2 for the conditions (**D**) are shown for each given peptide identified.

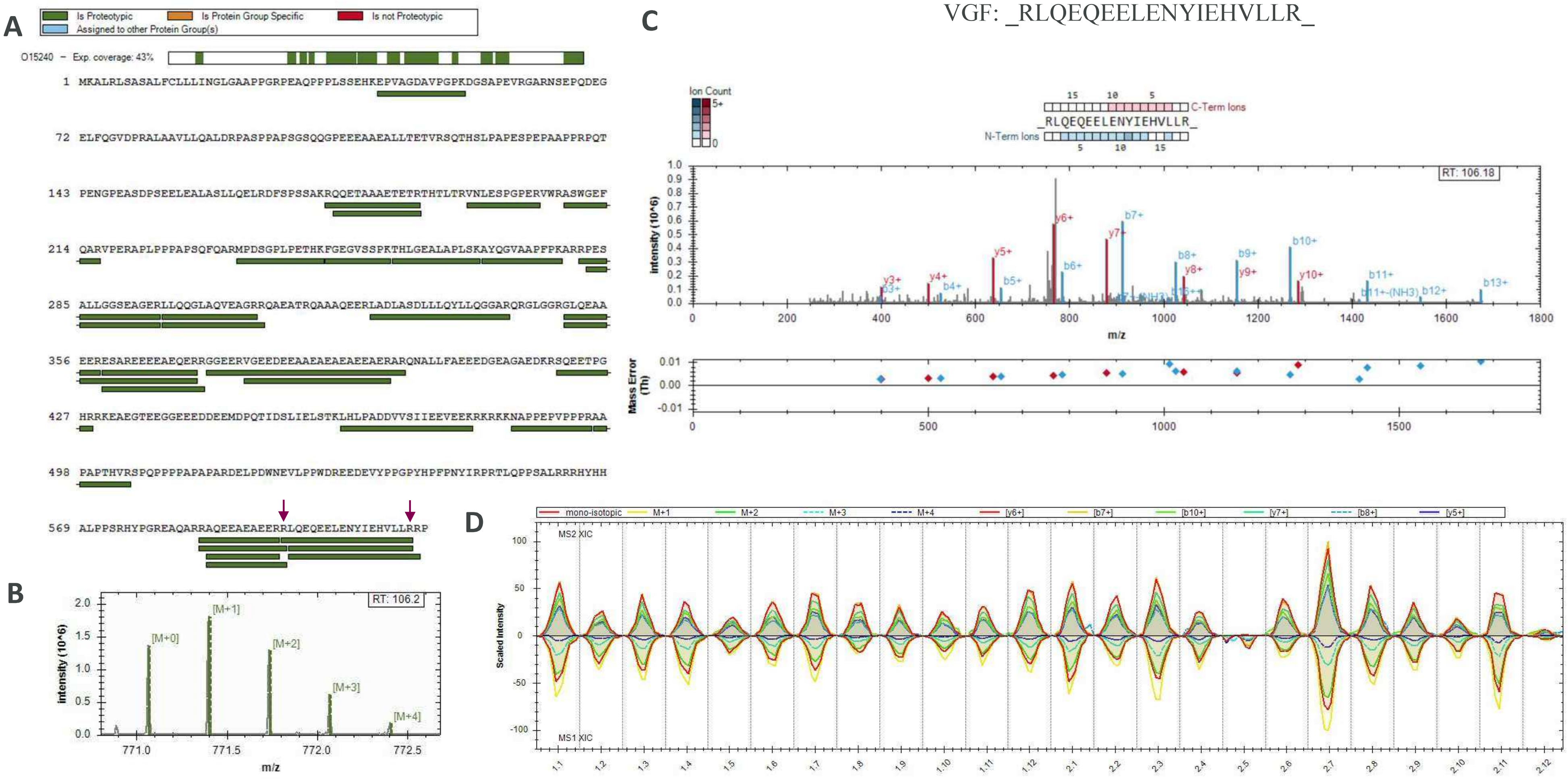


**Fig. S35. VGF proteotypic peptides identified by DIA secretome proteomics experiment at 72h.** Total protein coverage (**A**), MS1 isotopic envelope (**B**), MS/MS fragmentation match and mass error (**C**), and XIC of MS1 & MS2 for the conditions (**D**) are shown for each given peptide identified.

**Fig. S36. Potential bioactive peptides from other proteins in secretome of EndoC-βH1 cells.** CLUSTAL O(1.2.4) multiple sequence alignment of bioactive peptide sequences (yellow) vs peptide sequences detected in the DIA secretome experiment at 72h. Alignment was performed online with UniProt (https://www.uniprot.org/align/).

**(A) C-terminal peptide from Secretogranin-5**

```
Reference SVPHFSDEDKDPE
Peptide 1 SVPHFSDEDK---
```

**(B) Manserin from Secretogranin-2**

```
Reference VPGQGSSEDDLQEEEQIEQAIKEHLNQGSSQETDKLAPVS
Peptide 1 VPGQGSSEDDLQEEEQIEQAIK------------------
Peptide 2 VPGQGSSEDDLQEEEQIEQAIKEHLNQGSSQETDK-----
Peptide 3 ----------------------EHLNQGSSQETDK-----
```

**(C) NEUG(55-78) from Neurogranin**

```
Reference GPGPGGPGGAGVARGGAGGGPSGD
Peptide 1 GPGPGGPGGAGVAR----------
```

**(D) Thymosin alpha 1 from Prothymosin alpha**

```
Reference SDAAVDTSSEITTKDLKEKKEVVEEAEN
Peptide 1 SDAAVDTSSEITTKDLK-----------
```

**(E) Thymopoietin from TMPO**

```
Reference PEFLEDPSVLTKDKLKSELVANNVTLPAGEQRKDVYVQLYLQHLTARNR
Peptide 1 ---------------------------------DVYVQLYLQHLTAR–
Peptide 2 ----------------SELVANNVTLPAGEQRK----------------
Peptide 3 ----------------SELVANNVTLPAGEQR-----------------
Peptide 4 PEFLEDPSVLTK-------------------------------------
```

**(F) CNP-53 from C-type natriuretic peptide**

```
Reference DLRVDTKSRAAWARLLQEHPNARKYKGANKKGLSKGCFGLKLDRIGSMSGLGC
Peptide 1 --------------LLQEHPNAR------------------------------
```

**(G) Preptin from Insulin-like growth factor 2**

```
Reference DVSTPPTVLPDNFPRYPVGKFFQYDTWKQSTQRL
Peptide 1 --------------------FFQYDTWK------
```

**(H) Short peptide from Alpha-1-antitrypsin**

```
Reference MFLEAIPMSIPPEVKFNKPFVFLMIEQNTKSPLFMGKVVNPTQK
Peptide 1 ------------------------------SPLFMGK-------
Peptide 2 ------------------PFVFLMIEQNTK-------------
```

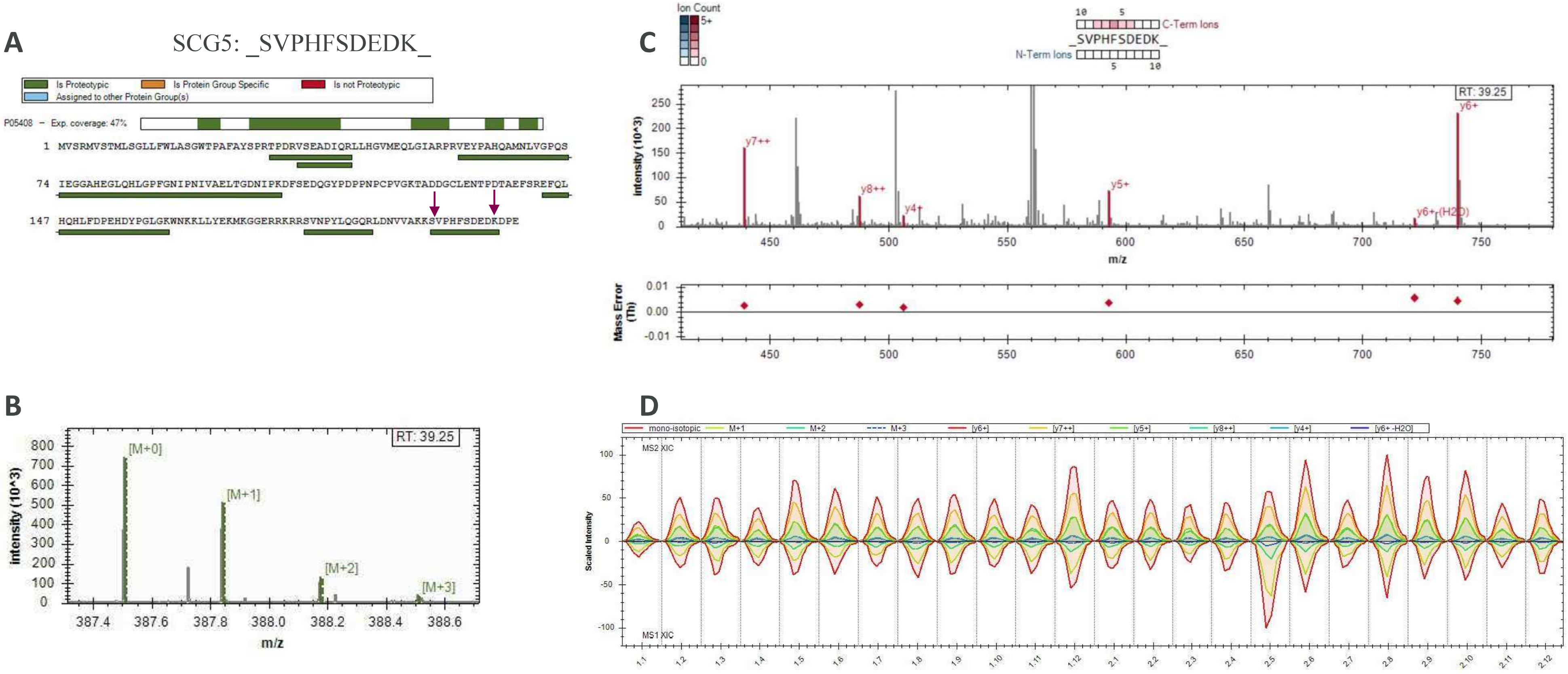


**Fig. S37. Proteotypic peptide corresponding to C-terminal peptide of SCG5 identified by DIA secretome proteomics experiment at 72h.** Total protein coverage (**A**), MS1 isotopic envelope (**B**), MS/MS fragmentation match and mass error (**C**), and XIC of MS1 & MS2 for the conditions (**D**) are shown for each given peptide identified.

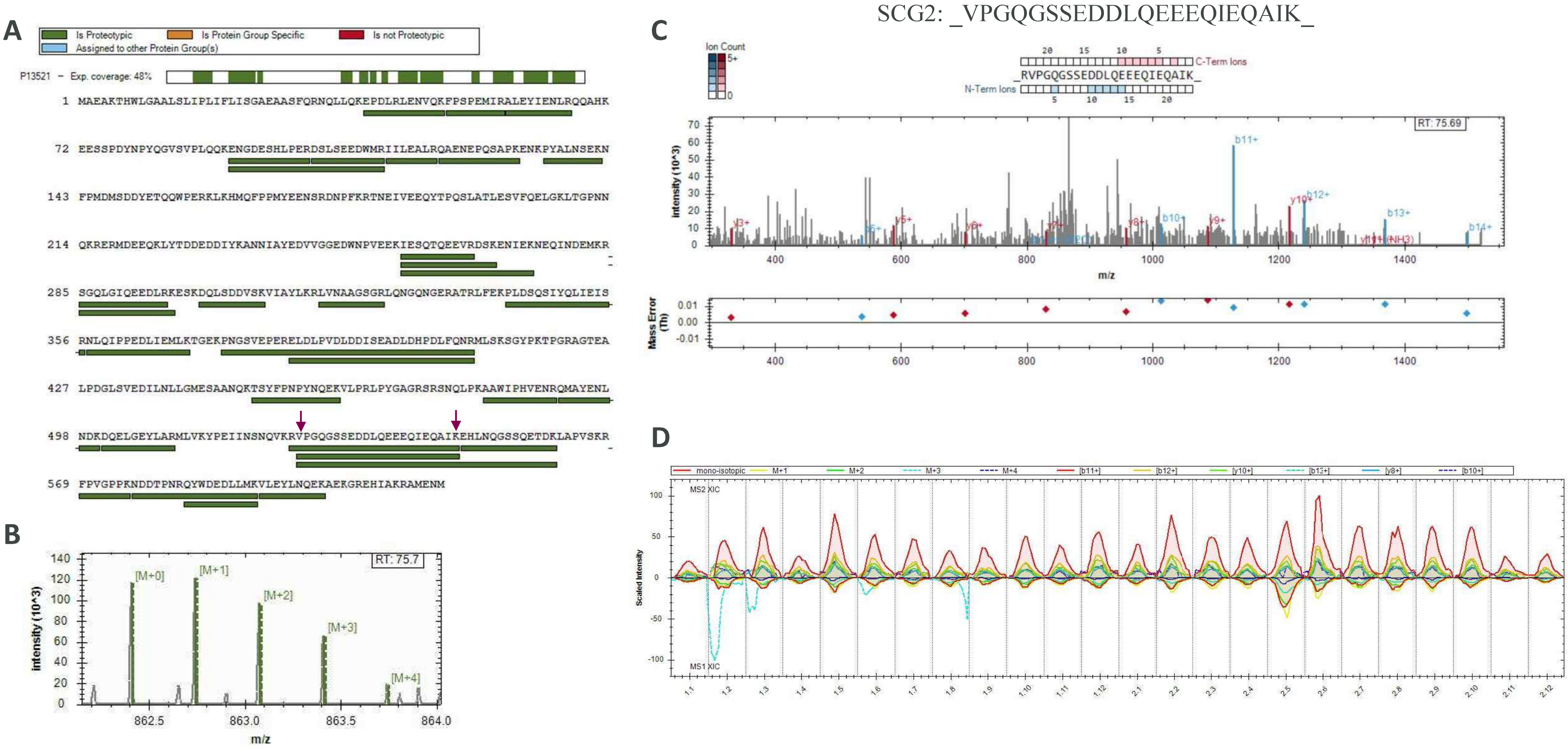


**Fig. S38. Proteotypic peptides corresponding to Manserin of SCG2 identified by DIA secretome proteomics experiment at 72h.** Total protein coverage (**A**), MS1 isotopic envelope (**B**), MS/MS fragmentation match and mass error (**C**), and XIC of MS1 & MS2 for the conditions (**D**) are shown for each given peptide identified.

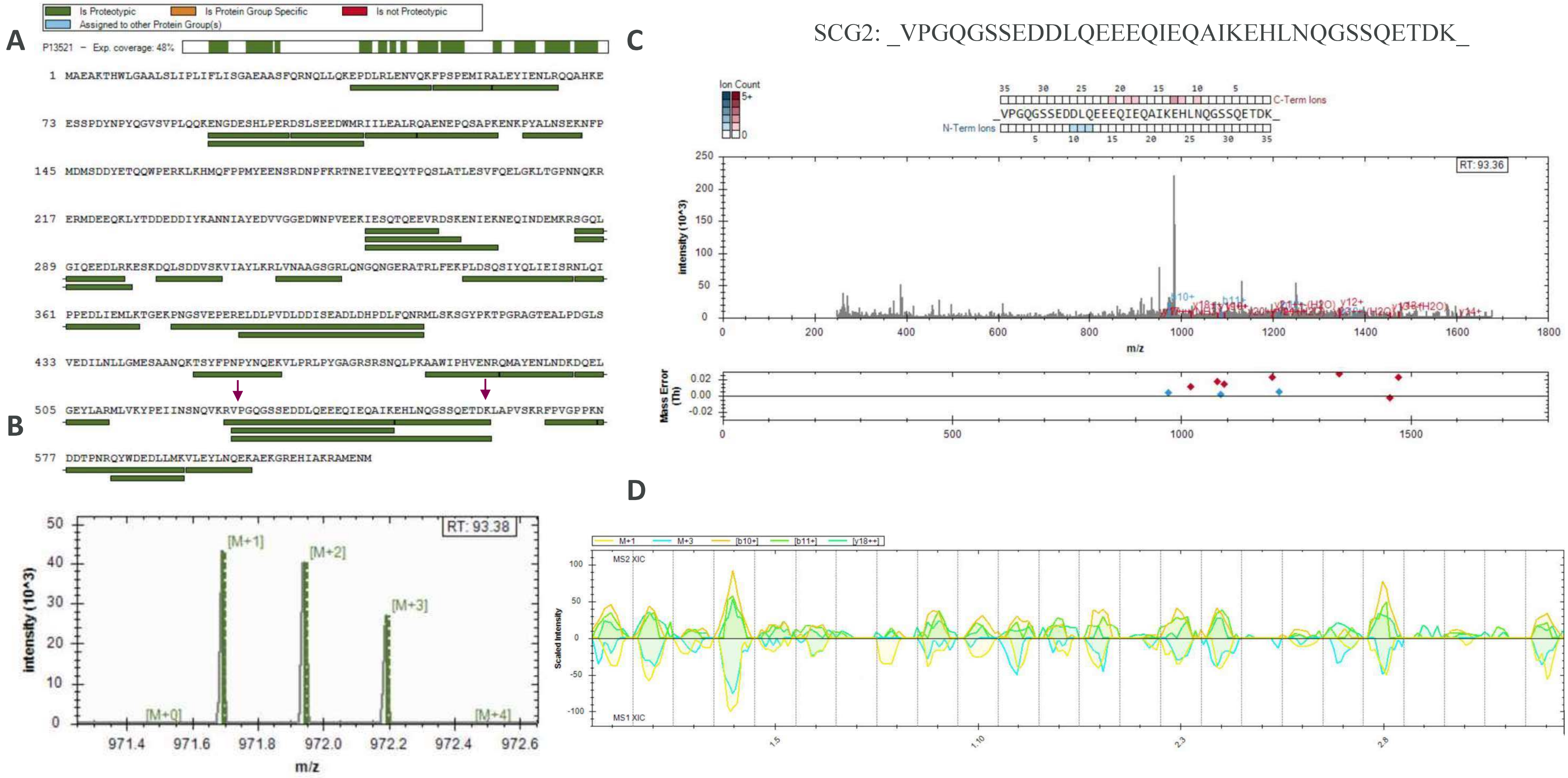


**Fig. S39. Proteotypic peptides corresponding to Manserin of SCG2 identified by DIA secretome proteomics experiment at 72h.** Total protein coverage (**A**), MS1 isotopic envelope (**B**), MS/MS fragmentation match and mass error (**C**), and XIC of MS1 & MS2 for the conditions (**D**) are shown for each given peptide identified.

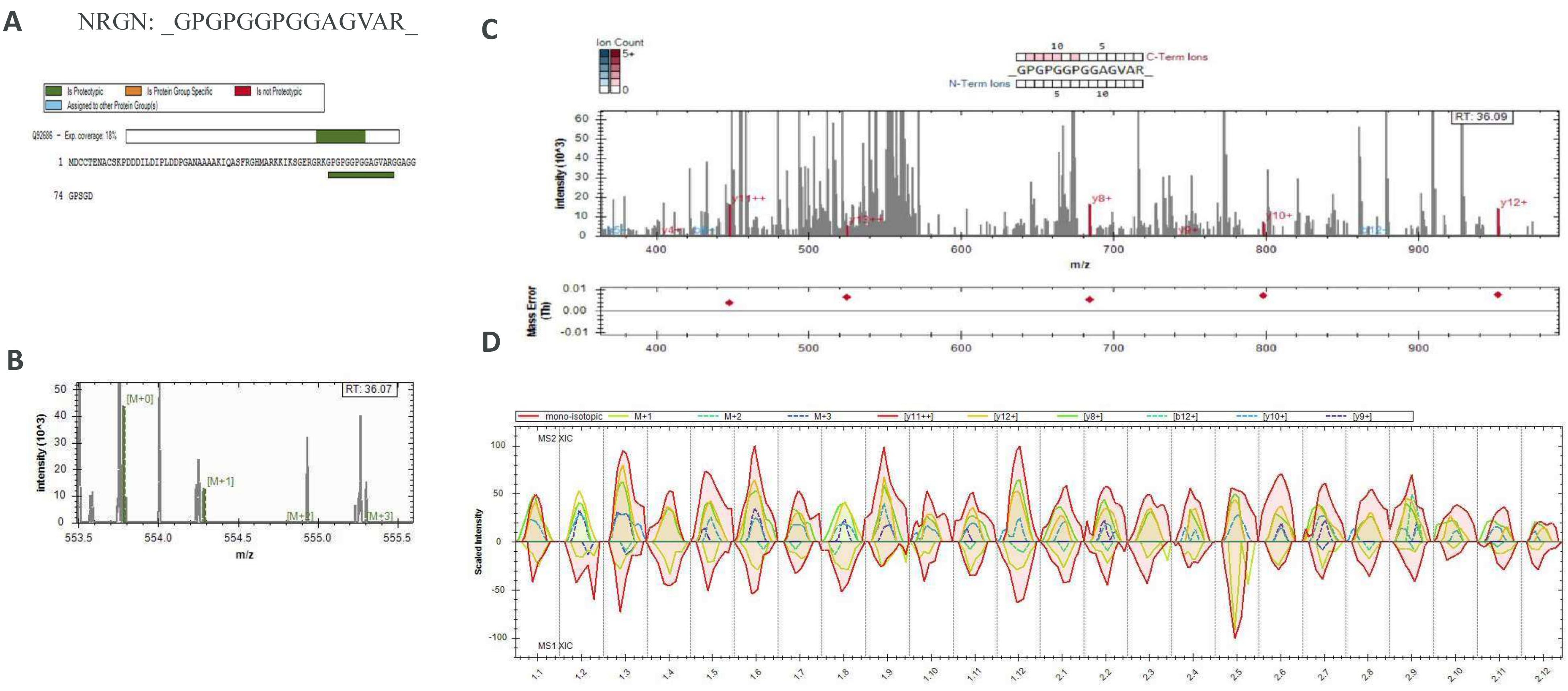


**Fig. S40. Proteotypic peptide corresponding to NEUG(55-78) from Neurogranin identified by DIA secretome proteomics experiment at 72h.** Total protein coverage (**A**), MS1 isotopic envelope (**B**), MS/MS fragmentation match and mass error (**C**), and XIC of MS1 & MS2 for the conditions (**D**) are shown for each given peptide identified.

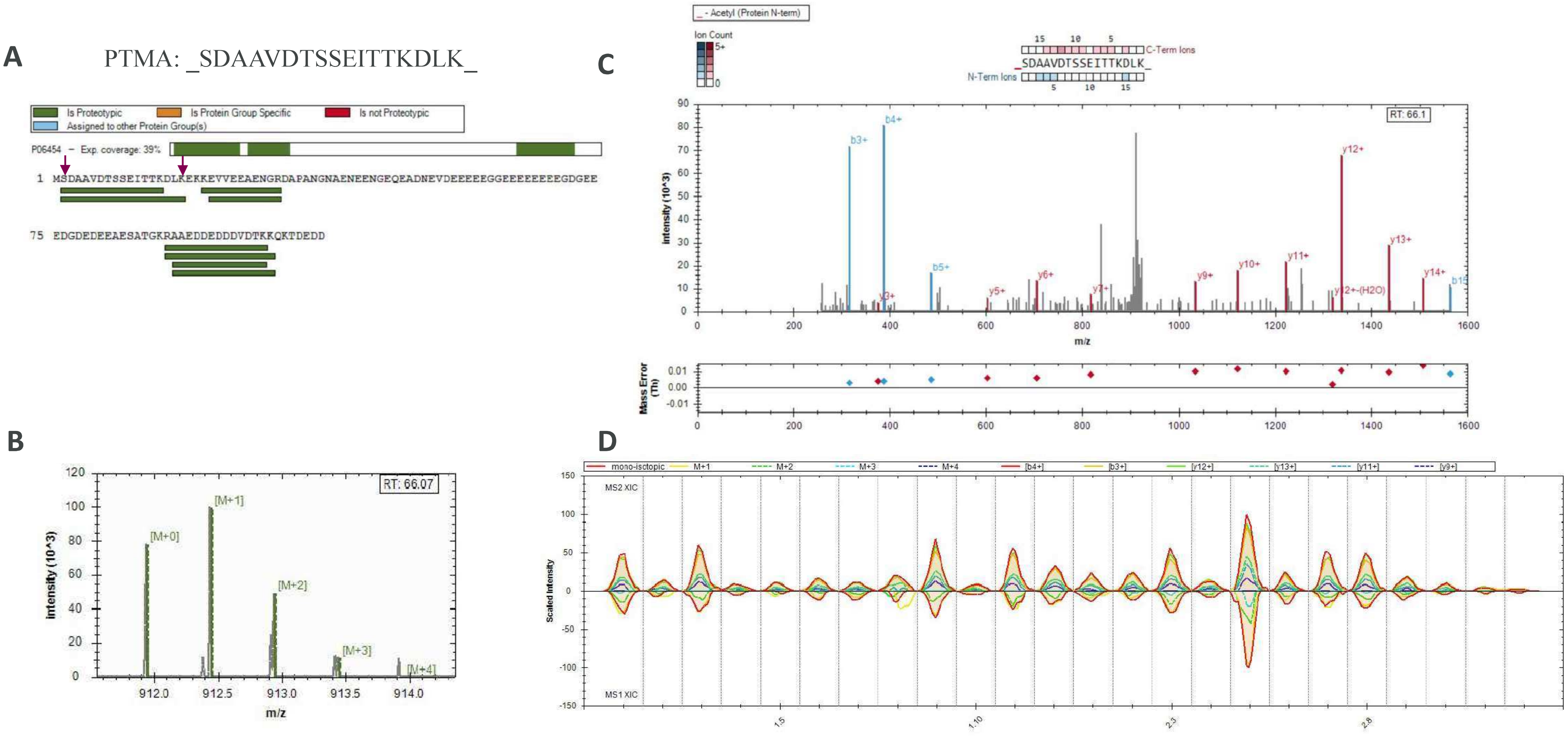


**Fig. S41. Proteotypic peptide corresponding to Thymosin alpha 1 identified by DIA secretome proteomics experiment at 72h.** Total protein coverage (**A**), MS1 isotopic envelope (**B**), MS/MS fragmentation match and mass error (**C**), and XIC of MS1 & MS2 for the conditions (**D**) are shown for each given peptide identified.

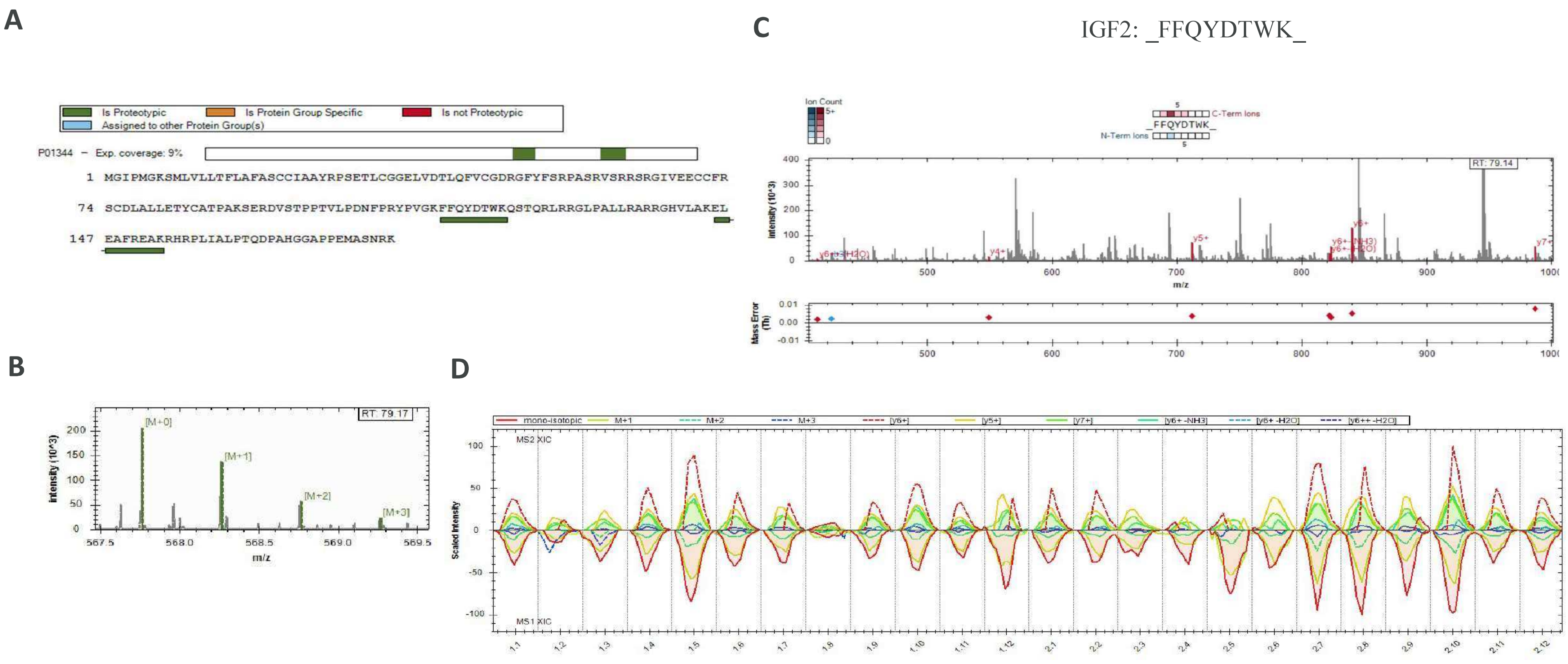


**Fig. S42. Proteotypic peptide corresponding to preptin in DIA secretome proteomics experiment at 72h.** Total protein coverage (**A**), MS1 isotopic envelope (**B**), MS/MS fragmentation match and mass error (**C**), and XIC of MS1 & MS2 for the conditions (**D**) are shown for each given peptide identified.

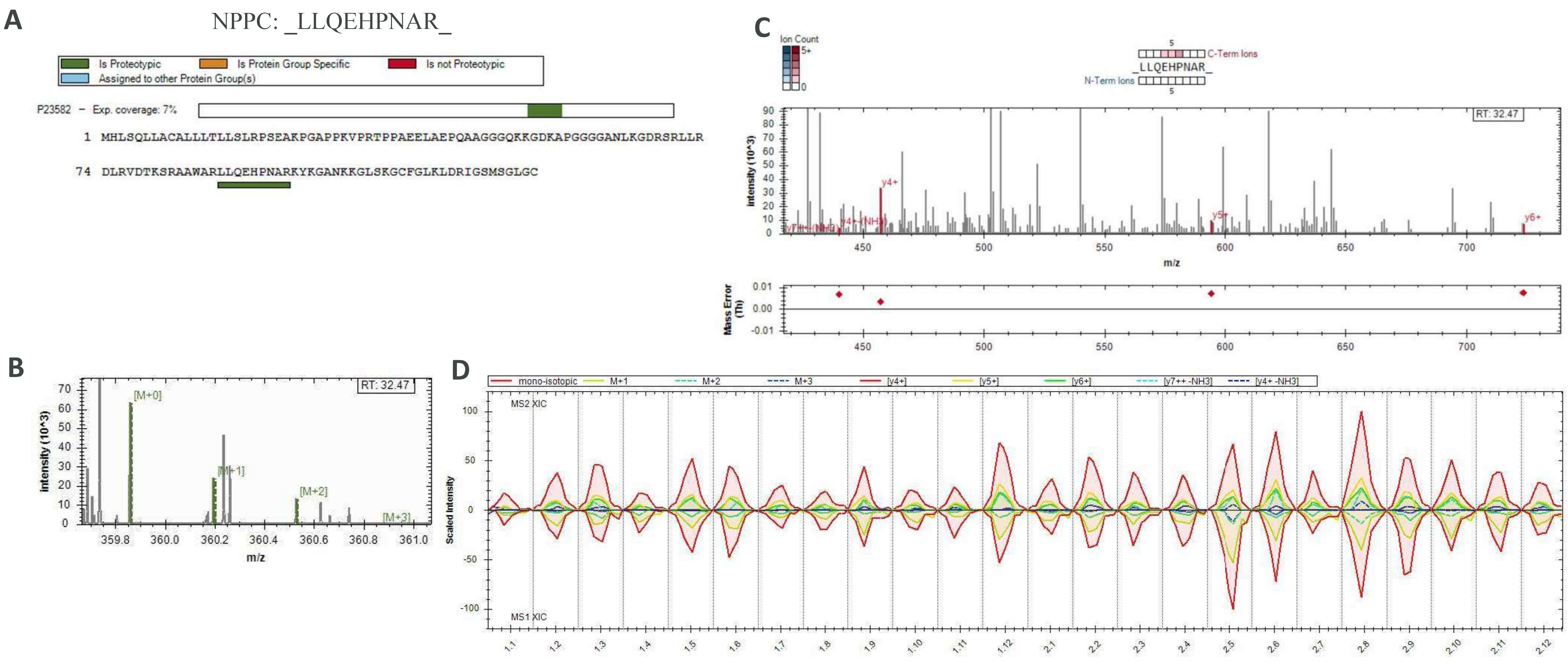


**Fig. S43. Proteotypic peptide corresponding to CNP-53 from C-type natriuretic peptide in DIA secretome proteomics experiment at 72h.** Total protein coverage (**A**), MS1 isotopic envelope (**B**), MS/MS fragmentation match and mass error (**C**), and XIC of MS1 & MS2 for the conditions (**D**) are shown for each given peptide identified.

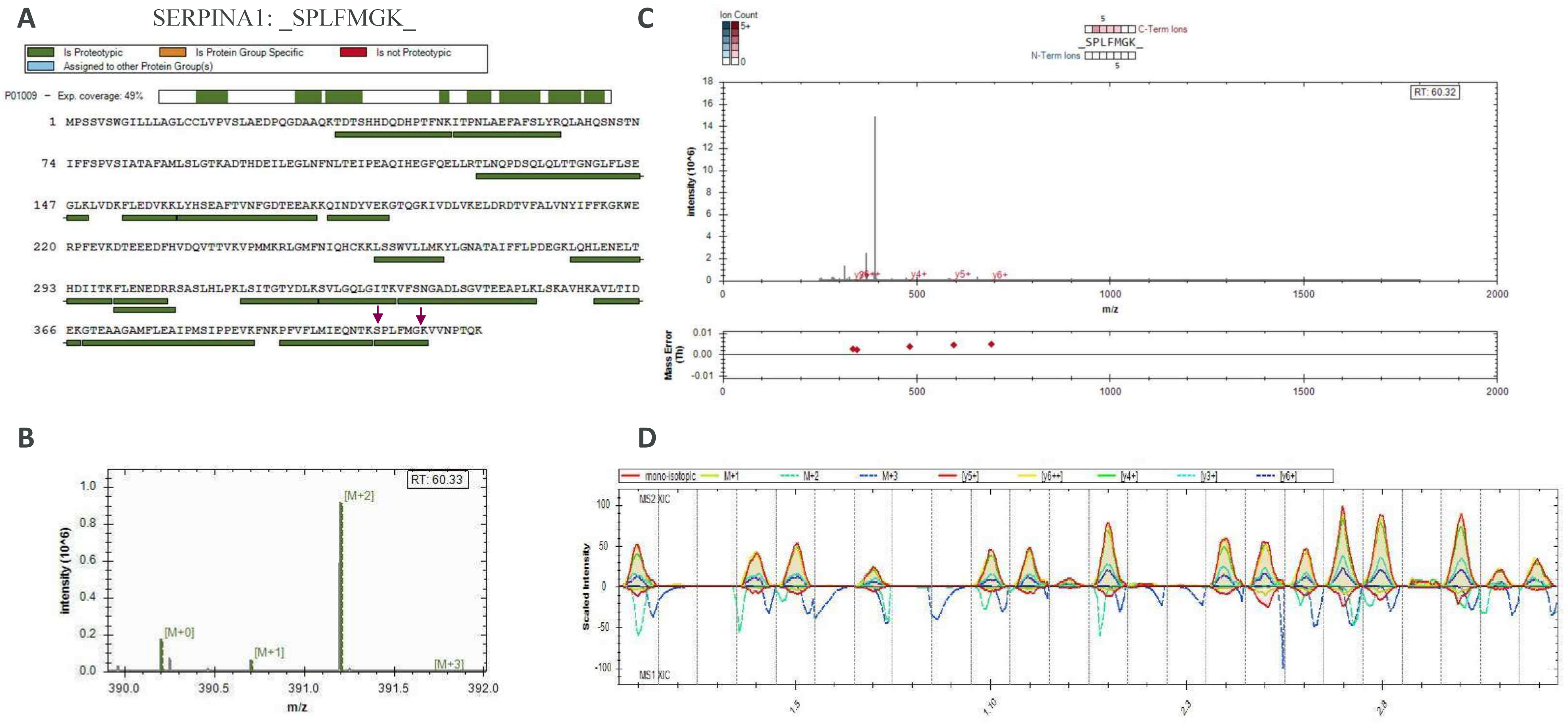


**Fig. S44. Proteotypic peptides corresponding to SERPINA1 short peptide in DIA secretome proteomics experiment at 72h.** Total protein coverage (**A**), MS1 isotopic envelope (**B**), MS/MS fragmentation match and mass error (**C**), and XIC of MS1 & MS2 for the conditions (**D**) are shown for each given peptide identified.

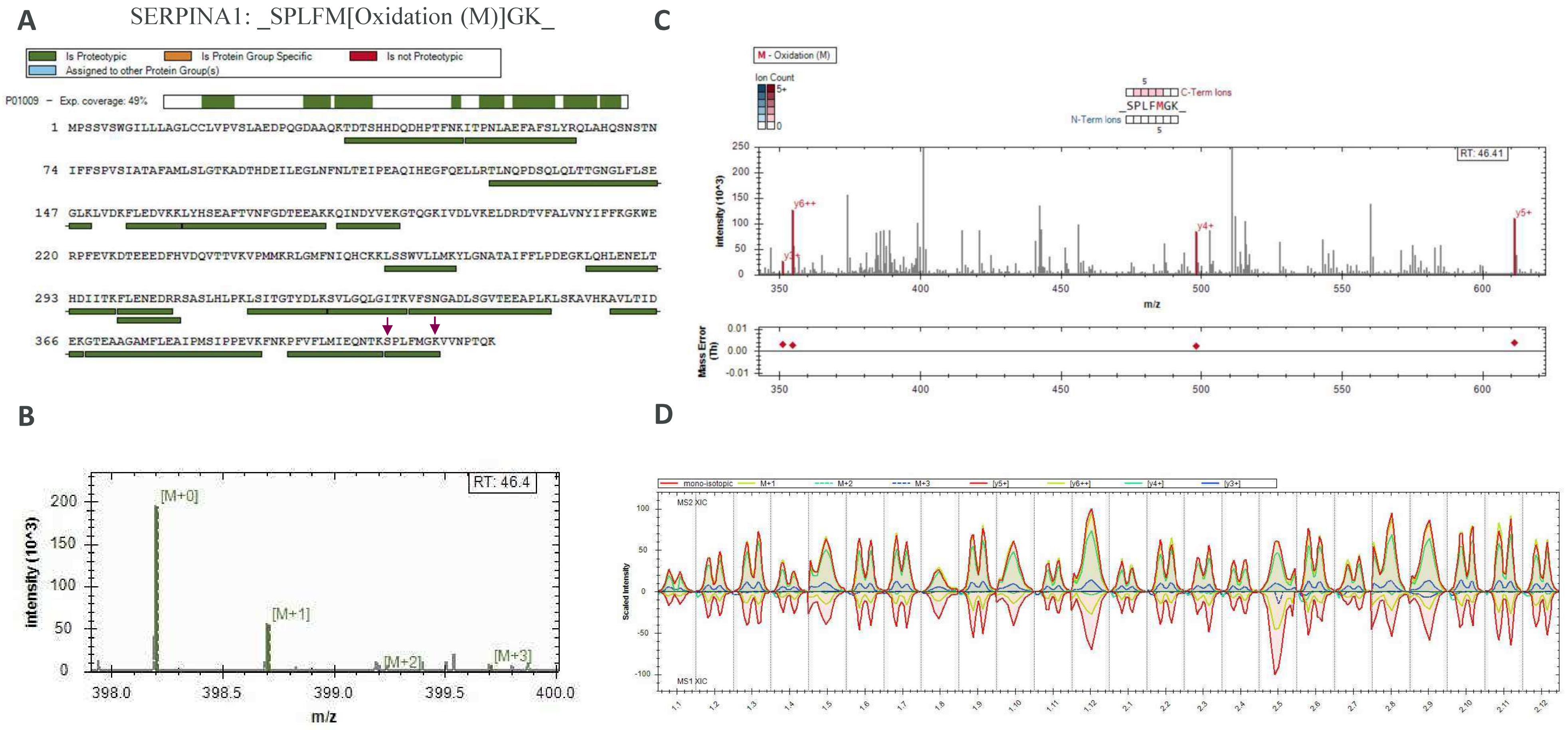


**Fig. S45. Proteotypic peptides corresponding to SERPINA1 short peptide in DIA secretome proteomics experiment at 72h.** Total protein coverage (**A**), MS1 isotopic envelope (**B**), MS/MS fragmentation match and mass error (**C**), and XIC of MS1 & MS2 for the conditions (**D**) are shown for each given peptide identified.

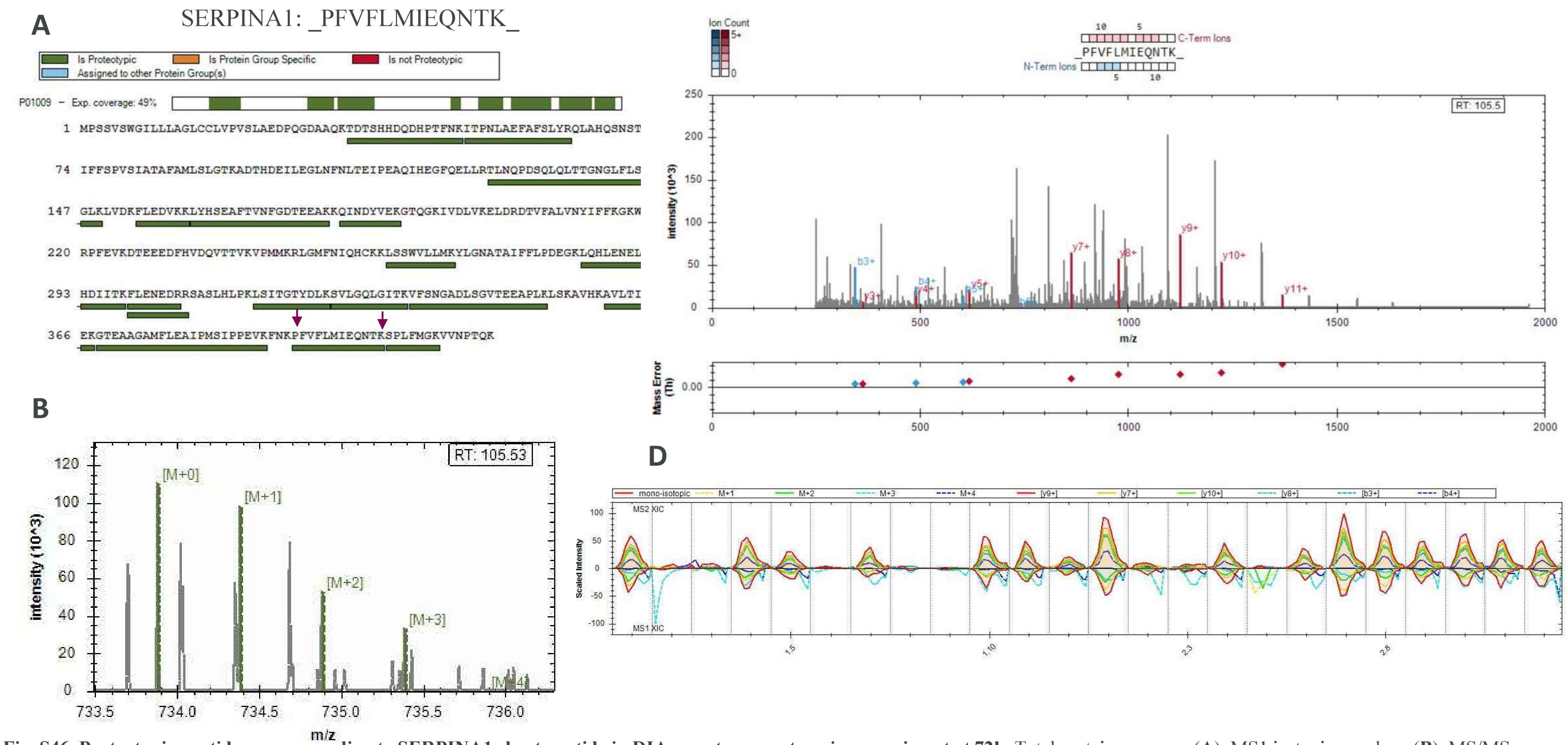


**Fig. S46. Proteotypic peptides corresponding to SERPINA1 short peptide in DIA secretome proteomics experiment at 72h.** Total protein coverage (**A**), MS1 isotopic envelope (**B**), MS/MS fragmentation match and mass error (**C**), and XIC of MS1 & MS2 for the conditions (**D**) are shown for each given peptide identified.

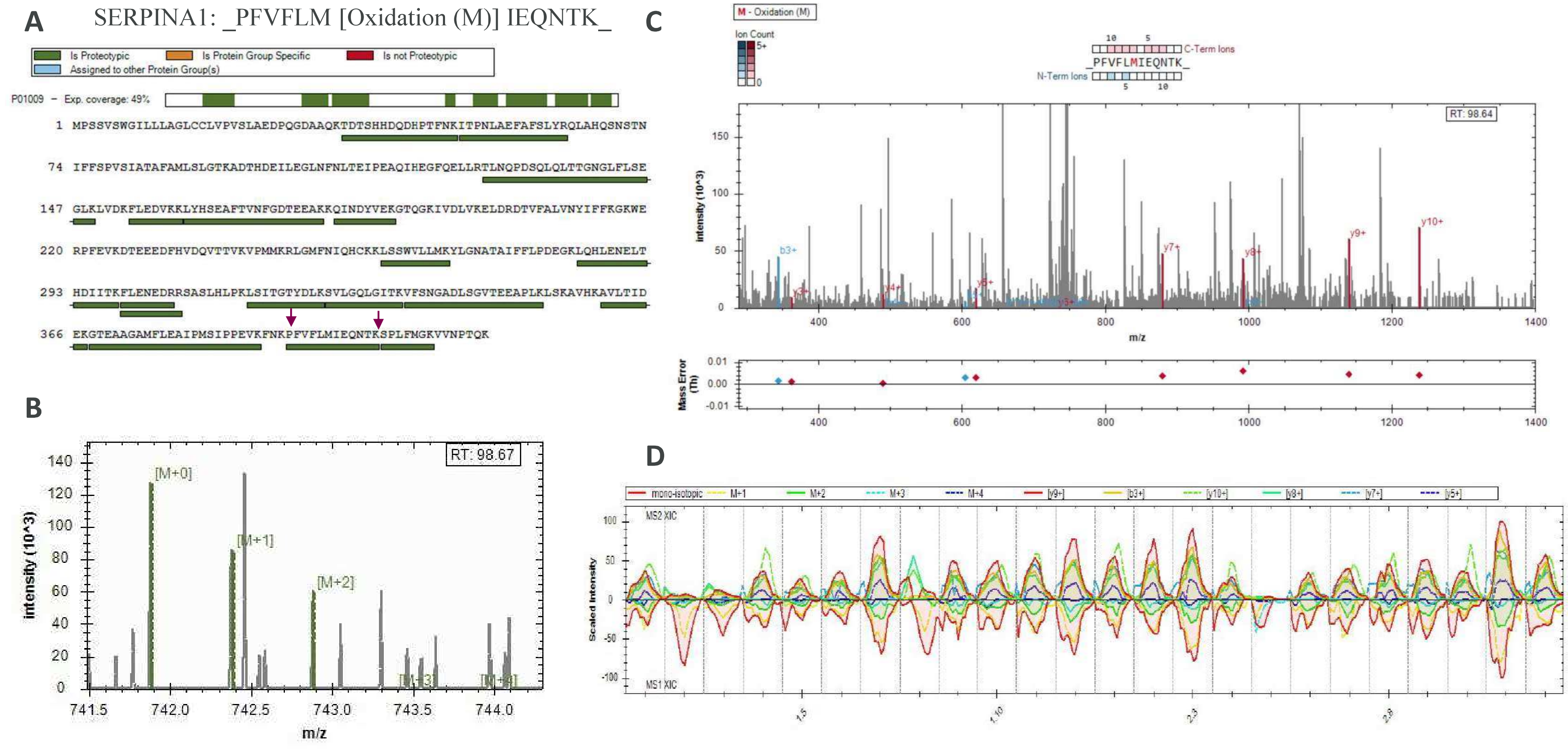


**Fig. S47. Proteotypic peptides corresponding to SERPINA1 short peptide in DIA secretome proteomics experiment at 72h.** Total protein coverage (**A**), MS1 isotopic envelope (**B**), MS/MS fragmentation match and mass error (**C**), and XIC of MS1 & MS2 for the conditions (**D**) are shown for each given peptide identified.

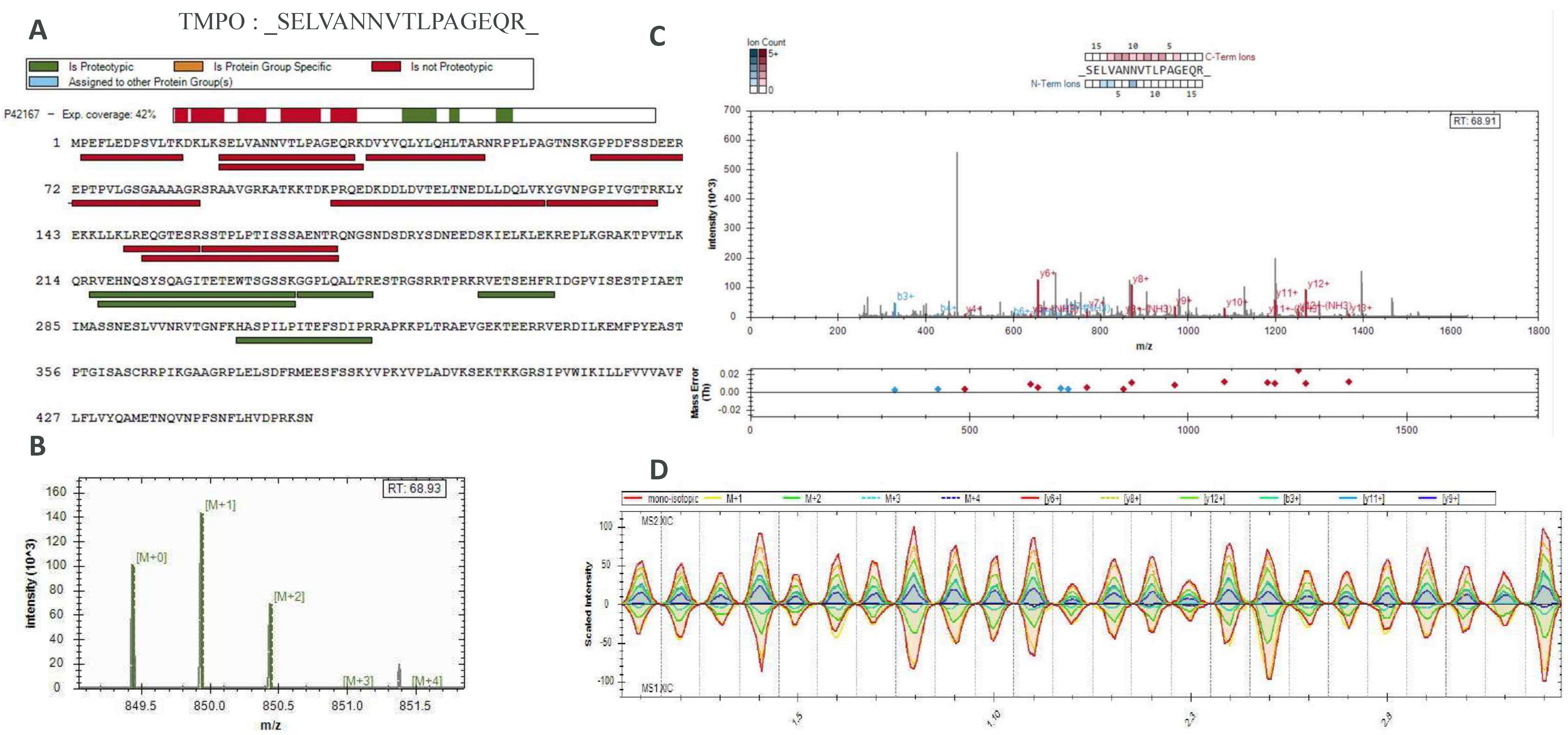


**Fig. S48. Proteotypic peptides corresponding to Thymopoietin in DIA secretome proteomics experiment at 72h.** Total protein coverage (**A**), MS1 isotopic envelope (**B**), MS/MS fragmentation match and mass error (**C**), and XIC of MS1 & MS2 for the conditions (**D**) are shown for each given peptide identified.

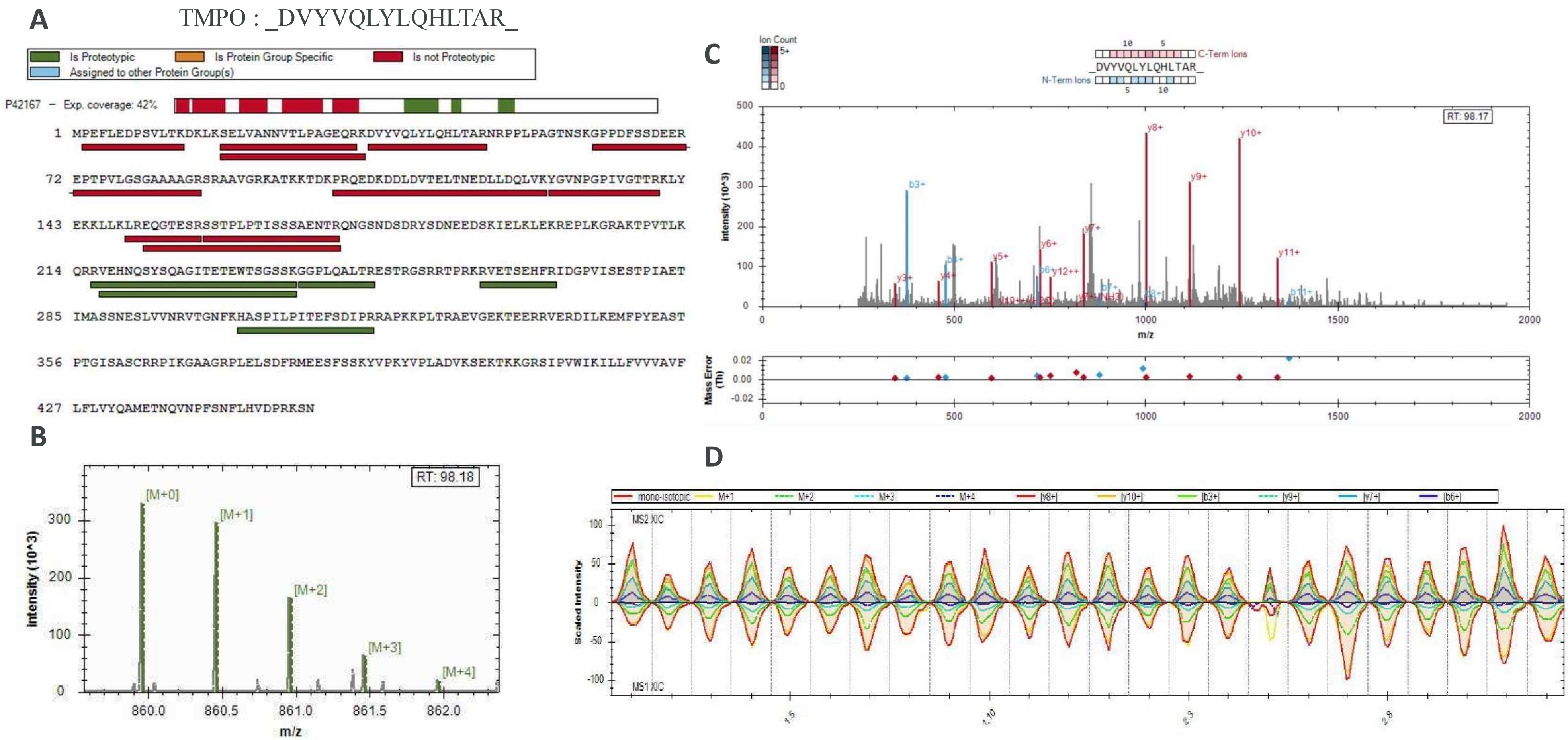


**Fig. S49. Proteotypic peptides corresponding to Thymopoietin in DIA secretome proteomics experiment at 72h.** Total protein coverage (**A**), MS1 isotopic envelope (**B**), MS/MS fragmentation match and mass error (**C**), and XIC of MS1 & MS2 for the conditions (**D**) are shown for each given peptide identified.

TMPO : _PEFLEDPSVLTK_

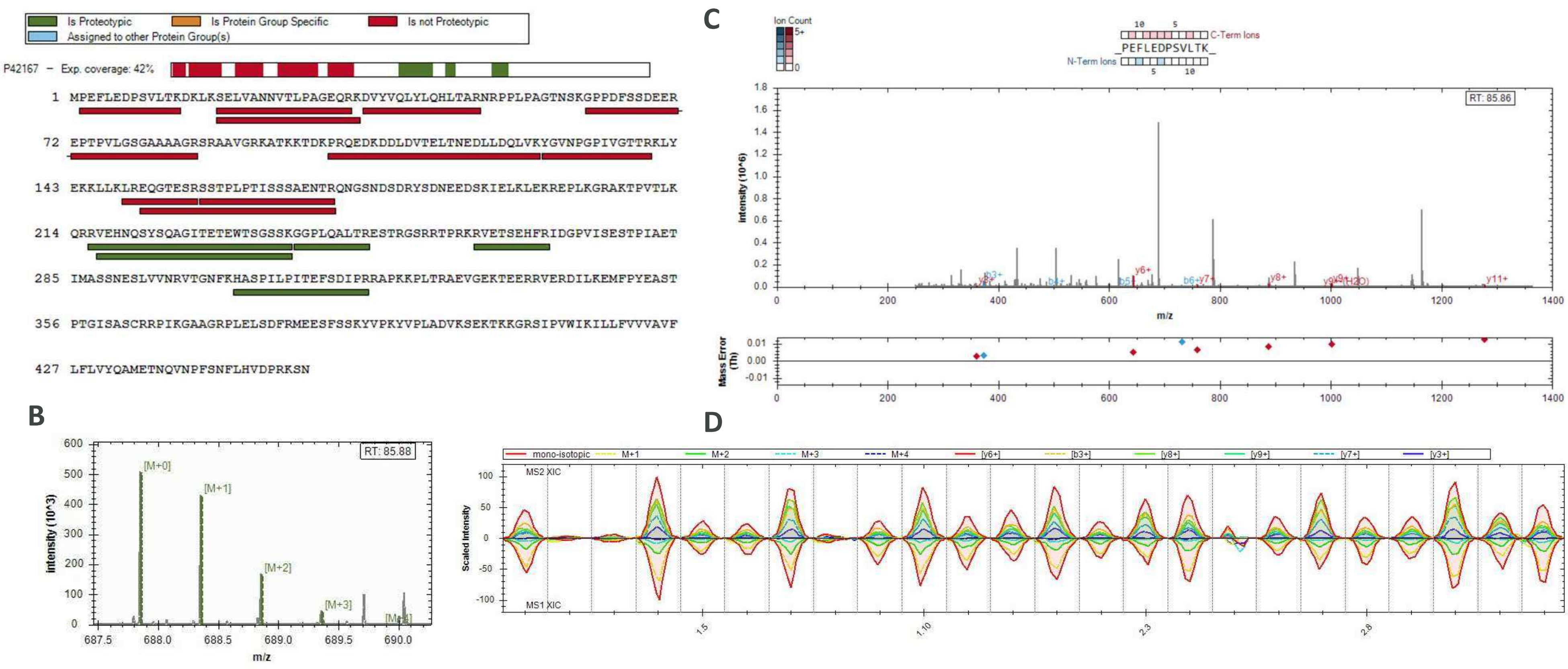


**Fig. S50. Proteotypic peptides corresponding to Thymopoietin in DIA secretome proteomics experiment at 72h.** Total protein coverage (**A**), MS1 isotopic envelope (**B**), MS/MS fragmentation match and mass error (**C**), and XIC of MS1 & MS2 for the conditions (**D**) are shown for each given peptide identified.

**Table S2. Top 150 secreted proteins in untreated EndoC-βH1 cells.** Proteins are ordered by decreasing median normalized expression abundance. N=3.

| rank | UniProt | Symbol | rank | UniProt | Symbol | rank | UniProt | Symbol |
|---|---|---|---|---|---|---|---|---|
| 1 | TF | P02787 | 51 | CUTA | O60888 | 101 | PRMT1 | Q99873 |
| 2 | ALB | P02768 | 52 | HADH | Q16836 | 102 | YWHAG | P61981 |
| 3 | CCDC197 | Q8NCU1 | 53 | TUBB2B | Q9BVA1 | 103 | NAXE | Q8NCW5 |
| 4 | KRT2 | P35908 | 54 | PDIA3 | P30101 | 104 | TMSB4X | P62328 |
| 5 | CHGA | P10645 | 55 | UBE2V1 | Q13404 | 105 | GAPDH | P04406 |
| 6 | HPX | P02790 | 56 | GPI | P06744 | 106 | RAN | P62826 |
| 7 | KRT1 | P04264 | 57 | FKBP2 | P26885 | 107 | TALDO1 | P37837 |
| 8 | KRT9 | P35527 | 58 | EPS15L1 | Q9UBC2 | 108 | RPS14 | P62263 |
| 9 | CCDC146 | Q8IYE0 | 59 | YWHAB | P31946 | 109 | UBE2V2 | Q15819 |
| 10 | MACF1 | Q9UPN3 | 60 | CFL1 | P23528 | 110 | ANXA2 | P07355 |
| 11 | USP47 | Q96K76 | 61 | RCN1 | Q15293 | 111 | QDPR | P09417 |
| 12 | VIM | P08670 | 62 | HSPE1 | P61604 | 112 | ALDOA | P04075 |
| 13 | KRT14 | P02533 | 63 | TPM4 | P67936 | 113 | GSS | P48637 |
| 14 | ACTB | P60709 | 64 | FN1 | P02751 | 114 | TXN | P10599 |
| 15 | PPIA | P62937 | 65 | HNRNPA2B1 | P22626 | 115 | PFN1 | P07737 |
| 16 | KRT10 | P13645 | 66 | PRDX1 | Q06830 | 116 | ZC4H2 | Q9NQZ6 |
| 17 | IGHG1 | P01857 | 67 | KRT19 | P08727 | 117 | LMNB1 | P20700 |
| 18 | IGHG3 | P01860 | 68 | TKT | P29401 | 118 | CALR | P27797 |
| 19 | IGHG4 | P01861 | 69 | ENO3 | P13929 | 119 | LDHB | P07195 |
| 20 | ZC3H4 | Q9UPT8 | 70 | PRDX3 | P30048 | 120 | GGH | Q92820 |
| 21 | IGKC | P01834 | 71 | PRSS3 | P35030 | 121 | PFN2 | P35080 |
| 22 | KRT18 | P05783 | 72 | PARK7 | Q99497 | 122 | SERPINH1 | P50454 |
| 23 | KRT6C | P48668 | 73 | P4HB | P07237 | 123 | ACTA2 | P62736 |
| 24 | H4C1 | P62805 | 74 | NUTF2 | P61970 | 124 | PGK1 | P00558 |
| 25 | ITIH4 | Q14624 | 75 | ACLY | P53396 | 125 | KTN1 | Q86UP2 |
| 26 | TPI1 | P60174 | 76 | HSPA8 | P11142 | 126 | SUMO3 | P55854 |
| 27 | LSM2 | Q9Y333 | 77 | FKBP10 | Q96AY3 | 127 | MANF | P55145 |
| 28 | IGHG2 | P01859 | 78 | ERP29 | P30040 | 128 | LDHA | P00338 |
| 29 | DBI | P07108 | 79 | UCHL1 | P09936 | 129 | GSR | P00390 |
| 30 | PLOD2 | O00469 | 80 | HSPA5 | P11021 | 130 | HSPA2 | P54652 |
| 31 | TTR | P02766 | 81 | NME2 | P22392 | 131 | TMSB10 | P63313 |
| 32 | SOD1 | P00441 | 82 | DDT | P30046 | 132 | LGALS1 | P09382 |
| 33 | INS | P01308 | 83 | MACROH2A1 | O75367 | 133 | DLD | P09622 |
| 34 | ENO1 | P06733 | 84 | RAP1GAP2 | Q684P5 | 134 | B2M | P61769 |
| 35 | YWHAE | P62258 | 85 | CSTB | P04080 | 135 | APOA2 | P02652 |
| 36 | MIF | P14174 | 86 | EIF4EBP1 | Q13541 | 136 | APEX1 | P27695 |
| 37 | YWHAZ | P63104 | 87 | TUBB4B | P68371 | 137 | LAMC1 | P11047 |
| 38 | MAN2B2 | Q9Y2E5 | 88 | ALDH1A1 | P00352 | 138 | SAG | P10523 |
| 39 | CPE | P16870 | 89 | GDI2 | P50395 | 139 | HPRT1 | P00492 |
| 40 | PPIB | P23284 | 90 | HSP90B1 | P14625 | 140 | SERPINC1 | P01008 |
| 41 | DCD | P81605 | 91 | GOT1 | P17174 | 141 | SLC3A2 | P08195 |
| 42 | IGHA1 | P01876 | 92 | SYMPK | Q92797 | 142 | HNRNPD | Q14103 |
| 43 | KRT8 | P05787 | 93 | TXNL1 | O43396 | 143 | MCFD2 | Q8NI22 |
| 44 | KRT5 | P13647 | 94 | MYDGF | Q969H8 | 144 | RPS3A | P61247 |
| 45 | PCBD1 | P61457 | 95 | GANAB | Q14697 | 145 | ANXA5 | P08758 |
| 46 | PGAM1 | P18669 | 96 | HNRNPA1 | P09651 | 146 | FKBP1A | P62942 |
| 47 | MDH1 | P40925 | 97 | GLO1 | Q04760 | 147 | BLMH | Q13867 |
| 48 | LTF | P02788 | 98 | IGKV4-1 | P06312 | 148 | S100A4 | P26447 |
| 49 | MDH2 | P40926 | 99 | NPC2 | P61916 | 149 | LAMB1 | P07942 |
| 50 | HDLBP | Q00341 | 100 | DNAJB11 | Q9UBS4 | 150 | NID1 | P14543 |
